# T-World: A highly general computational model of a human ventricular myocyte

**DOI:** 10.1101/2025.03.24.645031

**Authors:** Jakub Tomek, Xin Zhou, Hector Martinez-Navarro, Maxx Holmes, Thomas Bury, Lucas Arantes Berg, Marketa Tomkova, Emily Jo, Norbert Nagy, Ambre Bertrand, Alfonso Bueno-Orovio, Michael Colman, Blanca Rodriguez, Donald Bers, Jordi Heijman

**Affiliations:** Department of Anatomy, Physiology and Genetics (University of Oxford); Department of Pharmacology (UC Davis); Department of Computer Science (University of Oxford); Department of Physiology (McGill University); Ludwig Cancer Research (University of Oxford); Department of Pharmacology and Pharmacotherapy (University of Szeged); School of Biomedical Sciences (Leeds University); Medical Physics and Biophysics (Medical University of Graz)

## Abstract

Cardiovascular disease is the leading cause of death, demanding new tools to improve mechanistic understanding and overcome limitations of stem cell and animal-based research. We introduce T-World, a highly general virtual model of human ventricular cardiomyocyte suitable for multiscale studies. T-World shows comprehensive agreement with human physiology, from electrical activation to contraction, and is the first to replicate all key cellular mechanisms driving life-threatening arrhythmias. Extensively validated on unseen data, it demonstrates strong predictivity across applications and scales. Using T-World we revealed a likely sex-specific arrhythmia risk in females related to restitution properties, identified arrhythmia drivers in type 2 diabetes, and describe unexpected pro-arrhythmic role of NaV1.8 in heart failure. T-World demonstrates strong performance in predicting drug-induced arrhythmia risk and opens new opportunities for predicting and explaining drug efficacy, demonstrated by unpicking effects of mexiletine in Long QT syndrome 2. T-World is available as open-source code and an online app.

## Introduction

Computational modelling and simulations of cardiac cellular and organ physiology have become an integral part of contemporary cardiovascular research, providing insights into basic physiological mechanisms^1^, mechanisms of arrhythmia^1^, therapy guidance^2^, and drug safety assessment^3,4^. Digital evidence increasingly influences real-world applications in the pharmaceutical industry^5^ and regulatory bodies such as the FDA^6^ and EMA^7^. Combined with recent advances in data availability, hardware, and software, these models are driving the vision of the digital twin technology ^8^, producing a virtual tool that integrates clinical data acquired for an individual and that enables personalised diagnosis and treatment strategies.

Computational modelling and simulations also hold tremendous potential to contribute to reducing, replacing, and refining the use of animals in research (‘3R principles’). While some animal-based studies in cardiac research are indispensable, simulations can minimise animal use by guiding experimental design, predicting outcomes, and aiding interpretation. Human-specific virtual cells can also predict functional implications of animal data in the context of human physiology, addressing critical species differences that may, e.g., make a drug safe in mice but dangerous in humans^9^. Correspondingly, the European Medicines Agency has recognised computational modelling as a key trend in advancing 3R principles^10^.

Multiple successful models of human ventricular cardiomyocytes have been developed to investigate specific mechanisms of cardiac (patho)physiology and arrhythmia. Rudy-family models (ORd and ToR-ORd) excel at predicting drug responses and generating early afterdepolarisations in realistic conditions, making them valuable for drug studies comprising safety and efficacy assessment ^3,4^. Bers/Grandi-family models are known for realistic calcium handling^11–13^. The Ten Tusscher 2006 (TP06) model is widely used to study arrhythmia related to restitution properties^14^. Despite their strengths, each model family lacks generality, capturing only a small subset of arrhythmic behaviours and manifesting important discrepancies with experimental data on fundamental physiology. This limits their utility for mechanistic studies, analysing multifactorial drug effects, modelling complex diseases such as Type-2 diabetes (T2D) and heart failure, or integrative arrhythmia studies. Cells and their models are highly complex and include numerous components connected through non-linear feedback loops. As a result, flaws in one model component can cascade, leading to incorrect predictions in other components and behaviours. This limits a model’s predictive power and usefulness beyond its original focus. At the same time, the most innovative and relevant applications often arise precisely in these out-of-domain contexts. The lack of generality is in part also why different cellular models have typically been used to study aspects of arrhythmogenesis at cellular versus organ level^1^.

The absence of a comprehensive and physiologically accurate virtual cardiomyocyte impedes progress toward translational applications and expanding the context of use of cardiac simulations. To bridge this gap and unlock the full potential of cardiac simulations in research, industry, and clinic, we sought to develop a unified highly general virtual cell model. The generality should include 1) accurate recapitulation of human cellular cardiac physiology and its modulation by drugs or physiological changes, 2) the capability to manifest all key arrhythmogenic behaviours in conditions used to provoke them experimentally. This comprises early and delayed afterdepolarisations (EADs and DADs)^15,16^, alternans^17^, and steep restitution of action potentials (APs)^18^, 3) show suitability for multiscale modelling, enabling organ-level simulations.

Here, we present T-World, a novel virtual human cardiomyocyte that reproduces for the first time all key cellular arrhythmic mechanisms and shows comprehensive agreement with data on human cardiac physiology. Comparison to data not used in model creation demonstrates robust predictive accuracy at a broad range of tasks. T-World integrates electrophysiology, calcium handling, cardiomyocyte contraction, sympathetic stimulation, and sex differences, enabling comprehensive studies of their interactions. T-World is freely available via Matlab, CellML, C, and CUDA, with a free-to-use online graphical interface for non-coders (also runnable in Python). This model advances our understanding of cardiac electrophysiology and arrhythmogenesis by 1) identifying a likely sex-specific arrhythmia risk in females linked to restitution properties, 2) showing applicability to analysis of drug efficacy through analysis of mechanism of action of anti-arrhythmic drugs, and showing excellent performance in drug safety testing and, 3) identifying causes of high arrhythmia risk in T2D, and 4) suggesting NaV1.8 as a relevant treatment target in heart failure.

## Results

### Representation and validation of cell and organ physiology

Based on extensive experimental data, the T-World model represents a broad range of ionic currents and fluxes across distinct cellular compartments, as well as subcellular signalling pathways and contractility (see **Figure 1A** for a high-level overview). Distinct model components are described by sets of ordinary differential equations, constructed to recapitulate baseline experimental data on single ionic currents and other cellular elements. Coupling all those components together yields a virtual cardiomyocyte with a high degree of biological detail and realism, which can be used as a model system in cardiac research.

**Figure 1.**
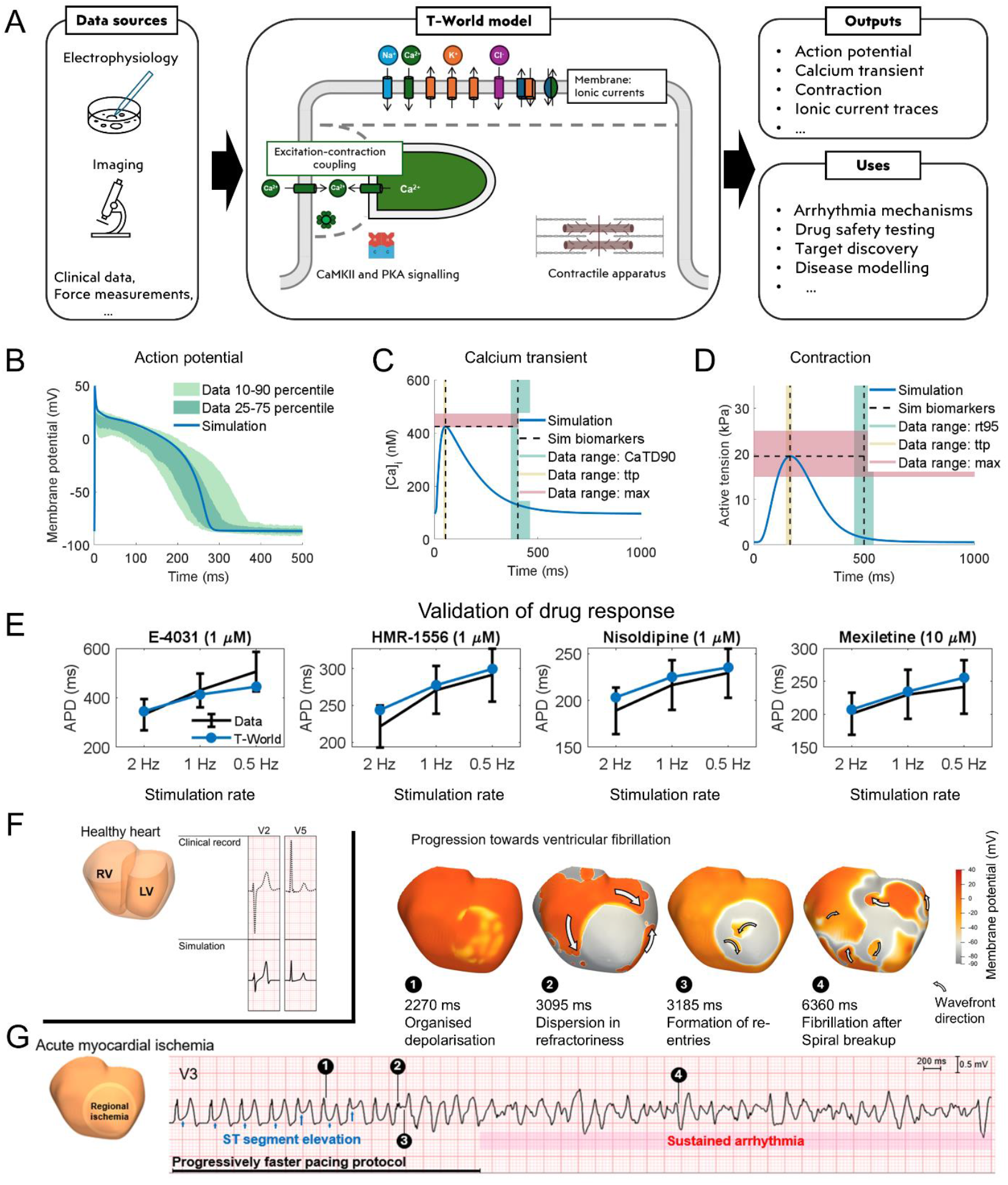
Cell and organ physiology. **A)** Conceptual diagram of the T-World model and its potential applications. See Methods for a detailed diagram including all ionic currents and cellular compartments. **B)** Endocardial action potential of T-World versus experimental ranges ^19^. Slightly higher peak in simulation versus data was chosen, given that our model is a single cell, whereas the experimental data are in small tissue samples which show a reduced peak due to cell-to-cell coupling. **C)** Calcium transient (CaT) of T-World with highlighted biomarkers versus experimental ranges for: CaT duration at 90% recovery level (CaTD90, shown in green), time to peak (ttp, shown in yellow), and calcium transient amplitude (max, shown in red), based on standard error of mean ranges data by Coppini et al.^20^ **D)** Active tension developed by T-World with highlighted biomarkers versus experimental ranges for: time from peak to 95% recovery (rt95, shown in green), time to peak (ttp, shown in yellow), and maximum active tension (max, shown in red), based on Margara et al.^21^. **E)** Independent validation of the APD prolongation or shortening induced by 1 µM E-4031 (70% I_Kr_ block), 1 µM HMR-1556 (90% I_Ks_ block), 1 µM nisoldipine (90% I_CaL_ block), and 10 µM mexiletine (54% I_NaL_, 9% I_Kr_, 20% I_CaL_ block) at 0.5, 1.0 and 2.0 Hz pacing in the four models. Drug concentrations and their effects on channel blocks are based on O’Hara et al._19_ Please note the distinct y-axes for the four drugs. **F)** Healthy ventricular model constructed from clinical MRI data and ECG simulation (solid line) compared with the ECG record from the patient used for the ventricular anatomy. **G)** Ventricular fibrillation simulation in the setting of acute ischemia, when stimulation rate is progressively increased. Heart snapshots above the ECG illustrate different stages of progression towards fibrillation.

The three key outputs of a cardiomyocyte model are its AP, calcium transient (CaT), and the resulting active tension during contraction. T-World shows a very strong agreement with human AP data^19^ with regards to AP duration (APD), resting membrane potential, and the overall shape of the AP during plateau and recovery (**Figure 1B**). It is in better agreement with human AP shape than most prior state-of-the-art models (**Supplementary Note 1**). The CaT is also in excellent agreement with human data on time to peak, duration, and amplitude^20^ (**Figure 1C**). T-World incorporates the Land model of contraction^22^ as in the work of Margara et al.^21^, and its outputs are fully consistent with experimental data on time to peak force, amplitude, and time to 95% recovery of contractility in human myocardium ^21^ (**Figure 1D**).

The cardiac AP is determined by the specific mixture of ionic currents, and a given AP shape can be achieved through various combinations and balances of currents^23^. To verify that the balance of key ionic currents in T-World is human-like, we validated it by simulating its exposure to four simulated channel-blocking drugs at three pacing rates (**Figure 1E-H**). The strong predictive performance predisposes T-World to applications in safety pharmacology. See **Supplementary note 2** for comparison to other models (noting, in particular, problematic performance of TP06).

Excitation-contraction coupling (ECC) in cardiomyocytes is a process that ensures that electrical signals translate into muscle contraction and pumping of the heart. It involves 1) electrical activation of the cell, 2) consequent opening of L-type calcium channels, 3) triggering intracellular calcium release from the sarcoplasmic reticulum (SR), 4) binding of the released calcium to the contractile apparatus and resulting physical contraction. To correctly represent ECC, we introduced numerous new developments in T-World compared to prior models, yielding a virtual cell that is in excellent agreement with available data, while being mechanistically realistic. See **Supplementary note 3** for details of ECC development and validation. Briefly, the model maintains the realism of calcium handling from Bers/Grandi formulations, while improving upon several important limitations of the framework. First, the model shows more physiological timing of calcium release in response to L-type calcium channel opening with implications for better AP shape and overall model plausibility. Second, T-World correctly responds to changes in SERCA pump function, which enables plausible representation of disease and sympathetic nervous activity. The model shows comprehensive agreement with heart-rate-dependence of various components of ECC, such as calcium transient amplitude, contraction force, or SR calcium content; something not achieved in prior models. Sodium concentration rate-dependence is also captured well, as is the relative contribution of SERCA, NCX, and sarcolemmal calcium pump to clearance of calcium from cytosol within an AP. Finally, T-World has human-like properties of the L-type calcium current (essential in ECC), including current-voltage relationship, recovery from refractoriness, and relative contribution of voltage- and calcium-dependent inactivation.

ECC and electrophysiology are strongly modulated by the β-adrenergic (βAR) signalling pathway, which mediates the myocardial response to sympathetic nervous stimulation. Our model includes the Heijman et al.^24^ βAR description, with modifications to account for updates to ionic currents and inclusion of the contractile apparatus in the model (see **Supplementary Methods**). The integrated model was calibrated based on human AP data, with subsequent validation demonstrating a correct effect on CaT and contraction dynamics (**Supplementary Note 4**).

Pronounced differences exist between hearts from females and males, which subsequently translate into differential risk of various adverse cardiac outcomes^25^. Given the extent and importance of sex differences in cardiovascular physiology, we constructed a male and a female version of T-World, based on available experimental data and prior simulation approaches^26–28^, showing correctly longer APD and slightly reduced CaT amplitude and contraction in female myocytes (**Supplementary Note 5**), supporting the utility of T-World for studies on sex differences in cardiac (patho)physiology. There are also T-World versions for endocardial, midmyocardial, and epicardial myocytes.

The virtual cell model can be used to build a virtual organ based on clinical MRI data, which enables organ-level studies and reconstruction of ECG. We assessed the model’s performance across scales by building a 3D model based on a patient’s anatomy, with the simulation yielding a human-like ECG signal (**Figure 1F**) ^29,30^. A major application of whole-organ models is the study of arrhythmia such as ventricular fibrillation (VF), where the impact of tissue-level phenomena such as fibrosis and conduction heterogeneities can be considered. However, numerous advanced models like ToR-ORd or ORd struggled to produce VF dynamics unless their parameters were specifically tuned for this purpose (in addition to imposing a pro-arrhythmic substrate such as localised ischemia)^30,31^. Importantly, T-World does reproduce VF, as shown in **Figure 1G**, where VF appears in the setting of acute anteroseptal ischemia and progressively increasing rate of stimulation. Initially, the electrical propagation is stable, only manifesting ST segment elevation (a hallmark of acute ischemia), but as the stimulation rate is increased, re-entrant wavefronts appear (**Figure 1G**, snapshots 2,3), and gradually progress to spiral wave breakup and VF (snapshot 4).

### Cellular arrhythmic behaviours in T-World

#### Early afterdepolarisations

EADs, extrasystolic depolarisations during an AP, contribute to arrhythmogenesis and are commonly linked to drug-induced cardiotoxicity and long QT syndromes, being typically driven by the reactivation of I_Ca,L_ during prolonged APD^15^ (**Figure 2A**). T-World replicates EADs under realistic conditions of drug-induced long QT (**Figure 2B**), similar to ToR-ORd and ORd models ^4,19^, with a 13-mV amplitude, similar to experimental observations ^32^. In contrast, the TP06 model requires nearly tripled I_CaL_ to manifest EADs ^34^, likely due to excessive I_Ks_ providing strong repolarisation reserve. The Morotti2021 model^13^ (the most recent human model from the Bers/Grandi family) similarly necessitates a +150% I_CaL_ increase to induce EADs (**Supplementary Figure S23**).

**Figure 2.**
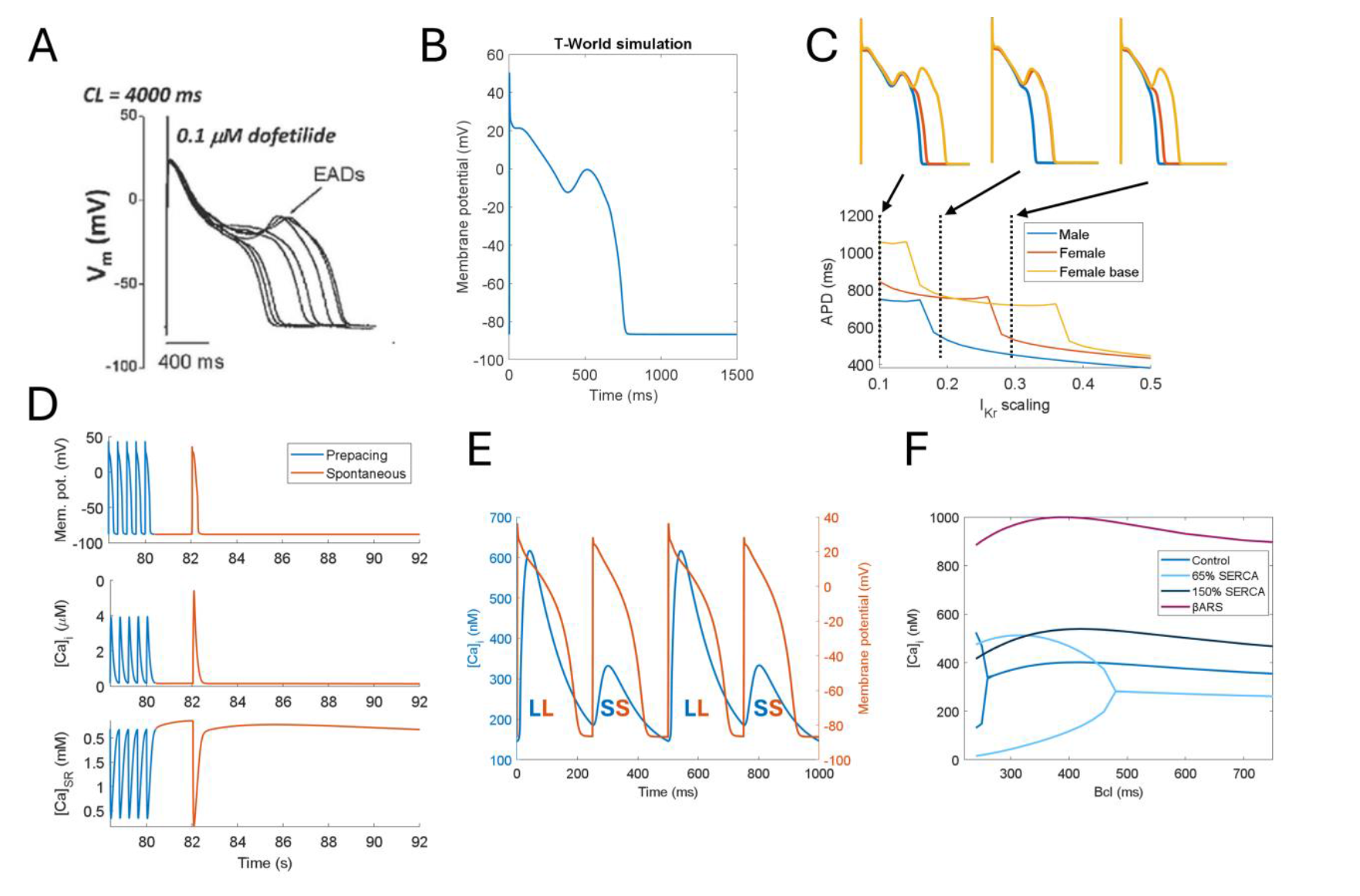
EADs, DADs, and alternans in T-World. **A)** Experimental data showing EADs at 0.25 Hz pacing, with 85% block of I_Kr_ with dofetilide^32^. **B)** EADs evoked in T-World under corresponding conditions. **C)** Demonstration of differential EAD formation under varying degrees of I_Kr_ availability in the following types of myocytes: male, female, and female + increased I_CaL_, reflecting the basal part of the heart in a rabbit study^33^. The y-axis shows action potential duration for a range of I_Kr_ scaling factors (fraction of current versus baseline) on the x-axis, with sharp transitions corresponding to changes in the number of EADs. Insets show APs at corresponding dashed lines. **D)** Examples of triggered activity resulting from DADs. The end of the pre-pacing train is shown in blue, with the spontaneous activity given in red. **E)** Illustration of concurrent oscillations in CaT and APD. LL = large/long CaT and APD respectively, SS = small/short. **F)** Modulation of calcium alternans by reduced and increased SERCA activity, as well as by βAR stimulation.

T-World also highlights sex differences in EAD vulnerability. The female T-World variant requires less I_Kr_ inhibition to induce EADs compared to the male variant, indicating greater EAD vulnerability (**Figure 2C**), supporting data showing higher risk of drug-induced arrhythmia in female hearts ^27,35^.

Additionally, female-specific apicobasal I_CaL_ gradients linked to estrogen^36^ suggest an additional risk of EADs in basal regions of the heart in female models. Spatially constrained EADs resulting from such gradients are likely to increase dispersion of repolarisation and may promote arrhythmogenesis at the tissue level beyond the EAD risk itself ^37^.

#### Delayed afterdepolarisations

DADs are arrhythmia triggers occurring during diastole. They result from spontaneous SR calcium release, generating inward currents (primarily via NCX) that depolarise the cell^38^, and are particularly prominent in diseased hearts, such as in heart failure^39^. DADs arise from stochastic subcellular calcium sparks best simulated by models with spatial calcium handling and stochastic gating ^40^. However, given the high computational cost of such models, ‘common pool’ models with similar complexity as T-World are often used to emulate DAD generation efficiently, especially Bers/Grandi-like models such as Morotti2021^11,13^. By contrast, ToR-ORd cannot produce any DADs due to its RyR activation mechanism, while TP06 can yield DADs following parametric changes, but these differ substantially from experimental recordings^41^.

T-World manifests spontaneous calcium releases and DADs (**Figure 2D**), and it can generate DAD trains, as observed in certain experiments^42^ (**Supplementary Figure S24**). Spontaneous releases are terminated when the SR content becomes sufficiently low following the spontaneous releases, similar to experimental parallel measurements of intracellular and SR calcium^43^. In this regard, our model differs from Morotti2021, where DADs stop occurring even when SR calcium keeps increasing (**Supplementary Figure S25**).

To validate DADs in the model, we confirmed that faster pre-pacing and RyR sensitisation promote DADs in T-World, as seen experimentally (**Supplementary Figure S26**). Furthermore, for applications requiring stochasticity of DADs, we developed a version of T-World that includes stochastic store-overload-dependent RyR release akin to the method by Colman et al.^44^ (**Supplementary Figure S27**).

#### Calcium and action potential alternans

Cardiac alternans, a periodic oscillation between long and short APDs, creates a pro-arrhythmic substrate, promoting conduction block ^45^ and increasing arrhythmia risk^46^. APD alternans is typically driven by underlying CaT oscillations and occurs at rapid heart rates^47^. While common at high pacing rates in living hearts, many computer models do not recapitulate it, including the Bers/Grandi family on which most of T-World calcium handling is originally based.

Thanks to its improved calcium handling, T-World produces AP and CaT alternans at realistic frequencies^48^, with mild alternans at 260 ms and pronounced alternans at 240–250 ms (**Figure 2E, Supplementary Figure S28**). Alternans is electromechanically concordant (long APD corresponds to large CaT), matching experimental data in human-relevant species^49–51^. T-World shows CaT alternans even when a fixed AP shape is imposed, confirming calcium oscillations as the primary driver (**Supplementary Figure S29**).

A major improvement of T-World compared to prior state of the art is its correct response of alternans to SERCA pump changes. Conditions like heart failure or pharmacological or transcriptional SERCA reduction increase alternans vulnerability^52–54^, with alternans appearing at slower pacing rates. T-World accurately reflects this by showing alternans at slower rates with SERCA reduction (**Figure 2F**). This is in contrast with ToR-ORd, where SERCA inhibition suppresses alternans (**Supplementary Figure S30**). Finally, we validated T-World by showing that an increase in SERCA function or βAR activation suppress alternans (**Figure 2F**), in line with experimental data^55,56^.

#### Steep S1-S2 restitution

Steep APD restitution promotes arrhythmias by facilitating reentry and spiral wave breakup in cardiac tissue^14,57^. It is measured using an S1-S2 protocol, where a premature stimulus (termed S2) follows a train of stimuli (termed S1), generating a curve relating APD to the preceding diastolic interval (**Figure 3A**). Slopes of the restitution curve greater than 1 facilitate proarrhythmia, and human studies show that maximum curve slopes slightly above 1 are not uncommon ^58,59^. The TP06 model has been historically popular, because of its steep restitution properties. At the same time, e.g. the ToR-ORd model, on which most of T-World’s electrophysiology is based, has a relatively flat restitution (peak slope ~0.5), limiting its utility in these aspects. However, our revised L-type calcium current and other developments lead T-World to exhibit good agreement with experimental restitution data (**Figure 3A**) and S1S2 slope >1 in a part of the curve for S1 interval of 1000 ms (**Figure 3B**). The importance of steep restitution is supported by the fact that T-World can reproduce VF (**Figure 1G**), unlike the prior ToR-ORd model, which required substantial adaptations to achieve VF.

**Figure 3.**
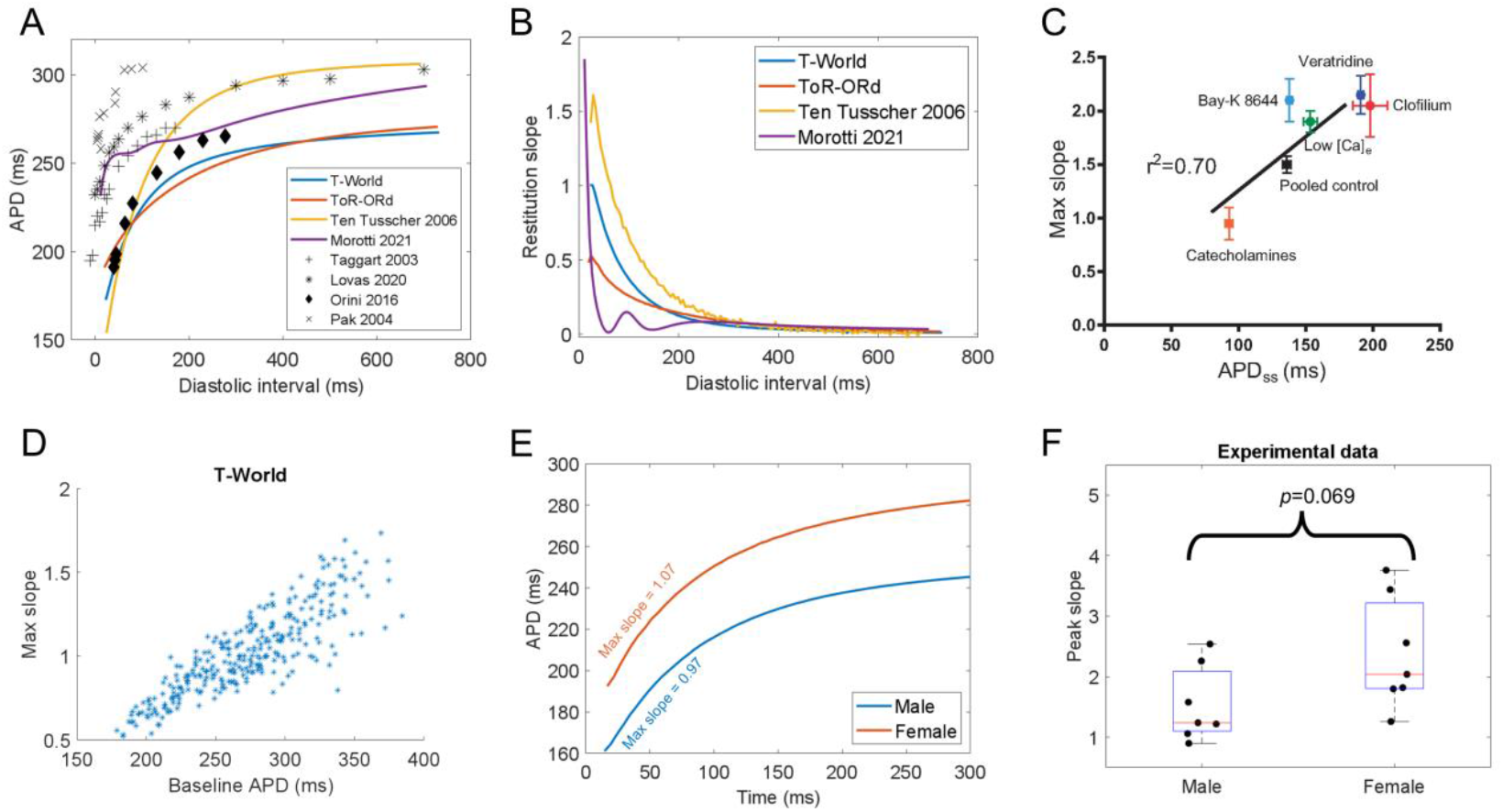
S1-S2 restitution and stability of arrhythmic behaviours. **A)** S1-S2 restitution curve in the T-World, ToR-ORd, Morotti2021, and TP06 models, and a range of human studies ^58,63–65^. **B)** Comparison of S1-S2 restitution slopes across T-World, ToR-ORd, Morotti2021, and TP06 models. **C)** Positive relationship between APD of a cell and its peak S1-S2 restitution slope, observed experimentally ^18^. **D)** Corresponding simulation in T-World showing that when a range of ion channel conductances are varied (see Methods for details), cells with longer APD generally show a steeper slope of restitution. **E)** Steepening of restitution in female versus male myocytes in baseline T-World. **F)** Experimental human data comparing peak restitution slope in males vs females (N=7 in both groups, p-value obtained using unpaired t-test).

In 2017, Shattock et al.^18^ showed that the maximum slope of the restitution curve is largely determined by the steady-state APD of a cell. Specifically, the longer the APD, the steeper the restitution (**Figure 3C**), which was corroborated by multiple studies using different means of changing APD ^60–62^. Importantly, T-World is the only model among those capable of steep restitution that recapitulates this feature (**Figure 3D**), with the TP06 model showing a weakly inverse APD-slope relationship, and the Morotti2021 a strongly inverse one (**Supplementary Figure S31**). This makes T-World uniquely suitable for studying how APD changes due to disease or drugs modulate arrhythmic risk via restitution changes.

One notable exception to the observation by Shattock et al. is the effect of βAR stimulation, which shortens APD, but steepens the S1S2 slope in humans^58^. As an independent validation, we simulated the effect of βAR activation in the T-World model, which correctly predicted the phenotype (**Supplementary Figure S32**). Furthermore, we validated that shortening of the S1 interval correctly flattens the S1S2 restitution (**Supplementary Figure S33**).

Measuring restitution slope separately for the male and female versions of T-World, we observed a steeper slope in the female myocyte (**Figure 3E, Supplementary Figure S34**). This would point to an increased risk of arrhythmia in female hearts through steeper restitution, but intriguingly, we were unable to find any experimental study addressing this hypothesis. However, we were able to obtain human ventricular data from the study by Lovas et al. ^63^ and re-analysed them for sex differences in peak slope. Mean (SD) peak slope in males was 1.54 (±0.63), increasing to a mean of 2.38 (±0.92) in females (p=0.069, t-test) (**Figure 3F**). This suggests that females may have steeper restitution properties, a previously underappreciated sex-specific hazard.

#### Stability of arrhythmic behaviours

To validate generality and robustness of T-World, we investigated the stability of the cellular arrhythmic behaviours under parameter perturbation using a population-of-models approach. While it is natural for cells, living and simulated alike, to manifest arrhythmogenic behaviours at slightly different conditions, the majority of cells should be fundamentally capable of manifesting them. In **Supplementary Note 6**, we demonstrate that T-World is highly robust with regards to its arrhythmia precursor capabilities, which are its intrinsic properties, rather than phenomena that only occur for highly specific sets of distinct parameters for each property.

### Using T-World to predict drug effects and elucidate arrhythmia mechanisms in disease

The robust representation of many cellular arrhythmia mechanisms makes T-World highly suitable to facilitate a better understanding of their role in (patho)physiological conditions. Here, we show how T-World can advance assessment of drug effects and provide insight into arrhythmogenic mechanisms in diseases that may include diabetes, heart failure, or monogenic arrhythmia disorders (e.g., the Long QT syndrome).

#### Drug safety and efficacy assessment

Prediction of drug safety through in silico trials is a successful translational application of mechanistic computer models, with considerable uptake by industry and regulators ^3^. This is crucial as cardiac side effects are a major cause of drug attrition and market withdrawal ^66^. To demonstrate utility of T-World for drug safety, we conducted an in silico trial using populations-of-models ^3,4^, comparing predictions against clinical risk data for 61 drugs (**Figure 4A**). Compared to prior works, we updated drug safety annotation based on the most recent version of the Crediblemeds classification ^67^, and pharmacological data for several compounds (see Methods).

**Figure 4.**
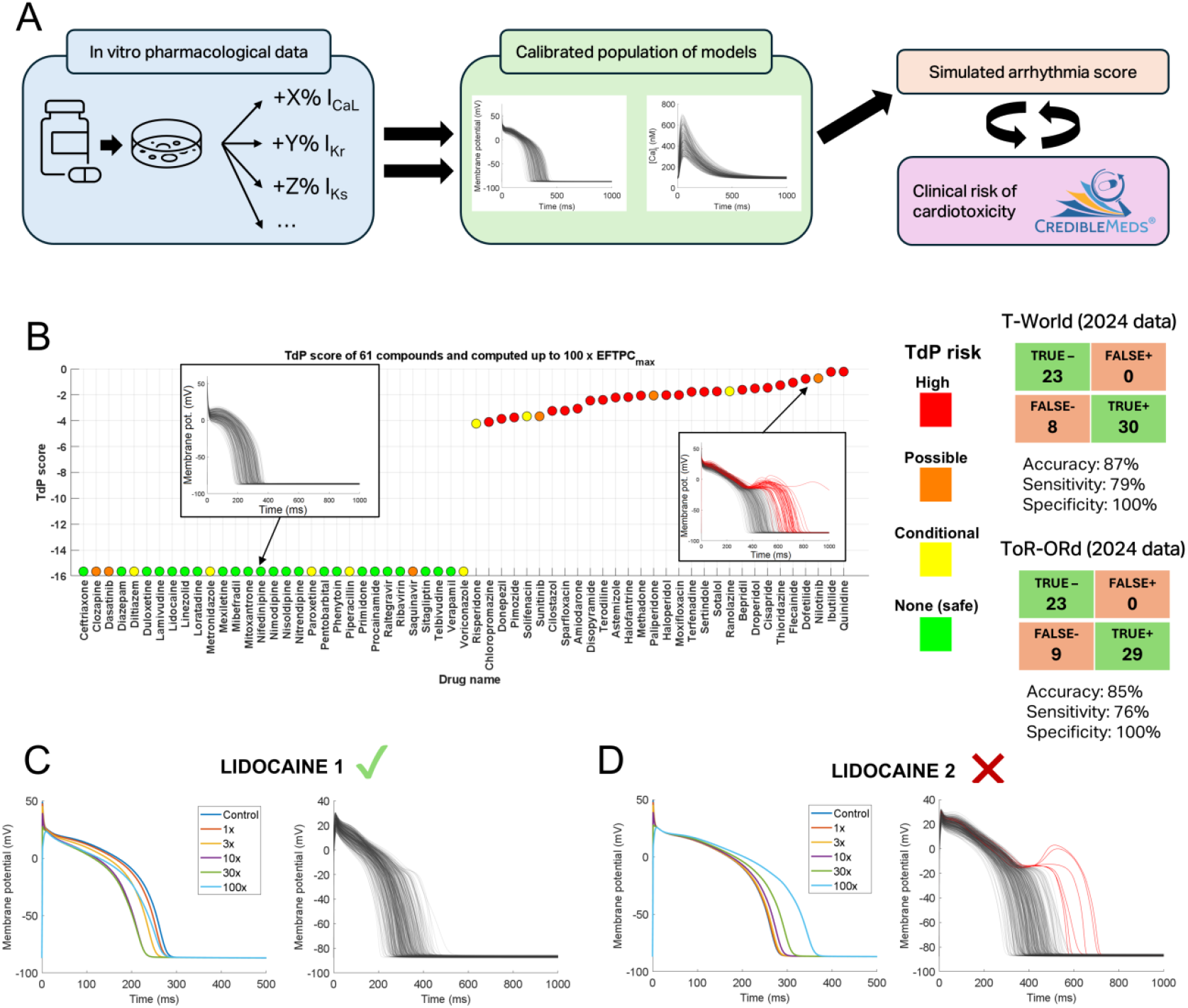
In silico drug safety and efficacy assessment. **A)** Schematic of the in silico drug trial procedure, showing how pharmacological data on dose-dependent inhibition of various currents by cardioactive drugs are applied to calibrated populations of models. Subsequently, arrhythmogenic behaviours are detected, scored, and the prediction can be compared to reference clinical risk. **B)** Predicted risk scores for 61 drugs with, with color-coded clinical risk, as in ^3,70^. Tables of true/false classifications are provided in the right part for T-World and ToR-ORd. Please see Methods for a summary of how several data updates lead to a subtly different performance of ToR-ORd in our study compared to the original publication^4^. **C)** Effect of the first lidocaine description (with I_NaL_ effect) on AP to the left, showing overall safety to the right (no model in the model manifests an EAD). **D)** Similar plot for the second lidocaine description available in the database, showing gradual dose-dependent APD prolongation to the left and repolarisation abnormalities to the right. In C, D, lidocaine effect is shown at the maximum concentration of 100x.

A population of 343 models, constrained by experimentally informed human reference ranges for APD, CaT, and contraction biomarkers (**Supplementary Figure S35**) was exposed to 61 drugs at doses up to 100 times therapeutic levels. The model correctly predicted all no-risk drugs as safe and all high-risk drugs as unsafe. Classifying the drugs into two categories of safe (*no risk*) and unsafe (*high, possible*, or *conditional* risk) yielded a prediction accuracy of 87%, with 79% sensitivity and 100% specificity (**Figure 4B**). This represents an improvement over the prior ToR-ORd model, highlighting the robustness of T-World despite its entirely different calcium-handling system and revised ion current formulations.

A drug effect prediction can be reliable only when the underlying drug description data are accurate. T-World can identify incorrect pharmacological descriptions of drugs, which can limit prediction accuracy. When a drug with known phenotypic effect (e.g. changes to APD or contractility) is simulated, a discrepancy between the simulation and known reality indicates that an important effect of the drug is not included in the drug description data. We illustrate this using lidocaine, a safe sodium-channel blocker, where one of two available descriptions was excluded a priori during data curation. Both versions block peak I_Na_ and weakly block I_Kr_, with one additionally potently blocking I_NaL_. Lidocaine is known to shorten APD^68^, and this is recapitulated only by the version including the I_NaL_ block, indicating its superiority (**Figure 4C,D**). Interestingly, the incorrect description generates EADs and falsely indicated arrhythmic risk (**Figure 4C,D**), highlighting the need to exclude incomplete drug descriptions. In this case, exclusion of the non-I_NaL_ formulation is independently supported by studies directly demonstrating I_NaL_ inhibition by lidocaine^68^.

Similarly, we also identified an inaccuracy in the description of cilostazol, with the original drug description failing to predict the effect of the drug on contractility. This resulted from its arguably main effect of PDE3 inhibition not being included in the pharmacological data, which focused on ion channel blockade only (**Supplementary Figure S36**).

Finally, T-World can be used for studies on drug efficacy, either by identifying promising combinations of single channel blocks, or by disentangling different pro- and anti-arrhythmic effects of drugs with complex multi-channel profiles. Recently, the multichannel blocker mexiletine was proposed against Long QT syndrome 2 (LQTS2) ^69^ caused by APD prolongation due to loss-of-function mutations in I_Kr_. In **Supplementary Note 7**, we unpick the positive effect of mexiletine in a LQTS2 version of T-World, linking it to dual inhibitory effect of the drug on I_NaL_ and I_CaL_, which outweigh its I_Kr_-blocking effect. Further research is required to assess whether the drug blocks I_Ks_, which could be problematic during βAR activity.

#### Assessing arrhythmogenesis in type-2 diabetes

The generality of T-World enables the creation of predictive disease-specific models. T2D is a major 21st-century epidemic linked to increased mortality, with cardiovascular disease as the leading cause of death. Sudden cardiac death from ventricular arrhythmia is the primary driver, yet the mechanisms behind ventricular arrhythmogenesis in T2D remain poorly understood ^71^. Limited human data on ionic currents and calcium-handling proteins ^72^ show only partial alignment with heterogeneous animal studies. Progressive cardiac remodelling in T2D further complicates consistent disease characterisation.

A significant portion of sudden cardiac deaths in T2D occur in patients using potentially proarrhythmic drugs^71^. This suggests hidden cardiotoxicity in T2D, with usually safe drugs (or safe drug concentrations) becoming dangerous. To investigate this, we created a range of models (D1-D6) reflecting key T2D phenotypes from literature, primarily using human data supplemented by animal studies (**Supplementary Methods – T-World applications**). All models exhibited AP prolongation (**Figure 5A**), consistent with clinical QT prolongation in T2D ^73–75^. All diabetic model variants are more vulnerable to EADs (**Figure 5B)**, requiring less I_Kr_ inhibition to trigger EADs. This includes those with reduced I_CaL_ (D3-D6), which could be thought to be more protected. Key drivers of EAD risk were I_Kr_ reduction and I_NaCa_ increase, further heightened by CaMKII hyperactivity and increased I_CaL_ in D1-D2 (**Figure 5C**). I_CaL_ reduction alone (a component of D3-D6) showed reduced risk but not enough to offset other remodelling effects. This suggests that across different formulations of T2D remodelling, T2D patients require less I_Kr_ inhibition to manifest EADs, therefore facing higher arrhythmia risk at drug doses considered safe for non-diabetic individuals.

**Figure 5.**
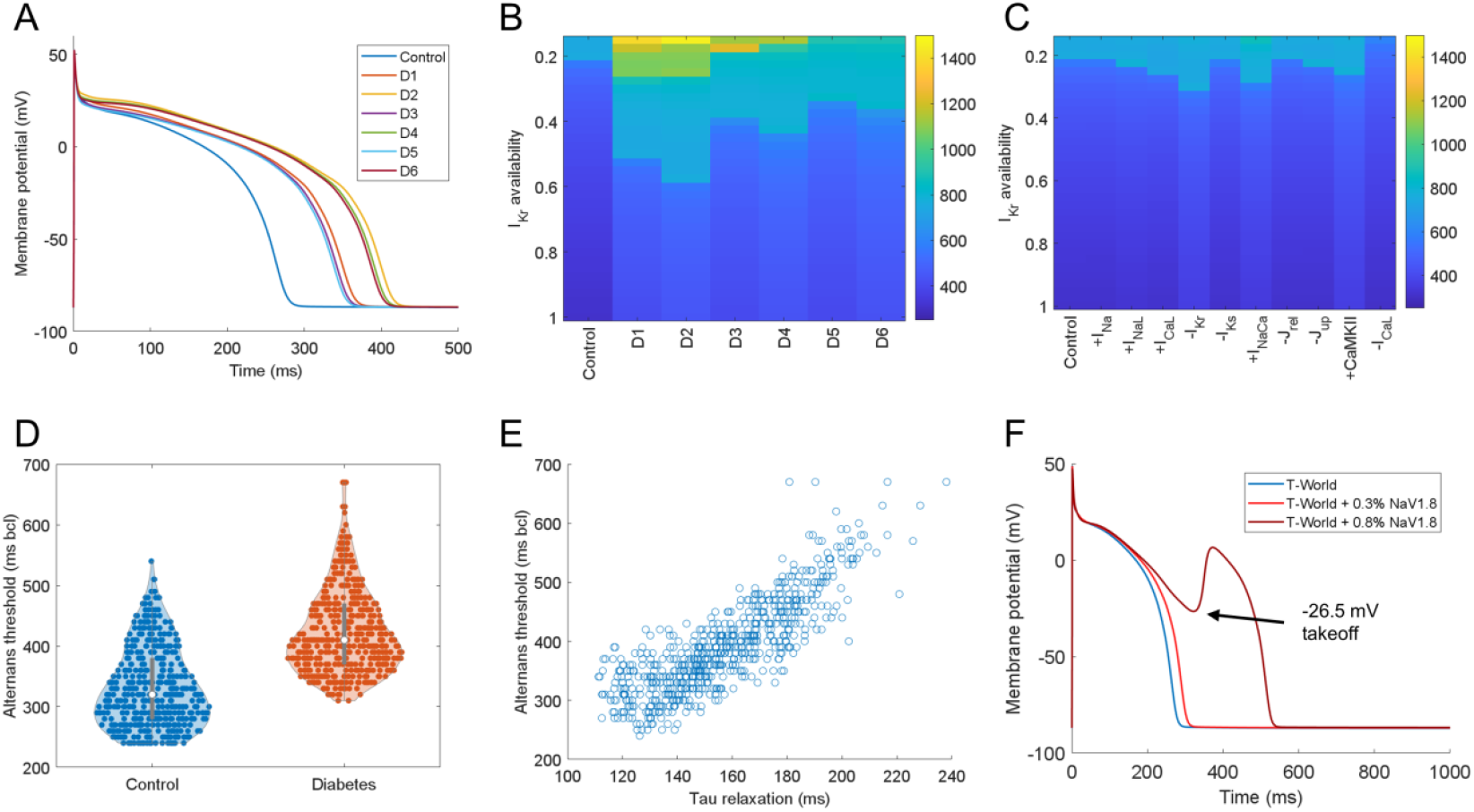
Arrhythmogenesis promotion by Type 2 diabetes and Na_V_1.8 current. **A)** Comparison of APs of six distinct formulations of T2D remodelling (see Methods for details. **B)** Action potentials of control and T2D models across a range of I_Kr_ multipliers (on y-axis). Sharp transitions between colours indicate a change in the presence of an EAD. Specifically, the lowest teal point on the y-axis (APD of ca. 900) indicates the highest I_Kr_ availability which supports EAD formation. **C)** The same type of visualisation showing how single elements of T2D remodelling considered throughout D1-D6 models, added to the control model, alter EAD vulnerability. The ‘+CaMKII’ column also involves an increase in I_NaL_, as described in Methods, and ‘−I_CaL_’ corresponds to 80% I_CaL_ density compared to control model. **D)** Comparing alternans vulnerability (slowest pacing rate which induces CaT alternans) between population of control vs T2D models. The calibrated population of 796 models used in Results: Stability of arrhythmic behaviours was used as the control population, with T2D models created by adding diabetic remodelling to each of those models. **E)** A scatterplot of tau of mechanical relaxation versus alternans threshold in the simulated T2D population **F)** Comparing control T-World model AP to APs obtained when two different amounts of NaV1.8 current are added (expressed as relative percentage of peak I_Na_).

A different arrhythmogenic behaviour that is markedly increased in T2D patients is alternans^76^. We used the D1 version of T-World, which has recapitulated the clinical observation, showing alternans at slower pacing compared to non-diabetic versions (**Figure 5D**). Interestingly, Bonapace et al. furthermore observed that alternans vulnerability is positively associated with and diastolic dysfunction in T2D patients^77^. To investigate this phenomenon in T-World, we correlated the slowest pacing rate for CaT alternans with diastolic function (tau of relaxation) in a population of models with perturbed parameters. Models prone to alternans exhibited impaired relaxation, consistent with clinical data (**Figure 5E**). We hypothesised and subsequently confirmed that reduced SERCA pump function, crucial for relaxation and calcium clearance, can causally drive this relationship (**Supplementary Note 8**).

#### Na_V_1.8 can drive EADs in a diseased heart

Cardiac disease may remodel ionic currents active under physiological conditions, but it can also involve expression of nonstandard ionic currents, absent in a healthy heart. We employed T-World to investigate the role of Na_V_1.8, a primarily neuronal sodium channel subtype with recently much debated functionality in the heart. While Na_V_1.8 is minimally expressed in healthy hearts^78^, it appears in hypertrophic or failing hearts, and may contribute disproportionately to the late sodium current I_NaL_^79,80^. Increased I_NaL_ can in general promote arrhythmias by prolonging APD (leading to EADs) or increasing sodium influx, reducing NCX calcium efflux and causing DADs. However, given Na_V_1.8’s unique biophysical properties, including right-shifted activation and inactivation compared to Na_V_1.5 (**Supplementary Figure S37**), we hypothesised it could directly generate EADs by providing depolarising current during the late AP plateau.

Introducing a small Na_V_1.8 current (~0.3% of peak I_Na_) to T-World prolonged APD (**Figure 5F**), consistent with its role as an I_NaL_ source. Strikingly, increasing Na_V_1.8 by 2.75-fold (to only 0.8% of peak I_Na_) triggered EADs at 1 Hz pacing (**Figure 5F**). These EADs emerged at a take-off potential of −30 mV, clearly distinct from I_CaL_-driven EADs at −13 mV (**Figure 4B**). Simultaneous tracking of Na_V_1.8 current and I_CaL_ during EADs revealed that Na_V_1.8 initiates depolarisation, subsequently activating I_CaL_ in a dual-current process (**Supplementary Figure S38**). Therefore, Na_V_1.8 can directly trigger EADs rather than merely prolong APD to facilitate I_CaL_ reactivation, potentially co-explaining elevated arrhythmic risk in those patients^81^. This mechanism suggests Na_V_1.8 as a possible anti-arrhythmic target, e.g., providing additional rationale for the use of ranolazine, which is protective in the hypertrophied heart^20,82^, and which blocks Na_V_1.8^83^. Future work on Na_V_1.8 modelling is suggested in **Supplementary Note 9**.

## Discussion

Here, we present the development, calibration, validation, and application of T-World, a novel computer model of the human ventricular cardiomyocyte. This model addresses a longstanding but unresolved need for a highly general virtual cardiac cell model with robust baseline physiology, capable of replicating all key cellular mechanisms of arrhythmia. The model utilises a range of new developments, as well as components and ideas from two influential modelling families from the Rudy^4,19^ and Grandi and Bers labs^11,12^. It is the first model to unify those two different approaches to modelling cardiac cells, improving upon their strengths, while resolving their known limitations. In addition to good representation of DADs stemming from the Bers/Grandi framework^11,12^ and presence of EAD in relevant conditions such as in ToR-ORd^4^, the model now manifests data-like restitution properties and calcium-driven alternans which correctly responds to changes in SERCA pumps. The model gains credibility through rigorous independent validation on unseen data, its construction from well-characterised components, and its human-specific design, which bypasses the species differences inherent to animal experiments (elaborated in **Supplementary Note 10**). Its universality, predictivity, and adaptability make it an excellent tool for basic cardiac research at cellular and organ level, pharmaceutical applications, and development of patient virtual twins ^8^.

T-World presents an important step towards realising the vision of the 3Rs: reduction, refinement, and (partial) replacement of animal use in research and industry. It can also work synergistically with *in vitro* models such as induced pluripotent stem cell-derived cardiomyocytes, helping interpret their so far still typically immature phenotype in the context of the adult heart^84^. T-World’s human nature can also be leveraged to utilize animal-based measurements (such as protein level changes) and predict their functional implications for the human heart, thereby “humanising” the data.

The fact that T-World directly represents cellular biology means that it can be used to investigate the modulation of its components by drugs and/or disease-related alterations. It can be for example used to understand mechanisms of high arrhythmia risk in a given disease, and then help discover drugs that can ameliorate such a risk. We used T-World to disentangle mexiletine’s antiarrhythmic effects in Long QT syndrome type 2, showing it results from a dual I_NaL_ and I_CaL_ blockade. Such insights may be also used in future to explore new therapeutic combinations of distinct drugs. T-World is furthermore suitable for carrying drug arrhythmia studies in a sex-specific manner, having reproduced the higher risk of drug-induced arrhythmia in females ^27,35^. Finally, with its representation of contractility and improved excitation-contraction coupling, T-World is well-suited for studying drug effects on contractility. This is another rapidly developing domain of applications with high relevance for industry^85^. An advantage of the comprehensive nature of T-World over single-purpose predictors is that it can be used to address compound queries, such as “find the most anti-arrhythmic drug for a given condition without compromising contractility”.

T-World will be useful in preclinical drug safety testing, one of the most established translational applications of non-animal methods, with significant industry adoption. We demonstrate T-World’s excellent performance in drug safety testing through population-of-models in silico trials, slightly surpassing the prior state-of-the-art ToR-ORd^4^. We believe that the main barrier to improved drug safety prediction now lies in data quality rather than model quality, as shown by our data curation process. Notably, we introduce a novel use of T-World simulations to identify inaccuracies or missing data in drug action datasets, based on the capability of the drug description data to reconstruct known phenotype. Discrepancies between simulated and observed drug effects on e.g., action potential or contractility can highlight missing mechanisms in drug data description, guiding additional measurements. Missing effects likely contribute to misclassification of several drugs in our study (**Supplementary Note 11**).

Notably, while the drug-induced arrhythmia risk mostly results from EADs, our study indicates via simulations and subsequently analysed human data that females are also at a higher risk of a steep restitution slope. At the same time, this is known to promote arrhythmia at the tissue level^14,57^. This finding highlights how T-World can be used to drive discovery in biological data. However, given the small sample size and exploratory nature of the data, further larger-scale studies are needed. Considering the elevated risk of EADs and steep restitution in females, our study reinforces the urgency of sex-specific drug dosing and treatment guidelines, which is currently not sufficiently addressed^86^.

Arrhythmias pose a significant risk in heart disease, and T-World’s improved baseline physiology and arrhythmic behaviour make it ideal for studying complex cardiac diseases. Using T-World, we constructed a pilot cell model for Type 2 diabetes (T2D), a condition with high arrhythmic burden but limited mechanistic understanding^71^. The model revealed increased risks of EADs and alternans, which can explain elevated rates of sudden cardiac death in this population.

Furthermore, the higher EAD risk indicates a heightened vulnerability to drug-induced arrhythmia, a major concern in T2D^71^. The suggested strong involvement of NCX in the elevated EAD risk may warrant investigation of therapeutic potential of NCX blockers such as ORM-10962, which inhibit both NCX and I_CaL_ (both pro-EAD factors) while not compromising contractility ^87^. At the same time, ORM-10962 was shown to inhibit alternans experimentally^88^, possibly targeting also the second pro-arrhythmic aspect in T2D. Despite promising results achieved, we note the urgent need to collect new, high-quality human datasets to characterise and understand how T2D dysregulates the heart, given the paucity of existing data.

T-World’s realistic calcium handling and ECC make it well-suited for diseases with significant calcium remodelling, such as heart and post-infarction remodelling. Unlike models like ToR-ORd, T-World can produce DADs, important in such diseases ^39^. A particular strength pertaining to arrhythmia mechanisms is that T-World exhibits increased alternans vulnerability with reduction in SERCA pumps, both hallmarks of those diseases ^52,53^. This is an improvement over major prior human models such as Grandi et al. ^12^ which lacked alternans, or ORd and ToR-ORd^4,19^, which do not respond well to SERCA changes.

T-World can be applied to study the role of nonstandard channels absent in healthy hearts, but present in disease. Here, we investigated the role of the “brain-type” Na_V_1.8 channel, which is known to be absent in healthy hearts ^78^, but can appear in disease, such as heart failure ^79,80^. We show that even if present in small amount, NaV1.8 may directly contribute to arrhythmia in disease through its unusual gating properties, highlighting it as a potential treatment target.

The inclusion of βAR signalling and excitation-contraction coupling modulation makes T-World highly suitable for exploring the neurocardiac axis in arrhythmia and sympathetic nervous system studies^89^. It is particularly well applicable for studies on arrhythmia and sympathetic nervous system, given that the validation has demonstrated strong predictive performance with regards to modulation of multiple arrhythmic mechanisms by sympathetic nervous activity.

Several limitations of T-World, most of which are intrinsic to the level of detail modelled, are given in **Supplementary Note 12**.

T-World opens new avenues in cardiac research. We envision that its universality and open-source nature will enable adaptation to represent other excitable cells, such as atrial, sinoatrial, Purkinje, or neuronal. It will also facilitate studies on newly discovered ionic currents, and on understanding signalling pathways and how they modulate the cellular physiology. To expand the model generality, we anticipate it will be particularly important to represent dynamic regulation of trafficking and transcription ^90^ enabling studies on long-term remodelling, representation of metabolism and reactive oxygen species^91^, integration with AI-driven structural modelling^92^ and omics analyses^93^.

## Methods

T-World is a virtual cell model using sets of ordinary differential equations to describe, based on experimental data, the dynamics of ionic currents, fluxes, and subcellular signalling. The overall cell architecture and calcium handling were mainly inspired by the Bers/Grandi family of models^11–13^, with most ionic current formulations being inspired by the ToR-ORd model^4^. In order to enable all key arrhythmic behaviours in relevant conditions, and to avoid limitations of these frameworks with regards to basic physiological behaviours and response to (patho)physiological changes, we introduced numerous innovations, such as a new hybrid approach to coupling L-type calcium current and ryanodine receptors, new L-type calcium current model, heavily revised model of the ryanodine receptor, re-developed model of sodium-potassium pump, and a wide array of changes to most cell components. The ‘World’ in the model’s name reflects the fact that model designs and expertise from the whole world were essential in its creation, and it goes beyond outputs of a single group.

Please see **Supplementary Methods** for a detailed description of the following:

1. Model architecture
2. Calibration and validation criteria for T-World development and evaluation
3. Description of equations describing the ionic currents and fluxes
4. Contractility representation
5. CaMKII and β-adrenergic signalling
6. Sex differences
7. Organ-level simulation methodology
8. Methodology for studies on arrhythmogenic behaviours.
9. Methodology for sample applications: in silico trials, type 2 modelling, and NaV1.8 current investigation
10. Graphical user interface
11. Notes on implementation
12. Sources of implementation of other models

T-World is distributed as open-source code and is available at **<will-be-provided-upon-acceptance>**, including sample scripts demonstrating its functionality. An online graphical user interface enabling running T-World simulations is available at **<will-be-provided-upon-acceptance>**. Background of the T-World development, including the description of various dead ends that we encountered during development, will be provided at the blog underlid.blogspot.com.

## Acknowledgments

J Tomek is supported by the Sir Henry Wellcome Fellowship (222781/Z/21/Z). J Heijman was supported by the Netherlands Heart Foundation (grant no. 01-002-2022-0118, EmbRACE: Electro-Molecular Basis and theRapeutic management of Atrial Cardiomyopathy, fibrillation and associated) and the Netherlands Organization for Scientific Research (NWO/ZonMW Vidi 09150171910029). D Bers is supported by NIH grants P01-HL141084 and R01-HL092097. This work was also supported by a Wellcome Trust Fellowship in Basic Biomedical Sciences to B Rodriguez (214290/Z/18/Z) and the CompBioMedX project (to B Rodriguez., EP/X019446/1). This study used high-performance computing resources from the Polaris supercomputer at the Argonne Leadership Computing Facility (ALCF), Argonne National Laboratory, United States of America. The US Department of Energy’s (DOE) Innovative and Novel Computational Impact on Theory and Experiment (INCITE) Program awarded access to Polaris. The ACLF is supported by the Office of Science of the US DOE under Contract No. DE-AC02-06CH11357. The project was further supported by the National Research Development and Innovation Office (NKFIH FK-142949 for N Nagy). TM Bury is supported by a Fonds de Recherche du Québec - Nature et technologies (FRQNT) postdoctoral fellowship. A Bueno-Orovio acknowledges support from the Innovate UK grant 10110728. M Colman is supported by Medical Research Council Career Development Award (Grant Number MR/V010050/1).

We thank Eleonora Grandi, Stefano Morotti, Haibo Ni for useful discussions on how models derived from Shannon et al. operate. We thank Dirk Gillespie, Dezso Boda, Pavel Jungwirth, and Geir Halnes for their insights on how ionic driving force through open L-type calcium channels should or should not be modelled.

For the purpose of open access, the authors have applied a Creative Commons Attribution (CC-BY-NC) public copyright licence to any Author Accepted Manuscript version arising from this submission.

