## Supplementary Materials for "T-World: A highly general computational model of a human ventricular myocyte"

### Supplementary Methods

#### Model architecture

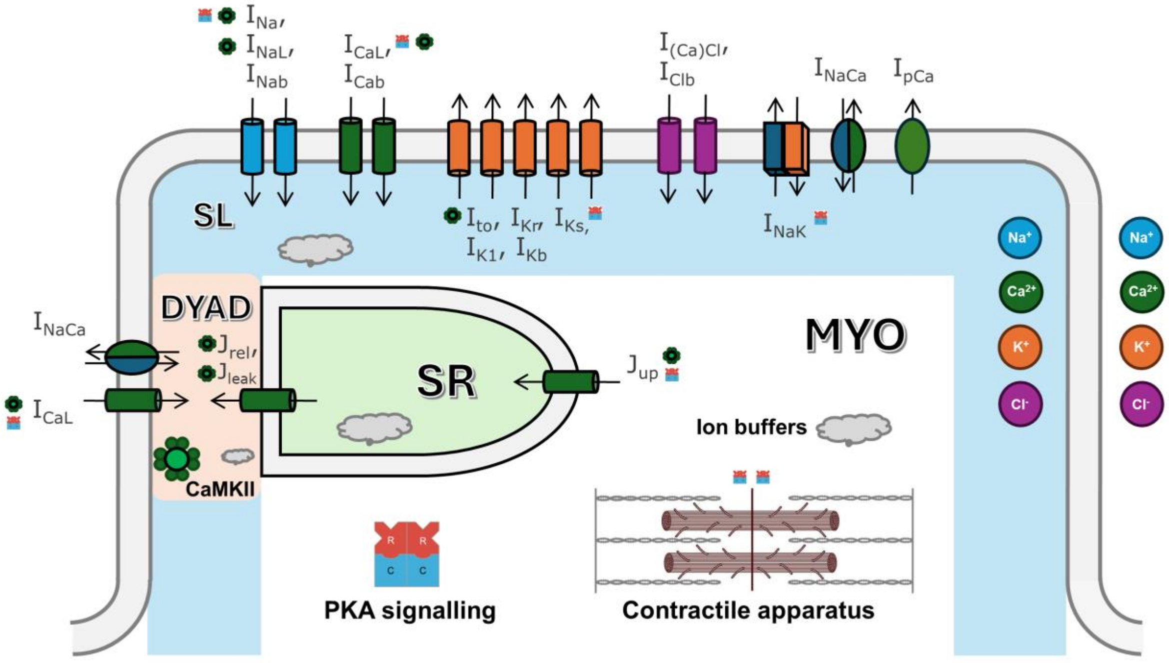

***Figure S1. Diagram of T-World.*** *In the diagram is shown the membrane with ionic currents (color-coded by ionic species). The inside of the cell is separated into the following compartments: DYAD (orange), subsarcolemmal (SL, blue), bulk myoplasm (MYO), sarcoplasmic reticulum (SR, green). CaMKII and PKA signalling pathways are included, with small icons adjacent to names of ionic currents, fluxes, or contractile apparatus indicating the site as a target of a given signalling pathway. Gray clouds indicate ionic buffers.*

The overall cell architecture and compartmentalization of T-World (**Figure S1**) is based on the architecture introduced in the Shannon et al. rabbit model [1]. We retained the compartment volumes and localizations of buffers. The majority of ionic currents and fluxes were replaced by alternative formulations, as described below.

A substantial update is that intracellular potassium and chloride concentrations are dynamic in T-World, being updated according to ionic currents, unlike the Shannon model and later models in the same family (e.g. [2,3]), where calcium and sodium were dynamic, but potassium and chloride were at fixed concentrations. Given the relatively uniform concentrations of potassium and chloride in a cardiomyocyte, we represent only a single intracellular concentration of those ionic species to keep the model simple and efficient. By contrast, sodium and calcium have separate concentrations in dyad, SL, and MYO compartments. Most ionic currents are localized similarly to Shannon-like models, with 89% localized in the SL compartment, with 11% placed in the dyad. The exceptions are $I_{CaL}$ (80% in dyads), $I_{NaCa}$ (31% in dyads), and $I_{(Ca)Cl}$ (50% in dyads). For potassium and chloride currents, the reversal potential was calculated based on the extracellular concentration and the single, homogeneous intracellular concentration.

The model has three versions according to transmural localization: endocardial, epicardial, and midmyocardial. The endocardial model is used as a default version throughout the article unless stated otherwise. Finally, T-World uses a conservative potassium stimulus [4], improving model stability compared to nonconservative-stimulus models.

#### Calibration and validation criteria

T-World was developed and calibrated to reproduce a wide range of pre-defined behaviours (**Table S1**), which was followed by validation on data not used in development (**Table S2**). The behaviours were chosen to reflect key properties of myocyte physiology and excitation-contraction coupling, and their modification by treatments and conditions with a known effect. Among the calibration criteria, we note that the criterion on alternans being promoted by SERCA reduction was fulfilled spontaneously once we were able to develop the Ca handling model to recapitulate alternans, without the development process looking at this feature. While it was included as a calibration criterion in later development stages to make sure the property is not lost, the fact that it emerged spontaneously serves as semi-validation.

On the other hand, the criterion on APD-slope relationship being positive was initially included as a validation criterion in the T-World development. However, this was failed by a model we considered near-final several years back, which warranted investigation of the issue and consequent replacement and redevelopment of the $I_{CaL}$ model. Given that this criterion was ultimately used in the model development, it is listed as a calibration criterion.

***Table S1. Calibration criteria of T-World.***

| Feature | Data |
|---|---|
| Action potential morphology | [5] |
| Calcium transient morphology | [6] |
| Dyadic calcium time to peak near 10 ms | [7–10] |
| Contraction biomarkers | [11] |
| $I_{CaL}$ I-V relationship | [12] |
| $I_{CaL}$ recovery from refractoriness | [13] |
| Positive correlation between CaT amplitude and SERCA levels | [14–17] |
| Biphasic rate dependence of CaT | [18,19] |
| Biphasic rate dependence of developed force | [18,20–23] |
| Monotonically positive rate-dependence of calcium in the SR | [20,24] |
| β-adrenergic activation shortens APD | [25,26] |
| EADs occur in experiment-like conditions | [27] |
| DADs can be evoked in the presence of β-agonist and elevated extracellular calcium | [28] |
| Alternans occurs at human-like rates | [29] |
| Alternans coupling between APD and CaT amplitude is positive | [24,30,31] |
| Alternans is potentiated by SERCA reduction and in diseases with reduced SERCA expression, and is inhibited by SERCA increase and β-AR stimulation (which increases SERCA) | [29,32–39] |
| S1S2 restitution has data-like shape | [40–43] |
| Peak slope of S1-S2 restitution exceeds 1 slightly | [40,42–45] |
| Relationship between cellular APD and its peak restitution slope is positive | [46] |

***Table S2. Validation criteria of T-World.***

| Feature | Data |
|---|---|
| Timing of $I_{CaL}$ voltage-dependent inactivation, as well as combined voltage- and calcium-dependent inactivation, is similar to data. | [5] |
| Appropriate rate-dependent response of APD to channel blockers (E-4031, HMR-1556, nisoldipine, mexiletine) | [5] |
| Negative inotropy of sodium blockers | [47–49] |

| | |
|---|---|
| Positive rate-dependence of $[Na]_i$ | [50] |
| During excitation-contraction coupling, most calcium is removed from cytosol by SERCA pumps, 20-30 by NCX, with a minimal contribution from pCa (sarcolemmal calcium pump) | [51] |
| Longer APD with slightly smaller CaT amplitude and contractility in female vs male cardiomyocytes | [52–55] |
| Female myocytes more vulnerable to EADs than male cardiomyocytes | [56–59] |
| β-AR stimulation markedly increases CaT amplitude | [60,61] |
| β-AR stimulation markedly increases contractility | [61–63] |
| β-AR stimulation steepens restitution | [43,64] |
| β-AR stimulation inhibits alternans | [34,65] |
| Shortening S1 flattens S1-S2 restitution slope | [43] |
| Faster prepacing promotes DAD formation | [66,67] |
| RyR sensitization promotes DAD formation in the presence of β-AR stimulation | [68] |

#### Ionic currents and fluxes

This section summarizes the formulations of ionic currents and fluxes in T-World. Baseline conductances are given in **Table S3**.

***Table S3. Conductances and transport rates of ionic currents and calcium handling fluxes in T-World.***

| Parameter | Maximum conductances and transport rates |
|---|---|
| $G_{Na}$ | 22.08788 |
| $G_{NaL}$ | 0.04229 |
| $p_{CaL}$ | 1.5768e-04 |
| $G_{to}$ | See corresponding section below |
| $G_{Kr}$ | $0.043 \cdot \sqrt{\frac{K_o}{5}}$ |
| $G_{Ks}$ | Calcium-dependent, see section below. |
| $G_{K1}$ | 0.6992 |
| $G_{NaCa}$ | 0.00179 |
| $\bar{I}_{NaK}$ | 2.10774 |
| $G_{PCa}$ | 0.02064 |
| $G_{(Cl)Ca}$ | 0.01615 |
| $G_{Clb}$ | 0.00241 |
| $G_{Nab}$ | 0.000594 |
| $G_{Cab}$ | 5.15575E-04 |
| $G_{Kb}$ | 0.010879 |
| $V_{max,SERCA}$ | 0.00543 |

Traces of key ionic currents at 1 Hz stimulation are given **in Figure S2**.

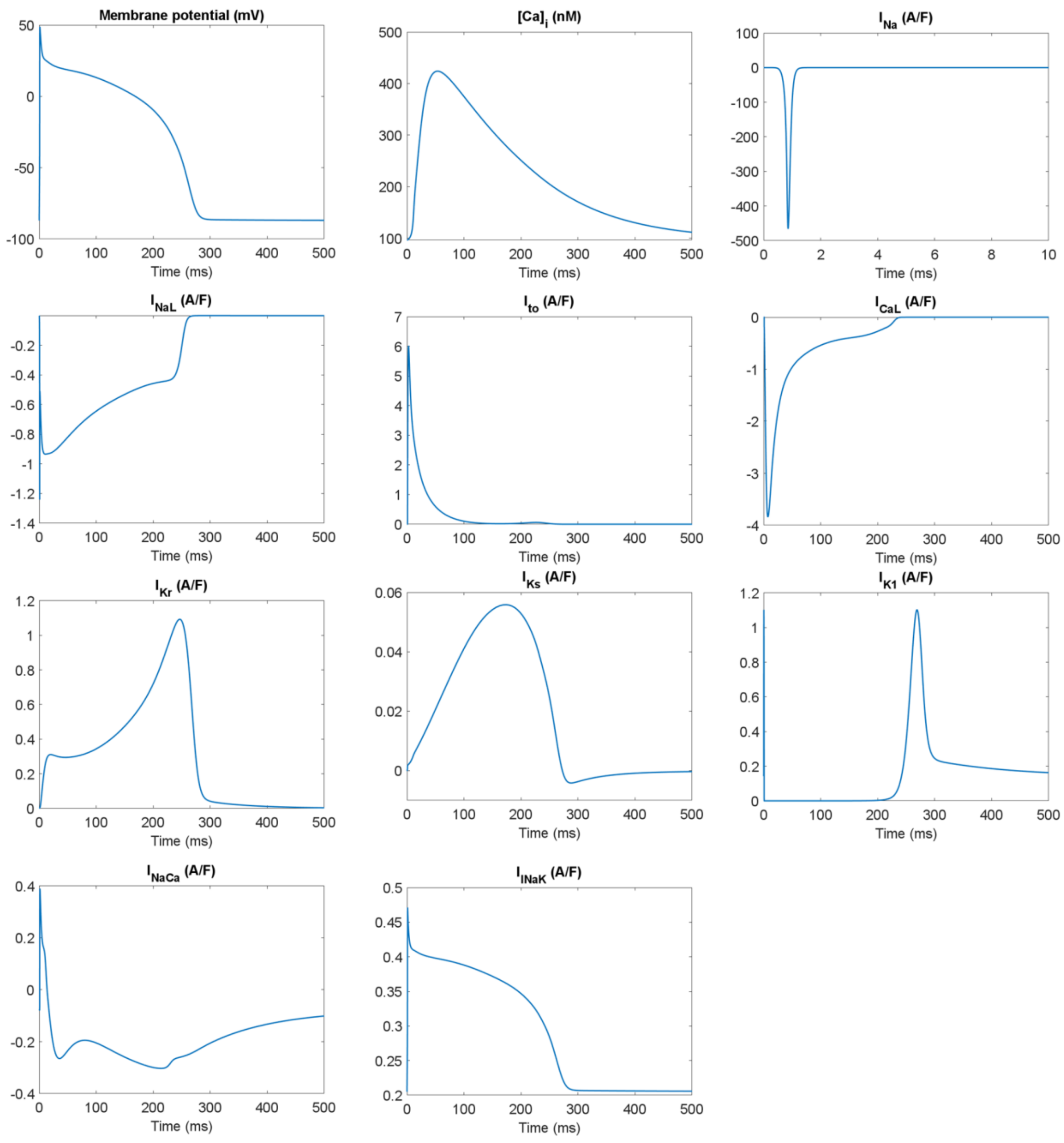

***Figure S2. Traces of key ionic currents in T-World.*** *Please note the different x-axis for $I_{Na}$ (top right), used given its extremely rapid activation and inactivation.*

#### Sodium current ($I_{Na}$, $I_{NaL}$)

The fast sodium current $I_{Na}$ was implemented based on ToR-ORd, extended with PKA-dependent phosphorylation, similar to Doste et. al. [69]. This yields four distinct populations of channels: unphosphorylated, PKA-phosphorylated, CaMKII-phosphorylated, and dual-phosphorylated. Compared to Doste et al., full PKA activation in our model induces 25% increase in $I_{Na}$ conductance, as used previously by Heijman et al. [70], and a small shift (5 mV) of the voltage-dependent $I_{Na}$ activation curve, similar to canine experimental data [71].

The late sodium current is based on ToR-ORd [72], with $\tau_{hL}$ reduced from 200 ms to 145 ms. This yields an $I_{NaL}$ profile that decreases throughout the AP, as observed in human myocytes (this differs from guinea pigs, where $I_{NaL}$ increases throughout most of the AP) [73]. Epicardial cells have 70% $I_{NaL}$ of the endocardium.

#### L-type calcium current ($I_{CaL}$)

The $I_{CaL}$ model is originally inspired by the formulation in ToR-ORd [72]. Beyond numerous parametric changes to its predecessor, it also includes an additional calcium-dependent inactivation gate $f_{Ca,CDI}$. This helps the model achieve a leaner $I_{CaL}$ profile, while maintaining the capability of manifesting EADs under experiment-like conditions. The leaner profile is advantageous in that much less $I_{Kb}$ is required to achieve a data-like AP shape, compared to ToR-ORd.

As in ToR-ORd, 80% $I_{CaL}$ is in the dyad, with 20% in the subsarcolemmal compartment.

βARS has been incorporated following the approach of Doste et al. in the sense that four distinct populations of $I_{CaL}$ are simulated: unphosphorylated, CaMKII-phosphorylated, PKA-phosphorylated, and dual-phosphorylated. However, PKA phosphorylation increases $I_{CaL}$ conductance 1.9-fold, which is in line with the substantial increase observed experimentally [74,75]. In addition, PKA phosphorylation shifts $I_{CaL}$ activation and inactivation leftward by 9 and 6 mV respectively, similar to experimental data [74,75].

Transmural differences in $I_{CaL}$ density were reduced compared to ToR-ORd, given experimental data in dog showing little difference in functional $I_{CaL}$ measurements across the wall [76]. Now, epicardium includes a 2.5% increase over endocardium (rather than the 20% in ToR-ORd), and midmyocardium includes a 10% increase over endocardium (rather than 100% in ToR-ORd). The midmyocardial versions of the original ORd and ToR-ORd models show EADs possibly too eagerly due to the strongly enhanced $I_{CaL}$, which is no longer the case in T-World.

Changes in equations compared to ToR-ORd (and/or the Doste et al. formulation of PKA effects on $I_{CaL}$) are given below:

$$d_\infty = \left(1.0763 \cdot e^{-1.007 \cdot e^{-0.0829 \cdot V + 3.62483}} \text{for } V \leq 31.4978 \,\middle|\, 1 \text{ otherwise}\right)$$

$$\tau_d = 1.5 + \frac{1}{e^{-0.05\cdot(V+6)} + e^{0.09\cdot(V+14)}}$$

$$\tau_{f,fast} = 6.171108886816713 + \frac{1}{0.00126 \cdot e^{\frac{-(V+26.63596)}{9.69961}} + 0.00126 \cdot e^{\frac{(V+26.63596)}{9.69961}}}$$

$$\tau_{f,slow} = 2719.22489 + \frac{1}{7.19411E\text{-}5 \cdot e^{\frac{-(V+5.74631)}{10.8769}} + 7.19411E\text{-}5 \cdot e^{\frac{-(V+5.74631)}{16.31535}}}$$

$$A_{f,fast} = 0.52477$$

$$\tau_{f,Ca,fast} = 13.50673 + \frac{1}{0.1542 \cdot e^{\frac{-(V-1.31611)}{11.3396}} + 0.1542 \cdot e^{\frac{(V-1.31611)}{11.3396}}}$$

$$\tau_{f,Ca,slow} = 177.95813 + \frac{1}{4.73955E\text{-}4 \cdot e^{\frac{-(V+0.79049)}{0.81777}} + 4.73955E\text{-}4 \cdot e^{\frac{-(V+2.4074)}{1.90812}}}$$

$$A_{f,Ca,fast} = 0.3 + \frac{0.6}{1 + e^{\frac{V-9.2424\acute{y}}{27.96201}}}$$

$$\tau_{jca} = 66$$

$$jca_\infty = \frac{1}{1 + e^{-\frac{V+17.66945}{3.21501}}}$$

$$K_{m,n} = 0.00222$$

$$K_{+2,n} = 957.85903$$

$$K_{-2,n} = jca \cdot 0.84191$$

$$\alpha_{n,dyad} = \frac{1}{\frac{K_{+2,n}}{K_{-2,n}} + \left(\frac{K_{m,n}}{[Ca^{2+}]_{dyad}}\right)^{3.80763}}$$

$$\alpha_{n,sl} = \frac{1}{\frac{K_{+2,n}}{K_{-2,n}} + \left(\frac{K_{m,n}}{[Ca^{2+}]_{sl}}\right)^{3.80763}}$$

(here, ',dyad' indicates dyadic variables, and ',sl' indicates subsarcolemmal variables)

Steady-state value, tau, and derivative of the new calcium-dependent inactivation (CDI) gate $f_{Ca,CDI}$ (which is combined with the activation gate by multiplication) for compartment X (dyad or subsarcolemmal) are as follows:

$$f_{\infty Ca,CDI,X} = 1 - \frac{1}{1 + \left(\frac{1.86532 \cdot [Ca^{2+}]_X}{0.032}\right)}$$

$$\tau_{f,Ca,CDI,X} = 1.0967 + (1 - f_{\infty Ca,CDI,X}) \cdot 141.4299$$

$$r_{recovery} = 0.02313$$

$$\frac{df_{Ca,CDI,X}}{dt} = -f_{Ca,CDI,X} \cdot \frac{f_{\infty Ca,CDI,X}}{\tau_{f,Ca,CDI,X}} + (1 - f_{Ca,CDI,X}) \cdot r_{recovery}$$

We have updated the driving force calculation based on our observation that the way driving force is typically calculated is problematic for ionic currents located in very small compartments such as dyads. Specifically, we believe it is problematic to use the gradient of extracellular space versus local, dyadic calcium concentrations. Both equations commonly used to describe the driving force (based on the difference from the Nernst potential, $V-E_{ion}$, or the Goldman-Hodgkin-Katz flux equation) are based on the assumption of well-mixed solutions on either side of the membrane. While this holds approximately for sodium, potassium, or chloride ions, the assumption clearly does not hold for calcium, the concentration of which changes by orders of magnitude in the dyad throughout the excitation-contraction coupling cycle. During peak release of calcium from the SR, local concentrations can reach ca. 100 µM, while bulk cytosol calcium concentrations will be in the order of low hundreds of nM at that time point. The actual physical driving force for dyadic channels at such a time point depends on the intracellular gradient of calcium (high average concentration in the dyad, decreasing with distance from the dyad). In the cell model, this gradient is discretized into two concentrations: dyadic and non-dyadic, using only the dyadic for the driving force. The exact nature of such a discretization becomes surprisingly important. If we use a small dyad model this leads to a very high local calcium concentration in the dyad. Conversely, if we work with a somewhat larger dyad, the average dyadic concentration will be lower. As a result, by changing the size of the modelled dyad, we are changing the average dyadic concentration markedly, and, by extension, we change the driving force markedly. At the cellular level, a small dyad with high local concentrations can then produce unphysiological $I_{CaL}$ reversal (corresponding to calcium efflux). However, the driving force should principally be determined by the true gradient, not a semi-arbitrary modelling choice. Based on discussions with experts in ionic fluxes across membranes (Prof. Dirk Gillespie from Rush University, Prof. Dezso Boda from University of Pannonia, Prof. Pavel Jungwirth from IOCB in Prague, and Dr. Geir Halnes from University of Oslo), we concluded that the driving force formulation for not-well-mixed solutions should also incorporate non-dyadic calcium concentrations.

Since the exact solution to this issue is currently unknown, different model families take different approaches to this apparent problem. The ToR-ORd model and its predecessors use a very large dyadic space (2% of the cell, like the whole subsarcolemmal space in Shannon-like models). This makes local concentrations in the dyad relatively low, and after the improvements in driving force formulation introduced in ToR-ORd, this leads to a well-shaped $I_{CaL}$, which does not tend to manifest current reversal. However, the 2% volume is far higher than best available estimates based on imaging data used in the Shannon et al. model and its successors, which is 0.0539%. One of the consequences is that it is extremely challenging to introduce genuinely calcium-sensitive calcium release in the ToR-ORd family, as the calcium influx through $I_{CaL}$ is a mere trickle in the vast dyad. Having RyR respond to the resulting minor calcium elevation produces a hypersensitive phenotype which is then prone to spontaneous release. On the other hand, the most up-to-date model from the Shannon-like family, Morotti2021 [3], made the $I_{CaL}$ driving force insensitive to dyadic calcium, using a constant in its place. This avoids the problem of current reversal, but makes the model insensitive to genuine changes inside the cell. The TP06 model [77] then employs much lower ionic activity inside the cell than outside (which does not seem to be theory-supported) and a shift in membrane potential used in the driving force by 15 mV (we could not find the origin of this shift, which could be a phenomenological way of avoiding driving force problems and current reversal).

In the absence of a clear answer on how the local and non-local calcium concentrations should be weighted to produce the driving force, we use a "one-compartment-further" approach. Thus, subsarcolemmal calcium concentration is used when calculating the driving force of dyadic $I_{CaL}$, and myoplasmic calcium concentration is used when calculating subsarcolemmal $I_{CaL}$. This non-local approach to driving force retains the calcium-sensitive nature of the driving force, but at the same time avoids problems arising from a fully local approach, which would lead to current reversal with high elevations of dyadic calcium. It produces a good data-like shape of $I_{CaL}$ traces, and we are not aware of any major weakness of this way of modelling driving force.

We note, that while the driving force is and should be partly non-local, processes like CDI that are local still use local concentration – the non-local "one-compartment-further" approach is used only when calculating the driving force.

In addition, ionic activity coefficients used in the driving force calculation now correctly use base of power 10, rather than *e*, such as the following for ionic species X and dyadic compartment:

$$\gamma_{X,dyad} = 10^{-A \cdot z_X^2 \cdot \left(\frac{\sqrt{I}}{1+\sqrt{I}} - 0.3 \cdot I\right)}$$

#### Transient outward current ($I_{to}$)

Baseline $I_{to}$ was formulated as in the Grandi 2010 model [2]. The conductance of fast and slow component of $I_{to}$ ($I_{to,f}$ and $I_{to,s}$) for different cell types is given in **Table S4.**

***Table S4. Conductance of $I_{to}$ components across the ventricular wall.***

| Cell type | $G_{to,f}$ | $G_{to,s}$ |
|---|---|---|
| Endocardial | 0.01276 | 0.0721 |
| Midmyocardial | 0.14928 | 0.04632 |
| Epicardial | 0.29856 | 0.02036 |

We have additionally incorporated the effect of CaMKII on inactivation of $I_{to}$, creating separate populations of phosphorylated and non-phosphorylated channels. CaMKII-phosphorylated channels have their activation shifted by 10 mV in the depolarizing direction:

$$x_{to,\infty} = \frac{1}{1 + e^{\frac{-(V-19-10)}{13}}}$$

To represent modulation of $I_{to}$ inactivation by CaMKII, we used an identical approach to that of O'Hara et al. [5], which involves multiplying time constants of inactivation of $I_{to,s}$ and $I_{to,f}$ by the product of $\delta_{CaMK,develop}$ and $\delta_{CaMK,recover}$, defined as follows:

$$\delta_{CaMK,develop} = 1.354 + \frac{10^{-4}}{e^{\frac{V-167.4}{15.89}} + e^{\frac{-(V-12.23)}{0.2154}}}$$

$$\delta_{CaMK,recover} = 1 - \frac{0.5}{1 + e^{\frac{V+70}{20}}}$$

#### Rapid delayed rectifier current ($I_{Kr}$)

$I_{Kr}$ is based on the ToR-ORd formulation, with the $\beta_i$ parameter reduced by 30% to improve restitution properties. Similar to the best available estimate of transmural changes [5], $G_{Kr}$ is scaled by 1.25 in epicardium vs endocardium, and by 0.7 in midmyocardium versus endocardium.

#### Slow delayed rectifier current ($I_{Ks}$)

Similarly to the Morotti2021 model, we have utilized the framework of calcium-sensitive formulation of $I_{Ks}$ by Bartos et al. [78] as a starting point, with the following changes compared to the Morotti2021 implementation. First, the default multiplier of the current (the variable 'gks_factor_SA' in the code) changed from 2.5 to 2.97. Second, the $I_{Ks}$ conductance is reduced down to 50% in midmyocardial cells, given canine data on transmural differences in $I_{Ks}$ density [78]. The equations describing the regulation of $I_{Ks}$ by PKA were adjusted as follows, to enable the model to reproduce the ~14-fold increase in peak $I_{Ks}$ evoked during an AP in βAR stimulated cells compared to controls [79]:

$$G_{Ks,0} = 0.01 \cdot (0.2 + 0.2 \cdot k_{PKA})$$

$$G_{Ks,max} = 0.01 \cdot (0.8 + 7 \cdot k_{PKA}),$$

where $k_{PKA}$ is the effective fraction of phosphorylated $I_{Ks}$. Subsequently, $G_{Ks}$ for the dyad and subsarcolemmal populations of $I_{Ks}$ are defined as follows:

$$G_{Ks,dyad} = G_{Ks,0} + \frac{G_{Ks,max} - G_{Ks,0}}{1 + \left(\frac{150E\text{-}6}{[Ca^{2+}]_{dyad}}\right)^{1.3}}$$

$$G_{Ks,sl} = G_{Ks,0} + \frac{G_{Ks,max} - G_{Ks,0}}{1 + \left(\frac{150E\text{-}6}{[Ca^{2+}]_{sl}}\right)^{1.3}}$$

#### Inward rectifier current ($I_{K1}$)

We employed the identical $I_{K1}$ formulation as in ToR-ORd, based on the work of Carro et al. [80]. Epicardial cells have 1.1fold conductance of endocardial ones, with midmyocardial cells showing 1.3fold conductance over the endocardium.

#### Sodium-calcium exchanger ($I_{NaCa}$)

The formulation of the sodium-calcium exchanger (NCX) was reused from the ToR-ORd model with modification to better represent conditions of sodium overload. We noticed that the model correctly shows increased influx and reduced efflux of calcium when intracellular sodium is high in most conditions, except at diastolic potentials and with resting levels of calcium. In that condition, the original model would paradoxically show increased efflux of calcium in the setting of sodium overload, complicating simulations of sodium-overloaded cells (e.g. with heavy $I_{NaK}$ inhibition such as following ouabain exposure). While sodium overload should translate into calcium overload due to more calcium retention via NCX, this was not captured well by the model given the increased calcium efflux during diastole. The following parametric changes were applied to avoid this problem while preserving the original model functionality:

$$kna1 = 11.9712$$

$$kna2 = 2.76$$

$$kna3 = 88.767$$

$$kassym = 19.4258$$

$$wna = 3.2978E04$$

$$wca = 5.1756E04$$

$$wnaca = 2.7763E03$$

$$kcaon = 3.4164E06$$

$$kcaoff = 3.8532E03$$

$$qna = 0.6718$$

$$qca = 0.0955$$

In addition, the membrane potential in the equations for *hca* and *hna* was shifted by -8.3117.

The fraction of NCX in the dyad is 31%, which is in line with a high degree of colocalization observed experimentally [81]. Similar to $I_{CaL}$, non-local calcium concentrations are used when calcium transport is calculated (dyadic NCX uses subsarcolemmal calcium concentration and subsarcolemmal NCX uses intracellular one), with the exception of allosteric regulation, which is fully local, and uses concentrations in the same compartment.

Epicardial cells have 1.1fold NCX of endocardium, and midmyocardial cells have 1.4fold NCX over endocardium.

#### Sodium-potassium pump ($I_{NaK}$)

We noticed that the initially considered $I_{NaK}$ formulation from the ToR-ORd model (which utilized a formulation from the ORd model by O'Hara et al. [5]) is problematic with regards to responses to changes in extracellular potassium. Reduction in extracellular potassium concentration is known to reduce the activity of the sodium-potassium pump [82]. The ToR-ORd/ORd model of $I_{NaK}$ reproduces this phenomenon when unphysiological ionic concentrations (also used in the experiment) are employed (**Figure S3A,B**); however, it does not do so when physiological levels used in actual cell model simulations are used (**Figure S3C**).

As such, we instead used and adjusted a formulation originally based on the Grandi et al. 2010 model [2], which represents the reduction in the pump activity with reduced extracellular potassium in a robust way. We changed the equation for $f_{NaK}$ to:

$$f_{NaK} = 0.75 + \left(0.00375 - 0.001 \cdot \frac{140 - Na_o}{50}\right) \cdot V$$

With this formulation, the voltage-dependence of $I_{NaK}$ is linear, compared to sublinear in Grandi et al. [2], based on the observation that the sublinear relationship is obtained with 50 mM intracellular sodium, whereas a physiological concentration of 8 mM yields a relatively linear relationship (**Figure S3D**). At the same time, the updated model recapitulates the observation that extracellular sodium alters the slope of the voltage-dependence of $I_{NaK}$ (**Figure S3E,F**).

$I_{NaK}$ activation via phospholemman phosphorylation by βARS is represented by the inclusion of a PKA-activated $I_{NaK}$ population which has increased affinity for intracellular sodium, similar to experimental measurements [83] ($Km_{Nai}$ = 11 for normal channels in the model, $Km_{Nai,PKA}$ = 8.4615 mM).

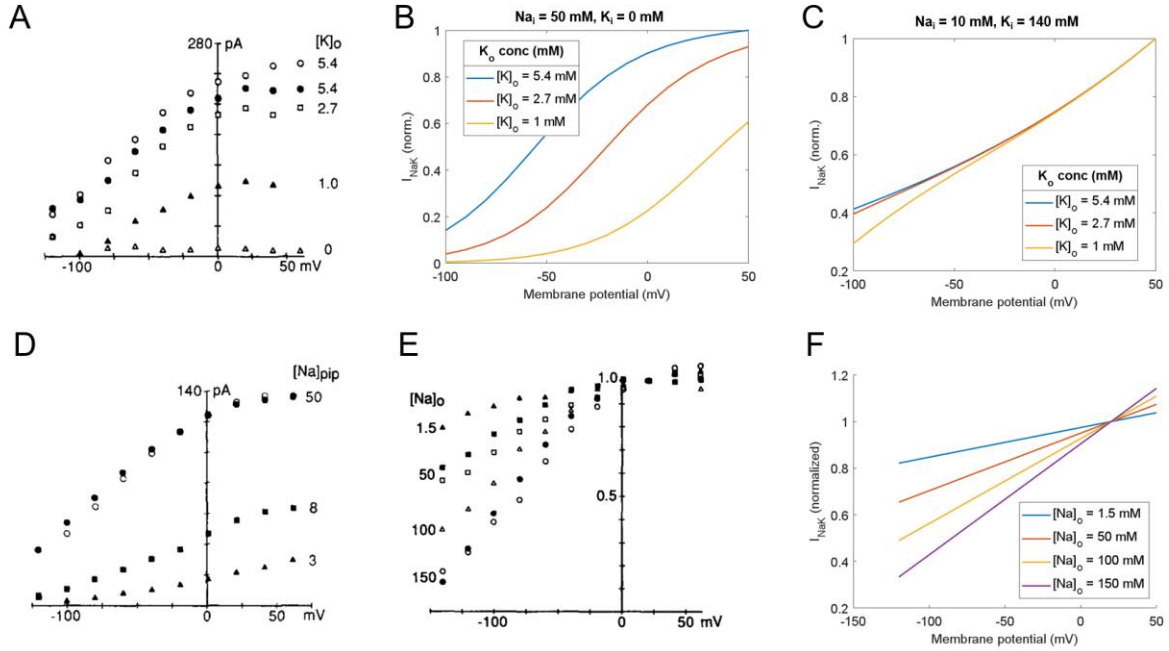

***Figure S3 Sodium-potassium pump properties**. **A)** Dependence of $I_{NaK}$ on extracellular potassium, figure reproduced from Nakao & Gadsby 1989 [84], as permitted by the CC-BY-NC-SA licence. **B)** Simulated $I_{NaK}$ using the ToR-ORd/ORd formulation, using high intracellular sodium and zero intracellular potassium, as used in the patch pipette in panel A. **C)** Demonstration of loss of potassium-dependence of $I_{NaK}$ when physiological intracellular concentrations (10 mM sodium and 140 mM potassium) are used in the same model. **D)** Demonstration of sublinear dependence of $I_{NaK}$ on membrane potential is at unphysiologically high intracellular sodium, but near-linear dependence at physiological levels. Figure reproduced from the work of Nakao & Gadsby 1989. **E)** Voltage-dependence of $I_{NaK}$ for varying levels of extracellular sodium, again reproduced from Nakao & Gadsby 1989. **F)** Simulation of the new $I_{NaK}$ model, corresponding to data in panel E.*

#### Chloride currents ($I_{(Ca)Cl}$, $I_{Clb}$)

The calcium-sensitive chloride current $I_{(Ca)Cl}$ and the background chloride current $I_{Clb}$ were formulated as in the Grandi 2010 model [2], with an update to the conductance.

#### Background currents ($I_{Nab}$, $I_{Cab}$, $I_{Kb}$)

Background sodium and calcium currents ($I_{Nab}$, $I_{Cab}$) were formulated as in the Grandi 2010 model [2] and the background potassium current ($I_{Kb}$) as in ToR-ORd, all with updated conductances. No transmural gradient of those channels is present.

#### Stimulus current ($I_{stim}$)

A conservative potassium stimulus is used as in ToR-ORd [72], as suggested by Hund et al. [4].

#### Calcium release from the SR ($J_{rel}$, $J_{leak}$)

The representation of calcium release from the SR ($J_{rel}$) combines the approaches from ToR-ORd and Shannon-like models of calcium-induced calcium release [1,72]. Shannon-like models are mechanistically more realistic, with $J_{rel}$ being triggered by dyadic calcium, rather than $I_{CaL}$ (which is the case in ToR-ORd), but they suffer from the issue of late-peaking release (see **Figure S12**). The mechanistic importance is not just for its own sake, but also for enabling the formation of delayed afterdepolarizations (DADs), which do not occur spontaneously in ToR-ORd. We therefore designed a hybrid scheme, where a very small population of RyR (ca. 4%) is directly coupled to $I_{CaL}$ (based on a modified ToR-ORd formulation), representing the most tightly coupled $I_{CaL}$-RyR clusters. This provides an early source of calcium influx, which then helps activate at the right time the much larger population of RyR that are modelled using a modified Shannon-like formulation. The latter has been modified predominantly to support the earlier time to peak of calcium release, and to enable calcium-driven alternans. In addition, passive leak from the SR to the dyad is included in the model. The equations for the distinct components are given below.

##### *$I_{CaL}$-activated RyR flux ($J_{rel,ICaLdep}$)*

The formulation is a reparametrized and updated version of $J_{rel}$ from ToR-ORd with a small maximum flux and two additional inactivation gates to keep its duration short and thereby keeping the overall amount of calcium released in this way low. The $J_{rel,ICaLdep}$ flux over time is shown in **Figure S4A**, demonstrating a minor role in the overall release. We furthermore confirmed that calcium-driven alternans manifests in T-World even when the $J_{rel,ICaLdep}$ component of release is turned off (**Figure S4B**). I.e., alternans arises from the "main" calcium-sensitive component of calcium release and is not driven by the $I_{CaL}$-sensitive component.

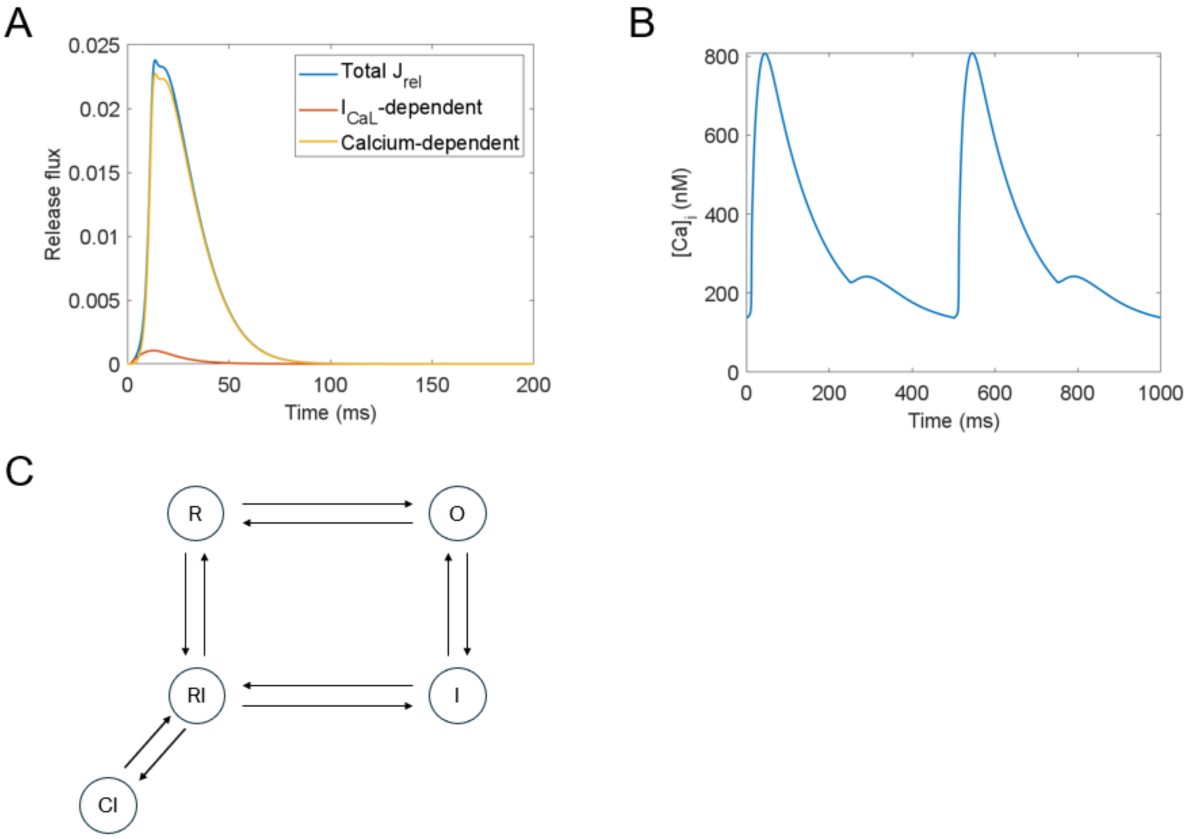

***Figure S4. Modelling ryanodine receptors. A)*** *Flux through RyR: total and its two sub components ($I_{CaL}$-dependent and calcium-dependent), demonstrating that the calcium release from the SR is largely calcium-sensitive (the $I_{CaL}$-dependent component is only 4.2% of total release with regards to peak and 3.8% with regards to integral).* ***B)*** *Illustration of presence of calcium-driven alternans at basic cycle length of 260 ms even when the $I_{CaL}$-dependent component of release is turned off.* ***C)*** *A diagram of the calcium-sensitive RyR Markov model.*

The derivative of the activation gate of the directly coupled RyR ($J_{rel,ICaLdep,act}$) is given by:

$$I_{Ca,junc,sigmoided} = 1 - \frac{1}{1 + \left(\frac{|I_{CaL,dyadic}|}{0.45}\right)^{4.5}}$$

$$J_{rel,ICaLdep,act,\infty} = 15.5959 \cdot \frac{I_{Ca,junc,sigmoided}}{1 + \left(\frac{0.95271}{[Ca]_{SR}}\right)^{7.72672}}$$

$$\tau_{rel} = \max\left(\frac{12.4767}{1 + \frac{0.0123}{[Ca]_{SR}}}, 0.001\right)$$

$$\frac{dJ_{rel,ICaLdep,act}}{dt} = \frac{J_{rel,ICaLdep,act,\infty} - J_{rel,ICaLdep,act}}{\tau_{rel}}$$

The derivatives for the two inactivation gates ($J_{rel,ICaLdep,f1}$, $J_{rel,ICaLdep,f2}$) are calculated as follows:

$$\frac{dJ_{rel,ICaLdep,f1}}{dt} = \frac{\frac{1}{1+\frac{I_{Ca,junc,sigmoided}}{0.001}} - J_{rel,ICaLdep,f1}}{64.11202}$$

$$\frac{dJ_{rel,ICaLdep,f2}}{dt} = \frac{\frac{1}{1+\frac{I_{Ca,junc,sigmoided}}{0.0006}} - J_{rel,ICaLdep,f2}}{119.48978}$$

The total $I_{CaL}$-sensitive RyR flux is calculated as follows:

$$J_{rel,ICaLdep} = 0.00174 \cdot J_{rel,ICaLdep,act} \cdot J_{rel,ICaLdep,f1} \cdot J_{rel,ICaLdep,f2}$$

##### *Calcium-activated RyR flux ($J_{rel,ICaLdep}$)*

The calcium-sensitive component of the SR release of calcium via RyR is based on an extensively reparametrized 4-state square Markov model by Shannon et al model [1]. This was extended with a single "calcium-inactivation" state (**Figure S4C**), providing an additional partial source of RyR refractoriness. In particular, this helps the model to reproduce at rapid pacing the combination of high SR load, but somewhat limited RyR release (indicated by the limited SR depletion e.g. in [24]), less than might be expected from the load-release relationship. The equations determining the transitions between states are as follows:

**R → O, RI → I:**

$$k_{Ca,SR} = 15 - \frac{14}{1+\left(\frac{0.75385}{[Ca]_{SR}}\right)^{5.09473}}$$

$$k_{o,SRCa} = \frac{23.87221}{k_{Ca,SR}}$$

$$transition_{R\to O\&RI\to I} = 0.52967 \cdot k_{o,SRCa} \cdot \left([Ca]_{dyad}\right)^{2.06273}$$

**O → R, I → RI:**

$$transition_{O\to R\&I\to RI} = 0.16219$$

**O→I, R→RI:**

$$k_{i,SRCa} = 0.39871 \cdot k_{Ca,SR}$$

$$transition_{O\to I\&R\to RI} = 0.94428 \cdot k_{i,SRCa} \cdot \left([Ca]_{dyad}\right)^{0.68655}$$

**I → O, RI → R:**

$$transition_{I\to O\&RI\to R} = 0.04311$$

**RI → CI:**

$$transition_{RI\to CI} = 3.0232E\text{-}4 \cdot \left(0.93249 + \frac{29.2005}{1+\left(\frac{0.001}{[Ca]_{dyad}}\right)^{5.93447}}\right)$$

**CI → RI:**

$$transition_{CI\to RI} = 0.00248$$

The overall calcium flux through the calcium-sensitive, non-phosphorylated RyR is as follows:

$$J_{rel,Cadep,NP} = 26.6 \cdot O \cdot ([Ca]_{SR} - [Ca]_{junc}),$$

where O is the fraction of RyRs in the open state.

A separate population is simulated for RyR phosphorylated by CaMKII. The equations are identical, except the transition from R→O (and symmetrically RI→I) is increased by 50% to represent the increased open probability of RyR following CaMKII phosphorylation [85]. Using this, the flux through calcium-sensitive phosphorylated RyR ($J_{rel,Cadep,P}$) is calculated analogously.

##### *Combined $J_{rel}$*

The combined flux is calculated as follows:

$$J_{rel} = (1 - f_{RyRP,CaMKII}) \cdot (J_{rel,Cadep,NP} + J_{rel,ICaLdep}) + f_{RyRP,CaMKII} \cdot (J_{rel,Cadep,P} + J_{rel,ICaLdep})$$

I.e., it is a weighted sum of CaMKII-phosphorylated and unphosphorylated RyR flux, weighted by the fraction of RyR phosphorylated by CaMKII.

##### *RyR leak ($J_{SR,leak}$)*

The leak between the SR and the dyads is represented similarly to Shannon et al. [1], extended to account for CaMKII-dependent potentiation of SR calcium leak [85] and super-linear load-dependence [86].

$$multiplier_{CaMKII} = 1 + 2 \cdot f_{RyRP,CaMKII}$$

$$multiplier_{nonlinear} = 0.2144 \cdot e^{1.83 \cdot [Ca]_{SR}}$$

$$J_{SRleak} = 1.59306E\text{-}6 \cdot ([Ca]_{SR} - [Ca]_{junc}) \cdot multiplier_{CaMKII} \cdot multiplier_{nonlinear}$$

#### Calcium reuptake to the SR ($J_{up}$)

The formulation of SERCA pumps was mainly based on the Shannon et al. formulation [1], with the following parametric changes (in addition to $V_{max,SERCA}$, listed in **Table X**): $K_{mr}$ = 2.31442, $K_{mf}$ = 0.30672E-03, H (Hill coefficient) = 1.02809. The maximum pumping rate $V_{max,SERCA}$ in the epicardium is increased by 20% versus (baseline) endocardium, following the report of an increased epicardial SERCA protein expression by Laurita et al. [87].

We have additionally incorporated SERCA potentiation by phospholamban phosphorylation (possible via the CaMKII and PKA pathways). This is represented by a separate population of SERCA pumps with a greater affinity for cytosolic calcium (half $K_{mf}$ versus unphosphorylated, $K_{mf,phosphorylated}$ = 1.5336E-04). The fraction of phosphorylated SERCA pumps is calculated as the sum of fraction of pumps phosphorylated by CaMKII and those phosphorylated by PKA, minus the product of those fractions.

Compared to ToR-ORd, PLB/SERCA are only marginally phosphorylated by CaMKII, reflecting experimental observations by Huke and Bers, who observed minimal phosphorylation by CaMKII at this site [88]. Nevertheless, major potentiation of SERCA pumping rate was observed experimentally at rapid versus slow pacing [89]. We represent this phenomenon phenomenologically using a separate set of equations similar to how CaMKII is activated by calcium, which are however distinct from CaMKII phosphorylation itself. Therefore, when CaMKII overexpression or hyperactivity are simulated in future studies, this calcium-based activation will be unaffected. We note that in normal conditions, this calcium-based activation is functionally similar to ToR-ORd, where SERCA/PLB can be relatively highly phosphorylated by CaMKII. The degree of activation of this mechanism is determined as follows (reusing some of the CaMKII-related named constants, but not state variables):

$$\alpha_{SERCA} = 0.05$$

$$bound_{SERCA} = CaMK_0 \cdot \frac{1 - SERCA_{casig,trap}}{1 + \frac{Km_{CaMK,Ca}}{[Ca]_{dyad}}}$$

$$SERCA_{casig,act} = bound_{SERCA} \cdot SERCA_{casig,trap}$$

$$\frac{dSERCA_{casig,trap}}{dt} = \alpha_{SERCA} \cdot bound_{SERCA} \cdot SERCA_{casig,act} - \beta \cdot SERCA_{casig,trap} \cdot (0.1 + 0.9) \cdot \frac{PP1_{tot}}{0.1371}$$

The $SERCA_{casig,act}$ variable is subsequently used in the equation describing the multiplier applied to $V_{max,SERCA}$:

$$V_{max,multiplier} = 1 + \frac{1.111423947401174}{1 + \left(\frac{0.4}{SERCA_{casig,act}}\right)^2}$$

A similar principle can likely be used in the future to rescue the relatively negative frequency-force relationship in models utilizing the Soltis-Saucerman CaMKII model, accelerating calcium reuptake at rapid pacing. However, this will also require a cell model reproducing the positive relationship between SERCA activity and calcium transient amplitude (as T-World does, see **Figure 2F** of the main manuscript).

#### Sarcolemmal calcium pump ($p_{Ca}$)

The sarcolemmal calcium ATPase was formulated as in the Grandi 2010 model [2], with an update to the conductance.

#### Buffering and ion diffusion between compartments

Cell compartmentalization and buffering are based on the Shannon et al. framework [1]. One exception is the buffering of calcium by Troponin C, which is replaced by the Land contraction model formulation (see next section). The following parametric changes were made:

$$Bmax_{SR} = 17.85854E\text{-}3$$

$$Bmax_{SLlow,sl} = 33.923E\text{-}3 \cdot \frac{V_{myo}}{V_{sl}}$$

$$Bmax_{SLlow,dyad} = 4.89983E\text{-}4 \cdot \frac{V_{myo}}{V_{sl}}$$

$$Bmax_{SLhigh,sl} = 12.15423E\text{-}3 \cdot \frac{V_{myo}}{V_{sl}}$$

$$Bmax_{SLhigh,dyad} = 1.75755E\text{-}4 \cdot \frac{V_{myo}}{V_{sl}}$$

$$Bmax_{CSQN} = 136.55214E\text{-}3 \cdot \frac{V_{myo}}{V_{SR}}$$

$V_{myo}$, $V_{sl}$, $V_{junc}$ and $V_{SR}$ indicate the volumes of myoplasmic, subsarcolemmal, dyadic (junctional), and SR compartments.

Diffusion coefficients of calcium between compartments are:

$$J_{Ca,juncsl} = \frac{1}{3.06685E12}$$

$$J_{Ca,slmyo} = \frac{1}{0.74556E11}$$

#### Contraction modelling

When including the representation of contraction in T-World, we were initially faced with the choice of whether to take as a starting point the model by Negroni et al. [90] or the Land et al. [91], the two currently leading published formulations.

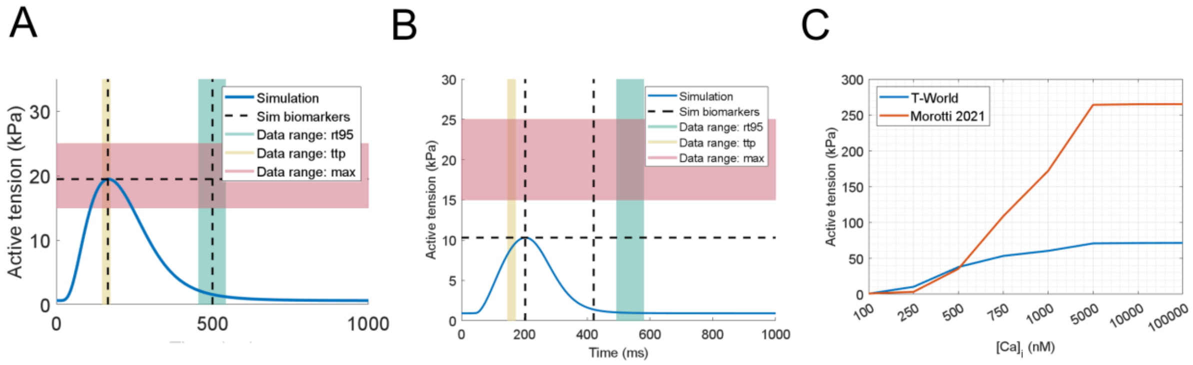

***Figure S5. Contraction properties of Land and Negroni-based models. A)*** *T-World contraction with biomarker ranges shown as in Figure 1 of the main manuscript.* ***B)*** *Corresponding simulation of contraction in the Morotti2021 model (please note that the model was not recalibrated for the experimental ranges like T-World was, i.e., the lower agreement does not necessarily imply an overall worse model).* ***C)*** *Comparison of steady-state active tension between T-World and Morotti2021 models evoked by a range of calcium concentrations. The end points of the curves reflect peak force achievable by the model.*

In principle, starting with the Negroni model would be simpler, given that it is already integrated in the calcium handling framework that T-World is based on (and it provides contraction representation e.g. in the Morotti2021 model [3], referred to extensively in this project). Both Land and Negroni models are capable of generally data-like force development when integrated in cardiomyocyte models (**Figure S5A,B**). The key factor in our decision was the correspondence between the models and the observation that the twitch force during normal activation by an action potential reaches ca. 25-70% of peak contraction force possible [92,93]. While the T-World model with the Land model is in agreement with this (ca 28% twitch force versus peak contraction shown in **Figure S5A** vs **Figure S5C**), this does not hold for the Negroni model embedded in Morotti2021, which reaches only around 4% of peak force during a normal twitch (**Figure S5B** vs **S5C**). This appears to be a result of combination of slow calcium binding to the contractile apparatus, and then additionally slow force development following calcium binding. Given the extensive reparameterization that would be needed, we decided to adopt the Land model as a baseline, with the following parametric changes:

$$TRPN_n = 1.65 \text{ (based on [94])}$$

$$Ktm_{unblock} = 0.02626$$

$$k_{off} = 0.07854$$

$$ca_{50} = 0.7645$$

$$nu = 10.15996$$

$$mu = 3.94046$$

$$T_{ref} = 80$$

The contraction model was coupled to the cellular model via the Troponin C calcium buffering, where the original formulation was replaced by the buffering carried out by the Land model.

We have additionally incorporated the effect of βAR stimulation on the contractile apparatus. Similarly to Negroni et al. [90], we separated the effects into two sites: Troponin I and Myosin binding protein C.

Troponin I phosphorylation is known to reduce the affinity of the contractile apparatus to calcium [95,96]. This is represented in T-World by scaling the $ca_{50}$ parameter (midpoint of calcium-force relationship curve) by the following expression:

$$ca50_{scaling} = 1.45 - \frac{0.45 \cdot f_{TnI,PKA}}{1-0.0031},$$

where $f_{TnI,PKA}$ is the effective fraction of Troponin I phosphorylated by PKA.

Similar to Negroni et al. the acceleration of contraction kinetics was assigned to the Myosin binding protein C phosphorylation site, with the scaling factor (applied to the $k_{ws}$ and $xb_{uw}$ parameter) being as follows:

$$PKA_{XB,acceleration} = 1 + 0.5 \cdot f_{MyBPC,PKA},$$

where $f_{MyBPC,PKA}$ is the effective fraction of Myosin binding protein C phosphorylated by PKA.

In addition, peak force $T_{ref}$ is scaled by $1+0.26 \cdot f_{MyBPC,PKA}$, reflecting the ca. 20-30% increase in maximal force observed following PKA activation [97].

During the development, we noticed a possible issue in the coupling of the Land model to our prior model ToR-ORd, linked to the buffering of calcium by the Land model being slower than the baseline buffering by Troponin C. As a result, the released calcium stays longer in the cytosol before being buffered, and in the setting of enhanced calcium release (e.g. in midmyocardial cells or when $I_{CaL}$ is otherwise increased), a non-data-like two-peak CaT may emerge (see e.g. Fig 4A, midmyocardial cell, in [11]). However, this issue is not present in T-World, which retains normal-looking Ca transient even in the setting of strongly enhanced calcium release (e.g. with βARS, see **Figure 3** of the main manuscript). This results from the inclusion of sarcoplasmic reticulum buffering sites in the myoplasmic compartment in T-World (similarly to Shannon-based models, but unlike ToR-ORd and its predecessors). These sites provide sufficiently strong buffering capacity to prevent the aberrant CaT morphology, even when the Troponin C buffering is switched from the original to Land-based formulation.

#### CaMKII signalling

We decided to use a relatively simple formulation of CaMKII activation by calcium, similar to ToR-ORd [72]. CaMKII targets in T-World comprise $I_{Na}$, $I_{NaL}$, $I_{to}$, $I_{CaL}$, RyR, and SERCA pumps (via PLB). We also considered using the more complex model by Soltis-Saucerman [98], but ultimately decided against it, given a) a much higher complexity increasing simulation time considerably, b) a problematic effect of CaMKII activation on cellular physiology. Specifically, we observed that CaMKII activation at rapid pacing leads to reduction in SR calcium content and mostly negative rate-dependence of calcium transient (as seen also in Supplementary note 3, Morotti2021 model, which utilizes the Soltis-Saucerman model).

We nevertheless took inspiration from the Soltis-Saucerman model, reflecting the prediction that different sites in the cell are likely phosphorylated to different extent at different heart rates (unlike ToR-ORd or ORd CaMKII representation, which phosphorylates all targets equally). This is achieved by using the following equations when calculating phosphorylation levels:

$$CaMK_{\infty phos,CaMKII,ICaL} = \frac{CaMKII_{active}}{CaMKII_{active} + 0.35}$$

$$CaMK_{\infty phos,CaMKII,RyR} = \frac{CaMKII_{active}}{CaMKII_{active} + 1}$$

$$CaMK_{\infty phos,CaMKII,PLB} = \frac{CaMKII_{active}}{CaMKII_{active} + 10}$$

$$\frac{df_{CaMKII,ICaL}}{dt} = \frac{CaMK_{\infty phos,CaMKII,ICaL} - f_{CaMKII,ICaL}}{10000}$$

$$\frac{df_{CaMKII,RyR}}{dt} = \frac{CaMK_{\infty phos,CaMKII,RyR} - f_{CaMKII,RyR}}{10000}$$

$$\frac{df_{CaMKII,PLB}}{dt} = \frac{CaMK_{\infty phos,CaMKII,PLB} - f_{CaMKII,PLB}}{100000},$$

where $CaMKII_{active}$ is the fraction of active CaMKII, $f_{CaMKII,X}$ is the fraction of target X phosphorylated by CaMKII, and ∞ in subscript indicates a steady-state value.

In addition, the value of the $CaMKII_0$ parameter of the CaMKII signalling was set to 0.1, and the $Km_{CaMK,Ca}$ to 0.0075.

The resulting phosphorylation levels at different heart rates are shown in **Figure S6**. $I_{Na}$, $I_{NaL}$, and $I_{to}$ use identical phosphorylation levels as the RyR.

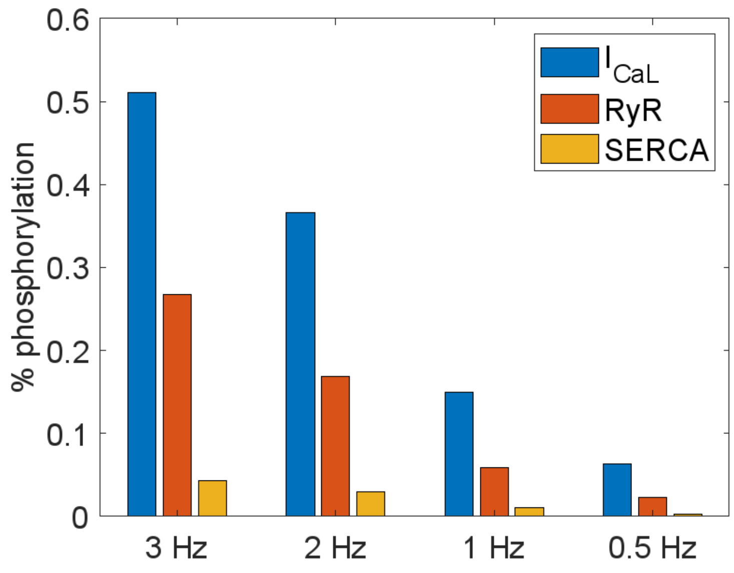

***Figure S6*. *Simulated fractional rate-dependent CaMKII activation at three different subcellular sites.*** *The phosphorylation for each target was determined as an average value throughout the last action potential out of 250 at a given stimulation rate.*

#### βAR signalling

The representation of effects of sympathetic nervous activation on cardiomyocytes via β-adrenergic (βAR) stimulation consists of two relatively separate parts. The first part consists of a signalling pathway, which calculates the effective phosphorylated fractions of Protein Kinase A (PKA) targets based on the concentration of a beta-agonist. This part is simulated based on the Doste et al. implementation of the Heijman et al. model [70]. The second part then describes how a given level of phosphorylation translates into changes in target function. We formulated these changes based on available literature, and they are described in corresponding sections above, when changes to ionic currents and fluxes are described. The targets affected by PKA phosphorylation are $I_{Na}$, $I_{CaL}$, $I_{Ks}$, phospholamban (regulating SERCA pumps and thus $J_{up}$), phospholemman (regulating the NKA pump and thus $I_{NaK}$), and the contractile apparatus (at two sites, Troponin I and Myosin binding protein C).

The signalling pathway may be turned off, with the user directly supplying the phosphorylated fractions of the PKA targets instead, which enables simulating βAR activation without the increased runtime resulting from simulating the signalling pathway.

#### Sex differences

The inclusion of sex as a biological variable is becoming an important factor for in-silico investigations due to the historical under-representation of the female sex in basic research [99] and clinical studies [100]. We include sex-specific variants of the electrophysiological model through variations of the conductance of specific ionic channels which have been identified with significantly different expression in human genomics studies [101] and validate these variant models through comparison with experimentally observed differences in male and female adult electrophysiology. Using a similar approach to Holmes et al. [102], Yang and Clancy [56], Peirlinck [103], we apply scaling factors to ionic channel conductance representative of the published differences in expression of corresponding proteins in un-diseased adult male and female human cells (**Table S5**).

*Table S5. T-World sex-specific conductance modifications and corresponding evidence.*

| Conductance | Male Scaling Factor | Female Scaling Factor | Evidence |
|---|---|---|---|
| $G_{NaCa}$ | 0.9325 | 1.0724 | Upregulated in female rabbits and rats [104]. (Furthermore upregulated via estrogen in humans [105]). |
| $G_{Kr}$ | 1.1 | 0.87 | $I_{Kr}$ subunits more expressed in males [101]. |
| $G_{Ks}$ | 1.09 | 0.905 | $I_{Ks}$ subunits more expressed in males [101]. |
| $G_{K1}$ | 1.07 | 0.92 | $I_{K1}$ subunits more expressed in males [101]. |
| $G_{pCa}$ | 0.8 | 1.28 | Higher calcium pump expression in females [101]. |
| $CMDN_{Max}$ | 0.909 | 1.1 | More calmodulin in females [101]. |

#### 3D simulations

We developed a multiscale computational framework for simulating human cardiac electrophysiology from ionic currents to electrocardiographic signals under healthy and ischemic conditions using the GPU-based solver MonoAlg3D[106]. The aim was to analyse the behaviour of the cellular model to trigger and yield ventricular arrhythmias when implemented in 3D ventricular meshes obtained from clinical MRI data. The human biventricular model was calibrated and evaluated using extensive clinical and experimental data, both under healthy and ischemic conditions, and at multiple scales including cell (ion dynamics, action potential), tissue (conduction velocity, fibre anisotropy), and organ (activation sequence, electrocardiographic signal, electrophysiological heterogeneities) levels as in[107]. Fibre orientation in the myocardium was represented using the rule-based method by Doste et al.[108] reproducing experimental findings[109].

Given the high pro-arhythmic potential of early (phase 1A) myocardial ischemia[110], we simulated acute regional ischemia in the anterior myocardial wall caused by left-anterior descending (LAD) artery occlusion to assess the cellular model's capabilities to trigger and yield pro-arrhythmic behaviours at an organ scale. Electrophysiological alterations derived from acute ischaemia, caused by hyperkalaemia, hypoxia, and acidosis, were added into the ischaemic area to model the changes in refractoriness and conduction velocity in humans[111,112]. The formulation of the ATP-dependant potassium current $IK_{(ATP)}$ [113] was incorporated to simulate hypoxia. Electrophysiological alterations in the ischaemic zone were included considering the ischaemic core zone (ICZ) and the lateral border zone (BZ), which represent an electrophysiological gradient between the ICZ and the remote myocardium[112]. The subendocardial border zone was included to account for the oxygen diffusion from blood in the ventricular cavities, reported experimentally[114].

Sinus rhythm was simulated to obtain realistic ECGs, implemented through a realistic patient-specific model of the ventricular conduction system based on previous work[115,116]. Cardiac activity at rest was simulated by sinus rhythm pacing at basic cycle length (bcl) of 800 ms. Arrhythmia was induced by a stimulation protocol with progressively shorter cycle length (2 beats at bcl of 400 ms, 2 beats at 350 ms bcl, 4 beats at 300 ms, 6 at 250 ms bcl) taking approximately 4 seconds to complete the pacing protocol. No further pacing was induced until the end of the 10-second simulation to assess arrhythmia sustenance.

### Arrhythmia studies

#### EADs

In order to evoke EADs, we simulated T-World for 100 beats at 0.25 Hz, with extracellular calcium of 2 mM, and with maximum conductance of $I_{Kr}$ reduced by 85%, based on the work of Guo et al. [27]. The last beat of the 100 was investigated for the presence of EADs.

#### DADs and stochastic DADs

To evoke a single deterministic DAD leading to triggered activity, we pre-paced T-World for 200 beats at 2.5 Hz with full β-AR stimulation and 3.25 mM extracellular calcium. This was followed by one more paced stimulus followed by several seconds of quiescence, during which the presence of DADs was recorded. To obtain a train of multiple stimuli, we used the same protocol, except 3 Hz pre-pacing for 150 beats was used. To sensitize RyR, we increased the R→O (and symmetrically RI→I) transition by 100%.

We also provide a separate implementation of T-World with stochastic DADs resulting from spontaneous overload-driven release, similar to [117]. This is provided in a separate model code function, given the use of global variables, which we found to be necessary as explained below, but which are otherwise discouraged in Matlab and we avoid using them in the baseline code. The overall idea is simple: provided that calcium in the SR is over a given threshold, there is a certain probability that a spontaneous release may occur, lasting for a parameter-determined duration. Here we used 0.9 mM as the minimum SR content, 100 ms as release duration, and the probability of spontaneous release such that on average 1 DAD occurs per 10 seconds. Cells were pre-paced for 100 beats at 2.5 Hz, using 3 mM extracellular calcium. Once a spontaneous release is determined to occur, starting at *timeStartDAD* and ending at *timeEndDAD*, with amplitude given by the parameter *amp*, the formula for the spontaneous release at time $t \in (timeStartDAD, timeEndDAD)$, which is added to the SL compartment) is:

$$J_{rel,stoch} = amp \cdot 2 \cdot \frac{\min(t - timeStartDAD, timeEndDAD - t)}{timeEndDAD - timeStartDAD}$$

I.e., it is a triangle-shape release, zero at the ends of the time interval, and with maximum at the midpoint between starting and ending time. Using such a spontaneous release function is straightforward when using a fixed-step method to solve ODEs. However, particularly for single cell simulations, it is much more efficient to use variable step solver, such as @ode15s in Matlab. In such a case, one needs to scale the probability of DAD starting at a given time step by the length of the time step (otherwise, if kept uniform, DADs would occur more frequently earlier during the beat's simulation, where time step is smaller). However, such scaling in Matlab is complicated by the fact that there is, to our knowledge, no clear way of finding the current step size. We designed an approximate scheme, which uses global variables to store the information on previous step's starting time, which can be used to calculate the current time step, as the current time is available in the ODE system code. In addition, we have furthermore limited the maximum time step to 25 ms. If left unconstrained, the solver can choose time steps over 100 ms in later phases of a beat simulation, which makes it skip a DAD that was determined to happen, which leads to under-occurrence of late DADs. Finally, we note that the current formulation of stochastic DADs with a fixed probability of spontaneous release when calcium in the SR exceeds a threshold is rather basic – future versions may modulate the probability by the SR content, and possibly also by calcium in the adjacent SL compartment.

The stochastic formulation of DADs is also a practical way of achieving subthreshold DADs as well as triggered activity in the same parameterization of a model (unlike the default T-World, which produces triggered activity). We note the fact that T-World DADs lead to action potential development, whereas they remain mostly subthreshold in the Shannon et al. model[1] is not primarily about differences in calcium handling, but about the amount of $I_{K1}$. The Shannon model has ca. fourfold $I_{K1}$ and hence it maintains the resting potential much more strongly, usually not permitting the development of DADs into triggered activity. We confirmed that when T-World has $I_{K1}$ increased four times, DADs are likewise converted to subthreshold ones (not shown).

#### Alternans

To evoke alternans, we simulated T-World for 250 beats at a range of frequencies: basic cycle lengths 240, 250, 260, ... 400, 420, 440,... 500, 600, 700, ..., 1000 ms, recording the last 4 beats. In addition to the control model, a version with 65% SERCA pump $V_{max}$ (0.65 $J_{up}$ multiplier), 150% SERCA $V_{max}$, and β-AR-stimulated versions were simulated.

#### Restitution

When measuring the slope of the restitution, we pre-paced T-World for 100 S1 beats, which were followed by a pair of stimuli S2 ms apart. S2 intervals explored ranged from S1 down to 200 ms. The default S1 interval was 1000 ms (1 Hz), with 600 and 400 used for a part of the simulation, as described in the corresponding figure. Action potential duration was measured at the level of -75 mV. Peak slope of the S1S2 restitution curve was determined as the maximum slope of diff($APD_{S2}$)./diff(diastolic interval), where APDS2 is the vector of APDs of all the S2 stimuli. Data corresponding to S2 stimuli which did not reach the threshold of 0 mV were discarded. To explore the relationship between the baseline cell APD and its peak restitution slope, we created 300 models with randomly perturbed $I_{CaL}$, $I_{Kr}$, $I_{Ks}$, $I_{NaL}$, $I_{NaK}$, $I_{NaCa}$ (each parameter scaled by a number between $e^{-0.5}$ and $e^{0.5}$; the exponent was randomly drawn from uniform distribution between -0.5 and 0.5). This was done for T-World, Morotti 2021 and TP06 models (in the case of TP06, $I_{NaL}$ was effectively not changed, as the model does not represent $I_{NaL}$).

#### Stability of arrhythmic behaviours

We assessed the stability/robustness of arrhythmic behaviours in a population of models with varied parameters, calibrated to human-derived criteria in **Table S6**. $I_{Na}$, $I_{NaL}$, $I_{to,f}$, $I_{CaL}$, $I_{K1}$, $I_{Kr}$, $I_{Ks}$, $J_{rel}$, $J_{up}$, $I_{NaCa}$, $I_{NaK}$ were varied between 67% and 150% (sampled using log-multipliers, so that x-fold reduction is equally likely as x-fold increase). 787 out of 1000 models generated in this way passed all the calibration criteria.

***Table S6. Biomarker ranges for the calibration of population of models used in the in silico trial.***

| Biomarker | Minimum | Maximum |
|---|---|---|
| AP peak | 7 mV | 55 mV |
| Resting membrane potential | -95 mV | -80 mV |
| Peak upstroke velocity | 100 V/s | 1000 V/s |
| APD90 (Action potential duration at 90% recovery) | 180 ms | 440 ms |
| APD50 (APD at 50% recovery) | 110 ms | 350 ms |
| APD40 (APD at 40% recovery) | 85 ms | 320 ms |
| 90-40 triangulation (APD90-APD40) | 50 ms | 150 ms |
| CaT duration at 90% recovery | 220 ms | 750 ms |
| CaT duration at 50% recovery | 120 ms | 420 ms |
| CaT amplitude | 200 nM | 600 nM |
| Peak of CaT | 200 nM | 1000 nM |
| Diastolic Ca | 0 nM | 400 nM |
| Peak active tension | 5 kPa | 40 kPa |
| Time to peak active tension | 120 ms | 200 ms |
| Time from peak active tension to 95% recovery | 200 ms | 600 ms |

To assess stability of EADs in the 787 chosen models, we varied together $I_{CaL}$ and $I_{Kr}$, adding those changes (via multiplication) to the parameterization of each model. For each model, we explored $I_{CaL}$ multipliers of 1, 1.01,...,1.45, concurrently with $I_{Kr}$ multipliers of 1, 0.98, ...0.1. In this way, EADs are gradually promoted through a medium-strength $I_{CaL}$ increase and a reduction in repolarization. For each model, we recorded the earliest pair of $I_{CaL}$ and $I_{Kr}$ multipliers producing EADs.

To assess the stability of deterministic DADs, we simulated each of the 787 models with fully active β-AR signalling, at extracellular calcium of 2, 2.2, ..., 6 mM. 200 beats of pre-pacing were used at 2.5 Hz, recording how much extracellular calcium each model requires to trigger DADs in a period of quiescence following the pre-pacing phase. These simulations were done separately in a baseline model and in a model with 50% increase in $J_{up}$, promoting SR overload.

To assess stability of alternans in T-World, we simulated each model with $J_{up}$ multiplier of 0.5, 0.6, ..., 1.5, at basic cycle length of 260 ms, recording the number of multipliers which yield alternans at the end of 100 beat pacing train.

To assess the stability of S1S2 restitution, we simulated the S1S2 protocol in each of the 787 models, recording the peak restitution slope (100 S1 beats followed by a S2 beat after 200,205,...,1000 ms).

### T-World applications

#### Drug safety and efficacy

We generated a population of models for the drug safety assessment study, representing variation in ionic currents and calcium handling [72,118]. We generated a population of 1000 models, with the following randomly sampled perturbations to ionic currents and fluxes versus baseline: 50-200% ($I_{Na}$, $I_{NaL}$, $I_{to,f}$, $I_{CaL}$, $I_{K1}$), 25-125% ($I_{Kr}$, $I_{Ks}$), 50-150% ($J_{rel}$, $J_{up}$), and 75-175% ($I_{NaCa}$, $I_{NaK}$). The ranges were chosen a priori to enable sufficient phenotypic diversity including vulnerability to EADs. 343 out of the 1000 models passed all the calibration criteria based on comparing biomarkers of the simulated models to experimentally observed ranges in human [11,119], summarized in **Table S6** listed previously.

The population was used to test 61 reference compounds with known arrhythmic risk at multiple concentrations versus the therapeutic dose: 1x, 3x, 10x, 30x, and 100x, using drug $IC_{50}$ values and Hill coefficients as in [118]. The occurrence of drug-induced repolarisation abnormalities, such as EADs, in the population of models was assessed and was used to calculate the Torsades de pointes (TdP) score as in [118]. CredibleMeds (Woosley and Romero, 2015) was used as gold standard for clinically established TdP risk, dividing the reference compounds in four categories: 1, high risk; 2, possible risk; 3, conditional risk; 4, safe (when not included in CredibleMeds). We used the most recent version of the CredibleMeds data, which differs from data used previously as the clinical reference [72,118], slightly reducing the performance of our model compared to other studies, as detailed in Results.

Compared to comparable prior *in silico* trials on this battery of compounds [72,118], which used 62 compounds, we excluded $BaCl_2$. This toxic salt is to our knowledge not used medically, and it is unlikely it would be listed in Crediblemeds in the first place. We did nevertheless confirm the drug is arrhythmogenic at higher concentrations when the population is exposed to it (not shown), which is in line with a range of experimental studies indicating its arrhythmogenic profile [120–122].

Based on experience when running studies drug safety prediction in prior models [72,118], we additionally curated the data describing channel blockade applied to the baseline model when we were aware of omission of an important drug effect in the data. This affected the formulation of 4 drugs: lidocaine, mexiletine, amiodarone, and cilostazol. For lidocaine and mexiletine, two near-identical formulations were present in the source data, differing only in their effect on late sodium current $I_{NaL}$ (In silico trials simulate multiple drug formulations when multiple source datasets describing pharmacological effects are available, using the most hazardous prediction to generate the final arrhythmic score). Given that it is well established that these two drugs block $I_{NaL}$ [123] (in fact, preferentially over $I_{Na}$), we decided to exclude the formulations without $I_{NaL}$ inhibition. Following this, we confirmed that no other drugs in the dataset should be excluded because of two near-identical formulation, differing only in inclusion of $I_{NaL}$ effect. In the case of amiodarone, potent chronic effects on key repolarizing potassium currents was reported [124,125]. In line with the study by Kamiya et al. [124], which used 100 mg/kg/day (ca. 10fold of therapeutic dose for arrhythmia treatment [126]), we applied the observed effect for simulations of 10x concentration and greater: a 29% and 83% reduction in $I_{Kr}$ and $I_{Ks}$, respectively. For cilostazol, the described $IC_{50}$ values describe direct channel block, but entirely ignore the primary effect of the drug, i.e., inhibition of PDE3, leading to increased $I_{CaL}$, the main ionic current promoting EADs and TdP. Consequently, we increased $I_{CaL}$ by +10% at 10x concentration, +22% at 30x, and +40% at 100x, based on Figure 2 in [127] (100x of therapeutic dose is 12.8 μM).

We note that the performance of ToR-ORd in our study differs subtly from the source 2019 publication where in silico drug assessment was also carried out [34]. This comprises two changes: First, the annotations of diltiazem and piperacillin changed from *no risk* to *conditional* risk in the reference data, leading to two additional false negatives in the current article, as both T-World and ToR-ORd predict the drugs to be safe. Second, a problematic formulation of lidocaine-induced changes was excluded in initial data curation carried out in this study, leaving a single other formulation of lidocaine in the dataset. Given that this formulation was driving a false positive result in our previous article from 2019 [34], we excluded this false positive from the contingency table for maximally fair comparison with T-World.

When simulating the effect of mexiletine in LQTS2, we explored two formulations of mexiletine: $MEX_{Crumb}$ (the same formulation as used in the in silico trial above, based on the data by Crumb et al. [128]) and $MEX_{Johannesen}$, based on a different set of measurements [123]. In the data by Johannesen et al., $I_{Na}$ and $I_{to,f}$ were not measured, and in the absence of measurements, we inferred the relative channel blocks based on the Crumb et al. IC50 values and Hill coefficients.  Drug concentrations were chosen to produce an approximately data-like change in action potential duration [129]: 25 µM for $MEX_{Crumb}$ and 10 µM for $MEX_{Johannesen}$. The effects on ionic currents at the given concentrations are summarized in (**Table S7**). LQTS2 was modelled as a 70% reduction in $I_{Kr}$ combined with an 82% increase in $I_{NaL}$ (the $I_{NaL}$ increase was described by Crotti et al. [129]).

***Table S7. Multipliers of ionic currents by two formulations of mexiletine.*** *Please note that the lack of effect of mexiletine on $I_{Ks}$ in $MEX_{Johannesen}$ follows from a direct measurement showing no visible effect, it is not due to a mere lack of data.*

| Ionic current | $MEX_{Crumb}$ (25 µM) | $MEX_{Johannesen}$ (10 µM) |
|---|---|---|
| $I_{Na}$ | 0.656086250445016 | 0.818648051180129 |
| $I_{NaL}$ | 0.192007760172465 | 0.5335 |
| $I_{to,f}$ | 0.920173174700735 | 0.96368476372007 |
| $I_{CaL}$ | 0.827890160668684 | 0.805 |
| $I_{Kr}$ | 0.756451488675938 | 0.9124 |
| $I_{Ks}$ | 0.560748793105810 | 1 |

#### Type 2 diabetes modelling

Data on cellular remodelling in Type 2 diabetes (T2D) in human remain rare, with animal data being heterogeneous and dependent on a specific animal model chosen [130]. We based our baseline model of T2D on the most comprehensive human-based dataset by Ashrafi et al. [131]. Compared to the model constructed there, we reduced the extent of $I_{Kr}$ reduction and $I_{NaCa}$ increase, as inclusion of the original values leads to unrealistically exaggerated phenotype (extremely long action potential and very small calcium transient). In addition, we omitted the change in $I_{K1}$, given relatively consistent data in animals suggesting no major change [130]. Given studies indicating reduced SERCA pump activity in diabetes, we have reduced SERCA pumps by 20% [132,133]. We label this baseline diabetic model D1 (**Table S7**).

To explore additional effects reported in literature, we formed model D2 by taking D1 as a starting point and doubling $I_{NaL}$ and increasing activity of CaMKII by increasing the $\alpha_{CaMKII}$ parameter by 50% (**Table S7**), reflecting $I_{NaL}$ and CaMKII increase described in the literature [134–136].

Finally, a part of animal-based studies on T2D indicates reduced, rather than slightly increased $I_{CaL}$. Given the importance of $I_{CaL}$ for the risk of EAD formation and thus arrhythmogenesis, we explored this possibility by constructing D3, D5 (based on D1) and D4, D6 (based on D2) by setting $I_{CaL}$ availability to 90% and 80% of $I_{CaL}$ compared to undiseased T-World (**Table S8**). The reductions replace the $I_{CaL}$ increase of D1 and D2; they are not used to merely multiply it.

***Table S8: Overview of T2D model versions explored in the article.*** *Percent given represent relative change to baseline (i.e., it is not additional).*

| Model | Description |
|---|---|
| **D1** | 121% $I_{Na}$ and $I_{NaL}$, 114% $I_{CaL}$, 126% $I_{to,f}$, 70% $I_{Kr}$, 95% $I_{Ks}$, 150% $I_{NaCa}$, 90% $J_{rel}$, 80% Jup (SERCA) |
| **D2** | D1, 200% $I_{NaL}$, 150% $\alpha_{CaMKII}$ |
| **D3** | D1, 90% $I_{CaL}$ |
| **D4** | D2, 90% $I_{CaL}$ |
| **D5** | D1, 80% $I_{CaL}$ |
| **D6** | D2, 80% $I_{CaL}$ |

#### NaV1.8 current investigation

A separate version of T-World including the NaV1.8 current was created by adding the Choi-Waxman formulation [137] to a baseline model. The equations for this two-gate Hodgkin-Huxley model are reproduced below. The only change to the original version is a small update to $h_\infty$, giving it a small degree of activity during action potential plateau, in line with its nature as a contributor to late sodium current.

$$\alpha_{m,NaV1.8} = 2.85 - \frac{2.839}{1 + e^{\frac{V-1.159}{13.95}}}$$

$$\beta_{m,NaV1.8} = \frac{7.602}{1 + e^{\frac{V+46.463}{8.8289}}}$$

$$m_{\infty,NaV1.8} = \frac{\alpha_{m,NaV1.8}}{\alpha_{m,NaV1.8} + \beta_{m,NaV1.8}}$$

$$\tau_{m,NaV1.8} = \frac{1}{\alpha_{m,NaV1.8} + \beta_{m,NaV1.8}}$$

$$h_{\infty,NaV1.8} = 0.02 + \frac{0.98}{1 + e^{\frac{V+32.2}{4}}}$$

$$\tau_{h,NaV1.8} = 1.218 + 42.043 \cdot e^{\frac{-(V+38.1)^2}{2 \cdot 15.19^2}}$$

$$\frac{dm_{NaV1.8}}{dt} = \frac{m_{\infty,NaV1.8} - m_{NaV1.8}}{\tau_{m,NaV1.8}}$$

$$\frac{dh_{NaV1.8}}{dt} = \frac{mh_{\infty,NaV1.8} - h_{NaV1.8}}{\tau_{h,NaV1.8}}$$

The ionic current through NaV1.8 in a compartment X (dyadic or subsarcolemmal) is

$$I_{NaV1.8,X} = f_X \cdot g_{NaV1.8} \cdot m_{NaV1.8} \cdot m_{NaV1.8} \cdot \left(V - E_{Na,X}\right)$$

The variable $f_X$ is the fraction of the current in compartment X. Like with most other currents, 11% of the current was placed in the dyadic compartment, with the rest placed in the subsarcolemmal one. $E_{Na,X}$ is the equilibrium potential for sodium in the given compartment.

For the purpose of simulations visualized in Figure 5 of the main manuscript, the three explored values of the conductance of $I_{NaV1.8}$ ($g_{NaV1.8}$) were 0, 0.1085, and 0.3 mS/μF (all substantially lower than the conductance of the main NaV1.5 current $I_{Na}$ with $g_{Na}$ > 22 mS/μF).

#### Graphical user interface

We provide a graphical user interface (GUI) for simulating T-World for a choice of parameter values and a variety of stimulation protocols. Users can visualize different variables and download simulation data for further analysis. The GUI supports four stimulation protocols: (i) regular pacing, (ii) S1-S2 pacing, (iii) pacing over a range of intervals (suitable, e.g., for alternans), and (iv) regular pacing followed by a pause (suitable, e.g., for DADs). For protocol (ii), the GUI generates a restitution curve, while for protocol (iii), it displays a bifurcation diagram of the action potential duration (APD) or another variable as a function of the pacing interval. The GUI is implemented in Plotly Dash and uses Myokit [138] for model simulation. It is accessible online or available for download. The simulator and further details are available at **<will-be-provided-upon-acceptance>.**

#### Tips on the use of genetic algorithms in model development

Throughout the development of T-World, we frequently used multiobjective genetic algorithm (MGA, @gamultiobj in Matlab) to fit parameters of single model components, as well as for integrating components together. In general, a range of protocols would be simulated with a candidate model, with the simulation outputs compared to reference values based on experimental data. Below we share several insights we have obtained throughout the process, which could be valuable to other users of the method.

- **Number of fitness dimensions.** Each simulated protocol produces a single error describing the deviation of simulation from experiments. We would often work with 10-20 different errors that are to be optimized. Putting them in a single fitness criterion as a weighted sum makes the MGA more likely to get stuck in a local (non)optimum. However, having each error as a separate dimension of a fitness function yields a Pareto front where the vast majority of creatures are useless (because being useless in a single dimension tends to be sufficient for the resulting model to be useless). As a result, the evolution tends to explore mostly useless creatures, and does not progress well. In practice we found that 3 fitness dimensions (in which the errors are allocated as weighted sums) was a sweet spot for most of the genetic algorithms, with 2 and 4 also useful. In general, we aggregated logically connected errors in the same dimension, i.e., fitness(1) would consider errors of basic CaT biomarkers, fitness(2) would consider alternans errors, etc.
- **Population size and number of generations.** We used a two-stage procedure, where a large[1] population was first simulated for several (5-10) generations. The resulting Pareto front was used as a starting population for a second stage MGA, which was ran with a smaller population. Using a large population has the advantage that it samples the parameter space well initially, and it explores a more diverse population than if a small population was used at the start. However, a large population subsequently undergoes a slow and inefficient evolution. On the other hand, using a smaller population accelerates the evolution (with a single generation taking much less time to simulate) – but if used exclusively, the initial parameter space would not be sampled as well. The two-stage procedure combines good sampling of initial parameter space with reasonably fast evolution later.
- **Early killing of ultimately infeasible creatures.** Early in the evolution, it is important to consider even creatures with fitness that will be ultimately deemed useless (e.g. models excellent in fitness(2) and fitness(3), but very poor in fitness(1)). While the creatures are not yielding useful models themselves, they can carry valuable information that can be combined with other creatures in crossover, creating excellent creatures and thus models in the future. However, in the second stage of optimization, which is more about accelerating evolution and parameter fine-tuning, it may be practical to avoid having too many creatures that will be ultimately deemed useless. To this end, we would sometimes include guards that set fitness to Inf (effectively guaranteeing the removal of such creatures from the evolved population) when any of the fitness dimensions is bad enough. Similarly, when a single error that is optimized is known to often drive such infeasibility, it is worth to calculate this error at the start of the fitness function and check right after what the error value is – if it is poor, the entire fitness can be

[1] What “large” means depends on the problem and fitness function. For our more comprehensive simulations where a single fitness evaluation took around 3 minutes, a population of 3600 creatures was considered large. However, for faster-to-run fitness functions, hundreds of thousands creatures can be simulated.

stopped there (returning vector of Inf to make sure the creature is discarded), before the rest of the fitness function is simulated, which saves time.

- **Saving intermediate results**. It is very useful to save every generation throughout the optimization, which can be achieved by using the 'OutputFcn' option of @gamultiobj. This enables intermediate monitoring of the evolution, early termination of optimization without losing any results, or restarting the evolution from the last saved point e.g. in case of power outages.
- **Pareto front size limit.** We observed that the Matlab implementation of MGA seems to have an internal limit of the Pareto front size. As a result, when the Pareto front size reaches 350, some creatures start being removed throughout the generation, so that the size of Pareto front is maintained. For some reasons, we noted that the removed creatures would be often the ones that encode the practically best models. For this reason, we suggest that the number of creatures on the Pareto front is monitored throughout the simulation, and caution is exercised when the limit of 350 is hit – it is possible that good creatures are lost from that point on. This issue is exacerbated with a growing number of dimensions of the fitness function. It can be partly ameliorated by the early discarding of creatures described above, which generally limits the space of feasibility of fitness values.

#### Notes on implementation

The default implementation is in Matlab, which was subsequently converted to CellML, C, and CUDA, enabling loading T-World in simulators such as Chaste[139], MonoAlg3D[106], or Myokit [138], with the latter also offering conversion to other languages.

The Matlab version is structured with a logic similar to ToR-ORd [72]:

1. The user defines parameters of the simulation (e.g. number of beats, stimulation rate, concentrations, multipliers of current conductances, etc.) in a Matlab simulation script, which then calls a runner function. When the simulation is finished, a structure with ionic currents and fluxes over time is extracted, and may be easily used to visualize traces (e.g. 'plot(currents.time, currents.V)' to plot membrane potential, etc.).
2. The runner function receives the parameters, unpacks them into single variables, and sets the undefined parameters to default values (in this way, the user has to define only the parameters that do not take the default value). Subsequently it solves the ordinary differential equation system in a model function using the ode15s solver.
3. The model function, containing the definition of the cell model, calculates the derivatives of state variables based on the input state variables vector.

This design has several advantages. We believe that it maximizes user comfort by allowing them to define a simple structure of parameters as an input, without them having to touch the model code itself. Using structures is convenient to neatly pack Matlab variables. Unfortunately, their use in Matlab also comes with a rather severe performance penalty. For this reason, we use them only on the user side for convenience; however, the runner function unpacks them into single, unstructured variables, which are then passed to the ODE solver itself. Unpacking the structure once does not cause any performance problem – however, using the structure of parameters in the model function would, which is why we avoid this.

An advantage of the model function being kept separate is that different model functions can be easily used in the Matlab framework we provide. E.g. a model function specific to a disease may be created by the operator, and then used by including in the structure of parameters which model function should be used (e.g. 'parameters.model = @model_HFrEF;').

An additional aspect of modularity is that the model function of T-World is structured logically into functions calculating different ionic currents and other components (in contrast to some other models using monolithic code). It is therefore easy to navigate the code, and to replace a chosen ionic current by including a different formulation as a function. We also use reasonably named variables throughout the code; e.g. 'ions_ca_i' referring to intracellular calcium concentration. This is in contrast with many other models which refer to elements of the state vector (e.g. y(38) or X(8), which  is difficult to navigate for researchers not familiar with the given model family.

Finally, we avoided Matlab global variables in T-World, given their performance penalty and given that they complicate parallelization of simulations. The one exception is the version of T-World with stochastic DADs, where we were unable to avoid their use when relying on ode15s for ODE solving.

#### Other models

T-World was compared in this study to ToR-ORd [72], Morotti 2021 [3], and TP06 models [77]. ToR-ORd was downloaded from Github (https://github.com/jtmff/torord), as was the Morotti 2021 model (https://github.com/drgrandilab/Morotti-et-al-2021-Cross-species-translators-of-electrophysiological-response). The endocardial TP06 model was simulated using a CellML version (https://models.cellml.org/workspace/tentusscher_panfilov_2006) loaded and simulated in Myokit [138].

#### Experimental methods

Experimental data for human S1S2 restitution, used to investigate sex differences in restitution slope, were collected as a part of prior study[41], with methodology described there.

### Supplementary notes

#### Supplementary note 1: Comparison AP morphology between state-of-the-art models

We compared T-World to several other computational human ventricular cardiomyocyte models that represent the state of the art in various areas: ToR-ORd[72] from the Rudy family of models, Morotti2021[3] from the Bers/Grandi family of models, and the TP06 model[77]. We compared the models' AP morphology, using the endocardial cell variants as the baseline (**Figure S7A)**. T-World is relatively similar to ToR-ORd, with the minor improvement that there is no notch after AP peak in T -World (see **Figure S7B** for an inset view), consistent with most human data on endocardial AP morphology that reflect minimal $I_{to,f}$ in the human endocardium[5]. The Morotti2021 model also does not manifest a notch after the AP peak, but it shows a kink in the early plateau, which is generally not observed in living cells, and which results from a calcium-handling issue inherent to this model family that is discussed more in **Supplementary Note 3**. The main limitation of the TP06 AP is its lack of AP triangulation (APD90-APD40), with a long, near-flat plateau followed by a steep repolarization.

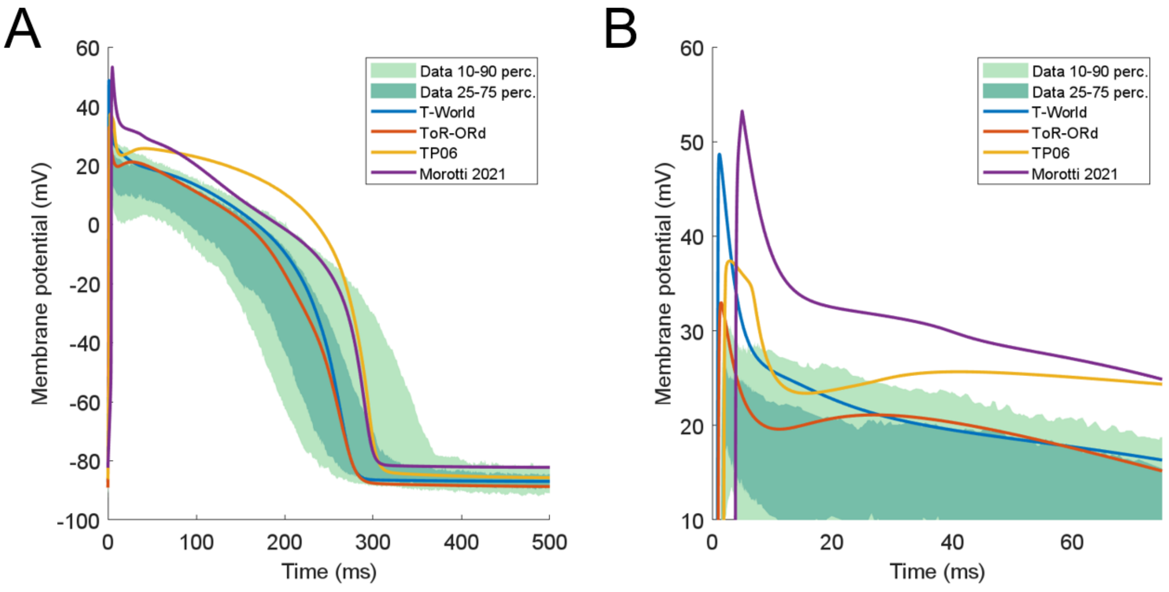

***Figure S7. Comparison of AP morphology between models. A)*** *Comparison of action potentials (APs) between four major computational human ventricular cardiomyocyte models and experimental data [5].* ***B)*** *A zoomed-in version panel A, showing a notch following the AP peak in ToR-ORd and TP06 models, that is typically not seen in endocardial cells. In addition, TP06 model shows unusually biphasic transition from peak to the notch.*

### Supplementary note 2: Comparison of models' responses to channel blocking drugs

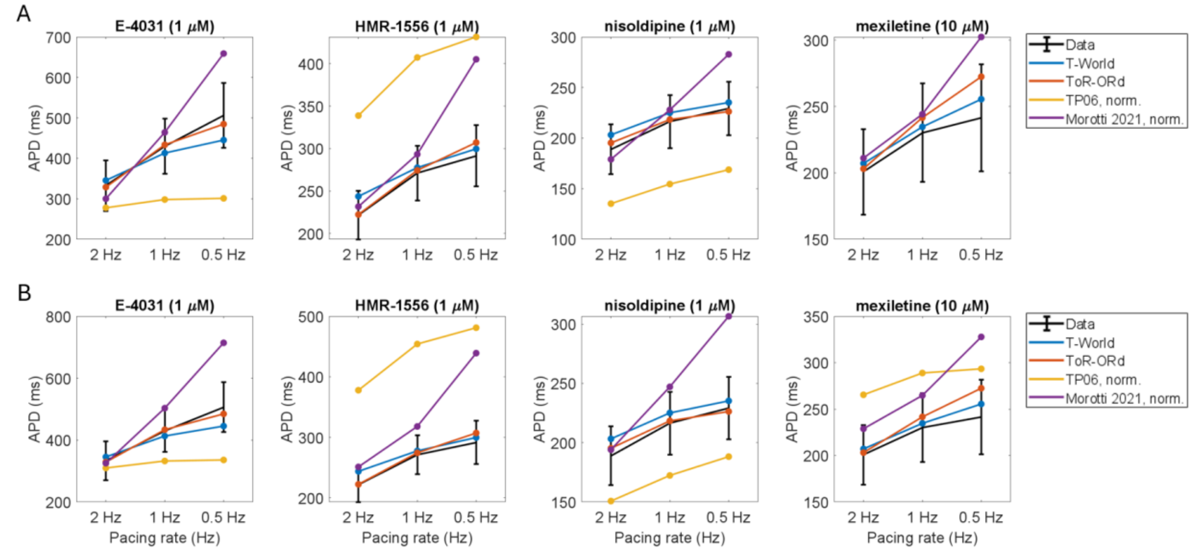

***Figure S8. Comparison of the response of four models to simulated channel blocking drugs.*** *Independent validation of the APD prolongation or shortening induced by 1 μM E-4031 (70% $I_{Kr}$ block), 1 μM HMR-1556 (90% $I_{Ks}$ block), 1 μM nisoldipine (90% $I_{CaL}$ block), and 10 μM mexiletine (54% $I_{NaL}$, 9% $I_{Kr}$, 20% $I_{CaL}$ block) at 0.5, 1.0 and 2.0 Hz pacing in the four models. Drug concentrations and their effects on channel blocks are based on O'Hara et al. [5]. In panel **A**, outputs of the TP06 and Morotti2021 models were scaled by $APD90_{model}/270$ to compensate for their slightly longer baseline APD: without the normalization, the discrepancy between those models and data is exaggerated artificially. The outputs of TP06 are not shown for mexiletine, as the model lacks the representation of the primary target $I_{NaL}$. For completeness, the version of the figure without normalization and with TP06+mexiletine is given in panel **B**.*

Comparing T-World to other state-of-the-art models (**Figure S8**), T-World retains the strong performance of ToR-ORd, showing excellent agreement with experimental data despite having a very different cell architecture and calcium handling. The Morotti2021 model is also mostly in agreement with the data at the more physiological heart rates of 1 Hz and 2 Hz. The TP06 model shows substantial discrepancies with the experimental data, both in the balance of key repolarizing currents (response to E-4031 vs HMR-1556), as well as in response to inhibition of depolarizing currents. In particular, the absence of the $I_{NaL}$ current in the model leads to excessive APD shortening during $I_{CaL}$ block with nisoldipine, whereas such overt shortening is prevented in the other models by the presence of $I_{NaL}$ as partial carrier of depolarizing current.

### Supplementary note 3: Excitation-contraction coupling

Calcium-induced-calcium-release (CICR) in cardiomyocytes involves a local activation of cardiac ryanodine receptors (RyR) on the sarcoplasmic reticulum (SR) by calcium entering the cells through L-type calcium channels ($I_{CaL}$), and this induces additional calcium-dependent RyR activation throughout the cell. Different models take a different approach to representing this process, with distinct advantages and disadvantages. Models like ORd and ToR-ORd coupled the RyR release flux directly to $I_{CaL}$, resulting in a realistic CaT shape, but this is mechanistically unrealistic and prevents naturally occurring DADs resulting from spontaneous SR calcium releases due to RyR dysfunction or calcium overload. On the other hand, the Bers/Grandi-like models, including Morotti2021, make RyR depend on local calcium concentrations, which is mechanistically accurate, but it induces problems with delayed SR calcium release, as shown below. To overcome these limitations, we took a hybrid approach in T-World, with a very small population of $I_{CaL}$-dependent RyR (representing the most closely associated $I_{CaL}$ and RyR clusters) that provides an early influx of calcium which helps to prime the majority population of Ca-sensitive RyR. The latter follows a revised formulation used in Morotti2021 and prior models of the family. This solution enables a model of appropriately timed CICR that is simultaneously calcium sensitive and thus capable of generating DADs.

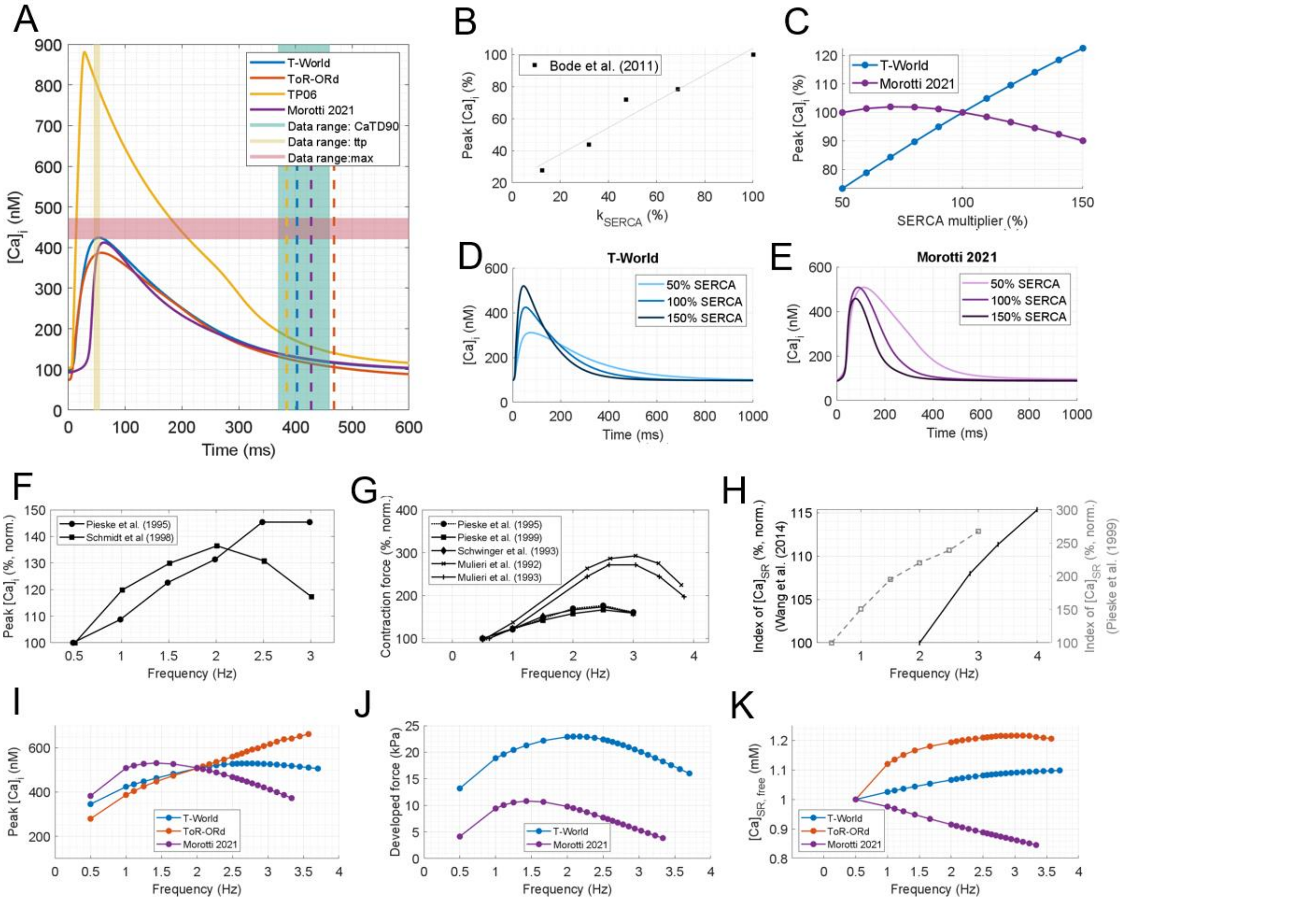

***Figure S9. Properties of excitation-contraction coupling. A)*** *Comparison of the four models and experimental ranges for calcium-transient (CaT) properties as in Figure 1; dashed lines give the CaT duration at 90% recovery for the four models.* ***B)*** *Experimental data on the relationship between SERCA activity and peak calcium concentration* [14]*.* ***C)*** *Corresponding simulations of the same relationship in T-World and Morotti2021.* ***D,E)*** *Sample traces of CaTs at three different SERCA levels.* ***F-H)*** *Experimental data for rate-dependence of peak calcium concentration* [18,19]*, developed force* [18,20–23]*, and sarcoplasmic reticulum (SR) calcium concentration* [20,24]*. In J), the Pieske et al. study in human used rapid cooling contractures, which is a less direct estimate of* $[Ca]_{SR}$ *than the direct measurement with a fluorescent dye by Wang et al. in rabbit.* ***I-K)*** *Corresponding simulations of rate-dependence in computational models. In J, only the models that include force generation are shown. The TP06 rate-dependence is very steep and is thus shown separately in* ***Figure S13****. See* ***Figure S10*** *for a comment on the importance of pre-pacing duration in ToR-ORd.*

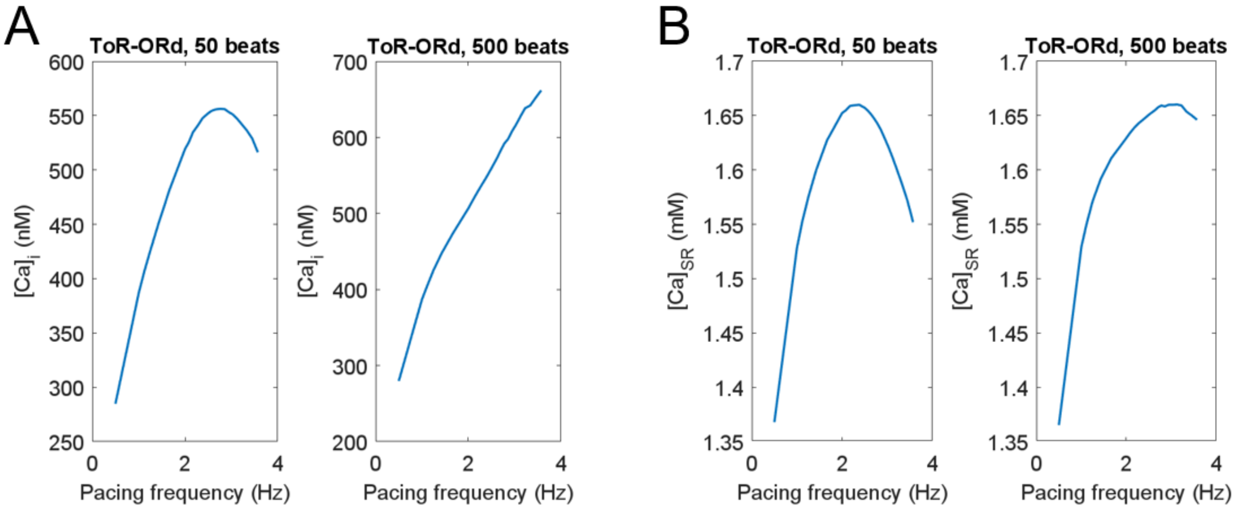

***Supplementary figure S10. Effect of pre-pacing duration on biomarkers in ToR-ORd.*** *We note that the monotonically positive rate-dependence of peak [Ca]ᵢ ToR-ORd shown in main text Figure 2 is not at odds with the more data-like biphasic rate-dependence shown in Figure 12 of the article describing the predecessor model of O'Hara-Rudy [5]. In our work, we show a steady-state value after 500 beats, whereas the figure in the O'Hara-Rudy article was generated following much shorter pacing. To clearly show the importance of pre-pacing duration on the pattern of CaT peak across rates, we compared this in ToR-ORd simulated for 50 beats (**panel A**, left) and 500 beats (**panel A**, right). The model paced for 50 beats appears more data-like in its biphasicity, but this is not a behavior that would hold close to steady-state. The difference is to a large extent driven by differences in the calcium content of the sarcoplasmic reticulum (**panel B**).*

The T-World CaT has relatively similar kinetics to that of ToR-ORd (**Figure S9A**). An attractive property of both models is their relatively linear CaT upstroke, which takes off early from the resting value following electrical cell activation. This leads to a relatively small delay between AP upstroke and CaT upstroke, in agreement with simultaneous measurements of calcium and membrane potential in cardiomyocytes [7,140]. However, T-World is much more mechanistically realistic than ToR-ORd, as it has a predominantly calcium-mediated CICR. The Morotti2021 model, which also has calcium-mediated CICR, shows a relatively long initial phase with slowly rising calcium level (**Figure S9A**), which is not supported by myocyte imaging data. The TP06 model manifests a very large and early-peaking CaT (**Figure S9A**), which is not consistent with human data biomarkers. In addition, its calcium concentrations in the dyadic compartment where calcium is released exceed experimental estimates by an order of magnitude, leading to problems with $I_{CaL}$ morphology (see **Supplementary Figure S11**).

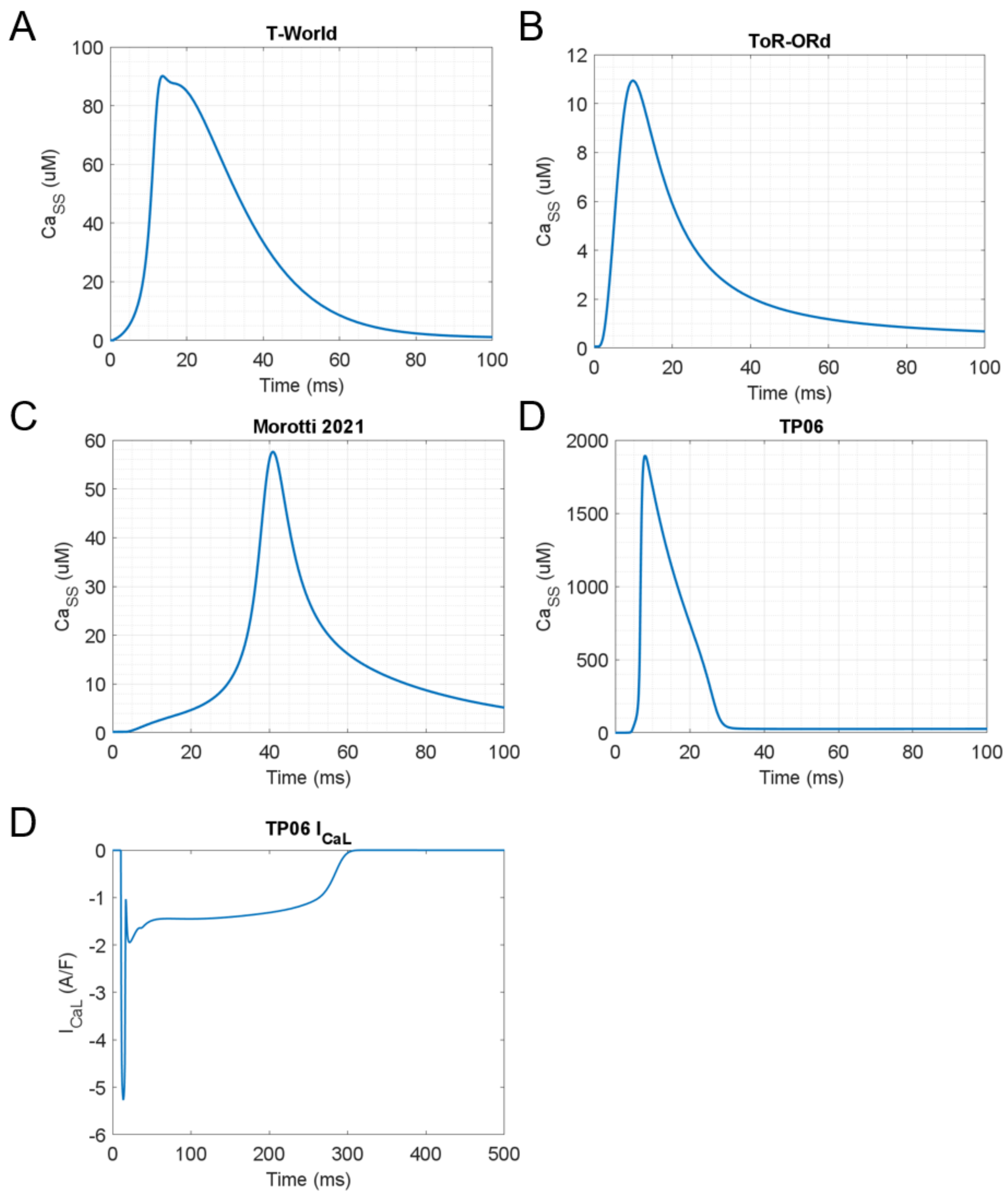

***Figure S11. Dyadic concentrations of calcium in four models. A-D)*** *dyadic calcium concentrations in T-World, ToR-ORd, Morotti 2021, and TP06.* ***E)*** *L-type calcium current during endocardial AP in TP06, showing spike-and-dome morphology which results from the extremely high dyadic concentration even at baseline conditions. This is to our knowledge not supported by data, and is furthermore exacerbated if a more standard formulation of driving force of $I_{CaL}$ was used. See the section on $I_{CaL}$ in Supplementary Methods for further comments on the driving force equation used in TP06.*

The biphasic upstroke of the Morotti2021 CaT (and other models of the same family) is a consequence of the relatively late-peaking SR calcium release flux through RyR (**Supplementary Figure S12**). Experimental studies mostly place the time to peak calcium release at around 10 ms [7–10] and T-World, ToR-ORd, and TP06 are consistent with this, whereas the Morotti2021 model peaks around 40 ms. First, this directly, leads to a slow early rise of the CaT. Furthermore, it delivers calcium for calcium-dependent $I_{CaL}$ inactivation later than it should following the $I_{CaL}$-based calcium influx. As such, the calcium-dependent inactivation needs to be parametrized to respond to relatively low calcium levels present at ~10-15 ms. Finally, the late elevation of dyadic calcium, peaking 30-40 ms after electrical stimulus is sensed by the sodium-calcium exchanger (NCX) present there. This leads to local calcium efflux, and given the electrogenic nature of NCX, a depolarizing current is generated around 30-40 ms that yields the suboptimal AP morphology shortly after peak, characteristic for this family of models. For those reasons, creating a CICR formulation with a faster time to peak release, while maintaining the attractive calcium-sensitive nature of Bers/Grandi models, was one of the important development targets when creating T-World.

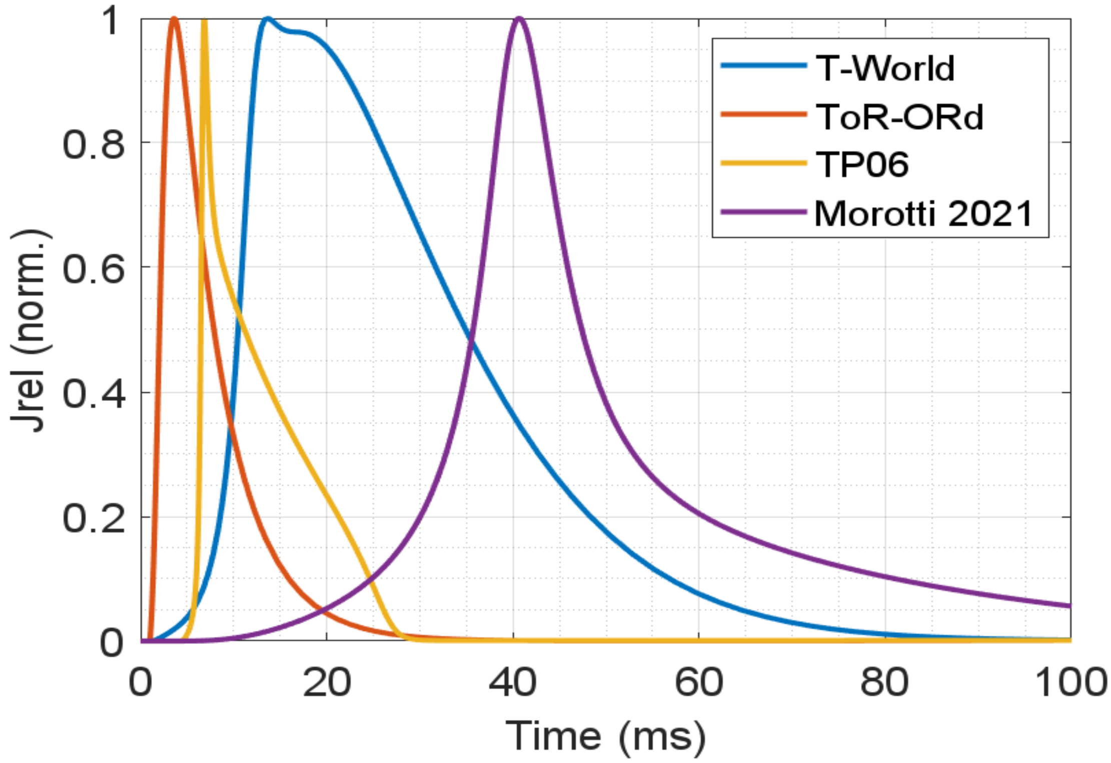

***Figure S12.*** *Comparison of timing of normalized SR release through RyR in four models.*

Another property of the calcium-handling system in the Bers/Grandi models that we wanted to improve is the response to changes in SERCA pump activity. SERCA activity is experimentally closely associated with CaT amplitude (**Figure S9B**), with SERCA potentiation increasing CaT amplitude and contractility [16,17], and vice versa for reduced SERCA activity [14,15]. Capturing this property is key for modelling conditions where SERCA function is altered, including various diseased states or during sympathetic stimulation. The baseline Morotti2021 model (and preceding models with the same CICR model) do not accurately reproduce the SERCA-dependence of CaT amplitude (**Figure S9C,E**), whereas the revised calcium-handling system in T-World provides a good agreement with the data (**Figure S9C,D**).

A crucial ability of cardiomyocytes is to adapt calcium handling and contraction to different pacing rates, which enables the heart to match the physiological needs of the human body. In humans, the peak CaT first increases with increasing stimulation rate, but subsequently decreases somewhat as the pacing rate is further increased (**Figure S9F**) [18,19]. A range of human studies show a similar biphasic pattern for the rate dependence of contractile force (**Figure S9G**) [18,20–23]. Conversely, data from human [20] and rabbit [24] cardiomyocytes indicate that the SR calcium concentration increases continuously with pacing rate up to the maximum measured rate (**Figure S9H**).

The T-World model is in good agreement with those data on rate-dependence, showing a nonmonotonic calcium- and force-frequency relationship, as well as positive $[Ca]_{SR}$-frequency relationship (**Figure S9I-K**). This is nontrivial, particularly given that the SR calcium load-release relationship is positive and steep in experiments [86] and computer models. Therefore, seeing an increase in SR content together with reduced CaT/contraction at the fastest pacing rates requires sufficient refractoriness of $I_{CaL}$ and/or RyR to outweigh the load-release relationship.

The agreement between T-World and multiple relevant datasets holds even quantitatively. E.g. the peak $[Ca]_i$ in **Figure S9I** is 150% of the value at 0.5 Hz, similar to data (**Figure S9F**). The peak developed force in T-World in

**Figure S9J** is ca. 170% of the value at 0.5 Hz, which is very close to three experimental datasets in **Figure S9G**. By contrast, other models show several qualitative discrepancies with the rate-dependence data. The ToR-ORd model shows a fully positive calcium-frequency relationship at steady state (**Figure S9I**). The Morotti2021 model shows a mostly flat or negative calcium/force-frequency relationship and $[Ca]_{SR}$-frequency relationship between 1 Hz and 3 Hz (**Figure S9I-K**). It therefore rather resembles the phenotype of cardiomyocytes isolated from heart failure patients [20,22]. The TP06 model shows very steep rate-dependence of peak calcium, and while it is also appropriately biphasic, this seems to result from a biphasic $[Ca]_{SR}$-frequency relationship, at odds with the data (**Figure S13**, shown separately given the different ranges).

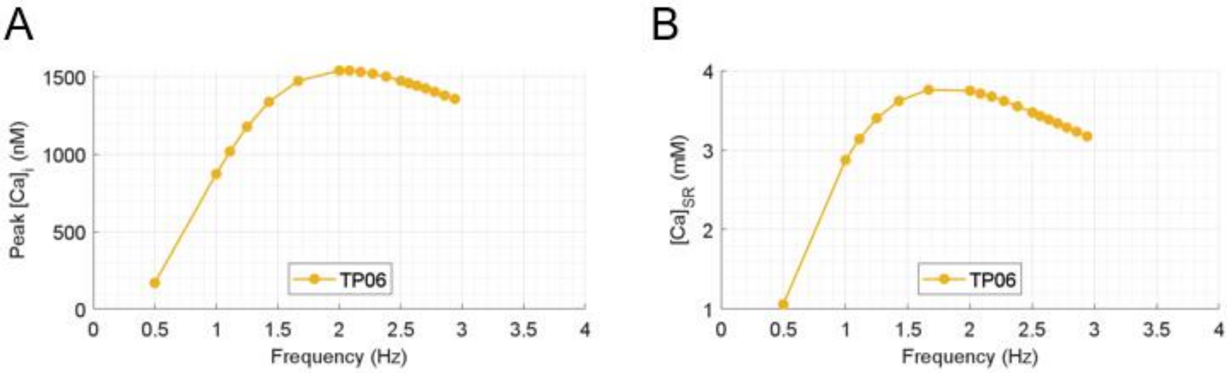

***Figure S13.*** *Rate dependence of calcium transient peak and sarcoplasmic reticulum calcium loading in the TP06 model.*

Characteristics of $I_{CaL}$ are critical for realistic calcium handling. **Figure S14** demonstrates that $I_{CaL}$ in T-World manifests: 1) a data-like current-voltage relationship, which follows from activation and inactivation properties, as well as maximum current amplitude, 2) a data-like recovery from refractoriness (substantially improving upon ToR-ORd), 3) a good balance between voltage-dependent and calcium-dependent inactivation.

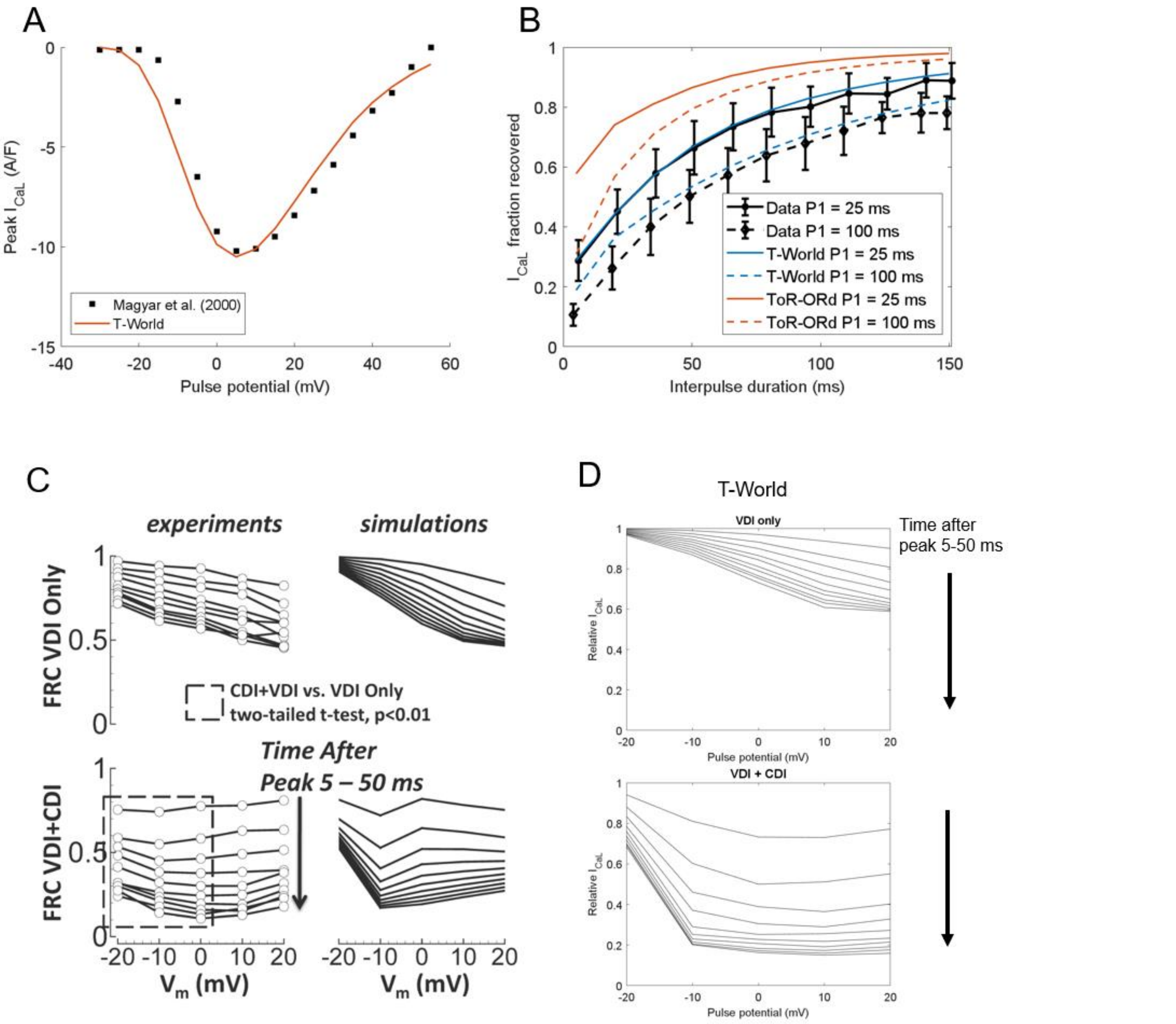

***Figure S14. Properties of L-type calcium current in T-World. A)*** *Current-voltage (I-V) relationship, showing a similar shape and amplitude between T-World and experimental data [12].* ***B)*** *Simulated P2-P1 protocol, a two-pulse protocol used to describe the recovery from refractoriness of $I_{CaL}$. The duration of the first pulse (which partly inactivates $I_{CaL}$) is either 25 ms or 100 ms (given in the legend). It is clear from the data that the longer the first pulse, the more extensive the inactivation of $I_{CaL}$ (dashed lines lie under corresponding solid lines). The T-World model provides a good agreement with the experimental data by Fulop et al. [13], unlike ToR-ORd, which underestimates the $I_{CaL}$ refractoriness.* ***C)*** *Separation of voltage-dependent inactivation (VDI) and Ca-dependent inactivation (CDI) of $I_{CaL}$, comparing experimental data with the O'Hara-Rudy article (reproduced based on [5]). The panels show fractional current at a given voltage pulse potential 5, 10, ..., 50 ms after the peak of evoked ICaL.* ***D)*** *Simulations of corresponding plots in the T-World model, which we performed as additional validation of the current formulation, given the relatively extensive changes from the preceding versions of the current in ToR-ORd (and O'Hara-Rudy). The simulations show a generally good agreement between the simulations and data, with pure VDI providing a relatively slow source of inactivation. We note that both experimental and simulation results will depend to some extent on specific conditions (such as pacing history and corresponding $[Ca]_{SR}$). In addition, it is known that Ba current used in the experimental study to obtain VDI is in fact a mix of VDI and a very small amount of CDI [141], thus slightly overestimating the VDI development speed. For those reasons, we believe it is not particularly useful to focus at whether a full quantitative agreement is achieved. Instead, it is encouraging that 1) in both O'Hara-Rudy and T-World, VDI+CDI develop much more rapidly than pure VDI, 2) VDI shows a clear voltage-dependence (faster inactivation VDI at higher potentials), whereas VDI+CDI are comparably flatter across most pulse potentials. We note that T-World was calibrated to achieve the behaviours in panels* ***A,B****, whereas panel* ***D*** *is independent validation.*

As additional validation, we confirmed that the model appropriately recapitulates the negative inotropic effects of sodium channel blockers (**Figure S15A**), and that the rate-dependence of sodium concentrations qualitatively matches experimental data **Figure S15B.**

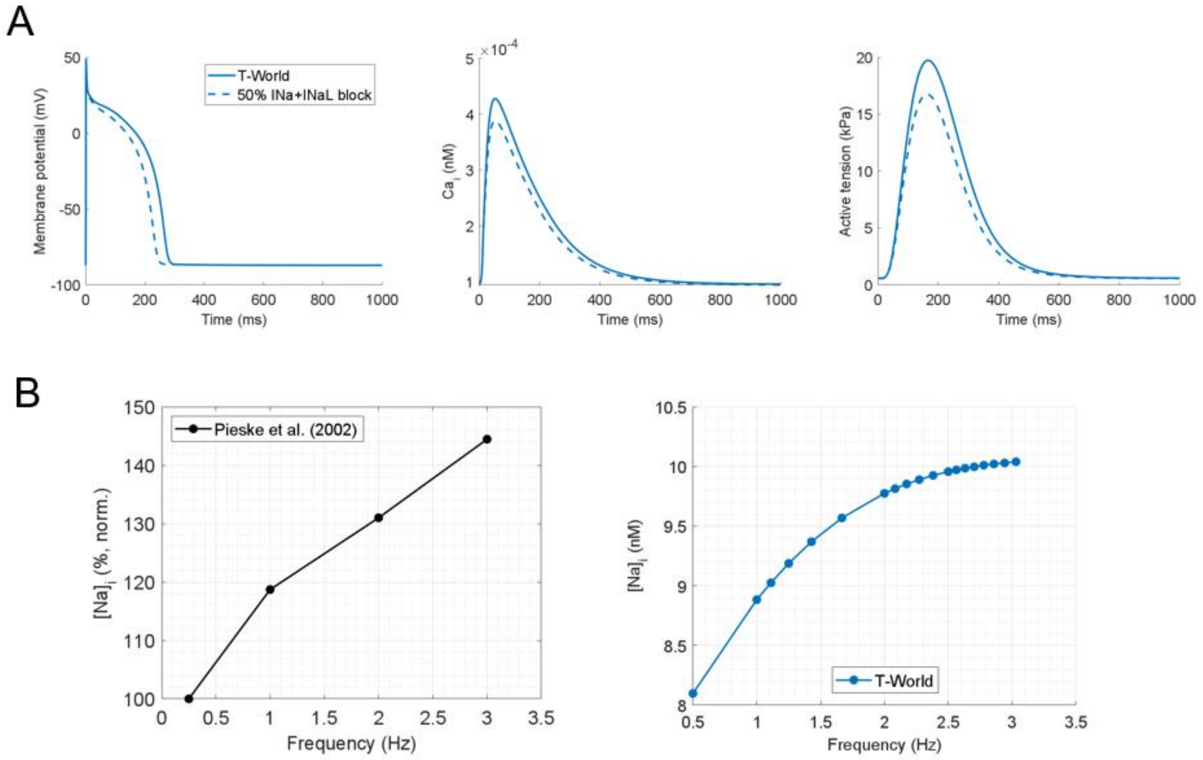

***Figure S15. Validation of T-World through sodium-related protocols***. ***A)*** *Response of the computational model to a combined 50% blockade of $I_{Na}$ and $I_{NaL}$, similar as done in Tomek et al.* [72]*. It shows APD shortening and a reduction in CaT amplitude and contractility, consistent with established negatively inotropic effect of sodium blocker drugs* [47–49]. ***B)*** *Experimental data* [50] *describing the rate-dependence of [Na]$_i$ concentration, and corresponding simulation of T-World, which also shows a positive rate-dependence relationship. In both model and the experimental study, [Na]$_i$ keeps increasing even at fast pacing where the contractility is already diminishing.*

Finally, we tested the relative contribution of SERCA, NCX, and pCa to calcium clearance during an action potential. T-World predicts 79.6%/20.15%/0.25% of calcium to be cleared by those means, which is consistent with studies in human-like species indicating a leading role of SERCA, 20-30% clearance by NCX, and a minimal contribution of pCa[51].

#### Supplementary note 4: β-adrenergic signalling

We calibrated T-World to achieve APD shortening with saturating activation of βARS, subsequently validating it by comparing the model outputs to human data on βARS-induced changes in CaT and contraction.

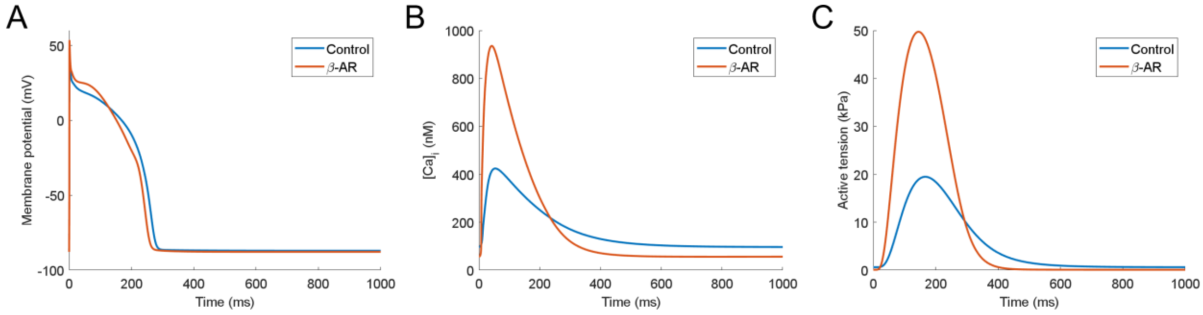

***Figure S16. Effect of sympathetic nervous stimulation on cell physiology. A-C)*** *show the impact of βARS on AP, CaT,, and contraction respectively.*

Activation of βARS in T-World causes a ~7% APD shortening (**Figure S16A**), in line with human studies indicating APD shortening following isoproterenol exposure [25,26]. The elevation of early plateau potentials in the model, resulting from the increased $I_{CaL}$ was also consistent with data from canine cardiomyocytes exposed to isoproterenol [142,143]. βARS activation induces a large 2.68-fold increase in CaT amplitude (**Figure S16B**). This is consistent with studies in humans and rabbits [60,61] that show a severalfold increase, although direct quantitative comparison is difficult, given uncertain mapping of calcium-induced fluorescence changes to calcium concentrations. The 24% shortening of CaT at half-amplitude is similar to the ~30% shortening observed in human samples [61]. Our model predicts a 2.63-fold increase in contractility (**Figure S16C**), which is generally in line with the highly heterogeneous human data that report an increase of 1.5-fold [62], 3.0-fold or 4.5-fold, depending on βARS-agonist [61], and 5.0-fold [63]. Changes in kinetics of upstroke and recovery of contraction are also highly heterogeneous across different studies, although there is general agreement that βARS accelerates development and recovery of contraction [61–63], as predicted by our model.

Similar predictions of substantially increased CaT amplitude and contraction are made by the Morotti2021 model, which also includes βARS, although APD does not shorten visibly in this model and there is almost no plateau elevation (**Figure S17**).

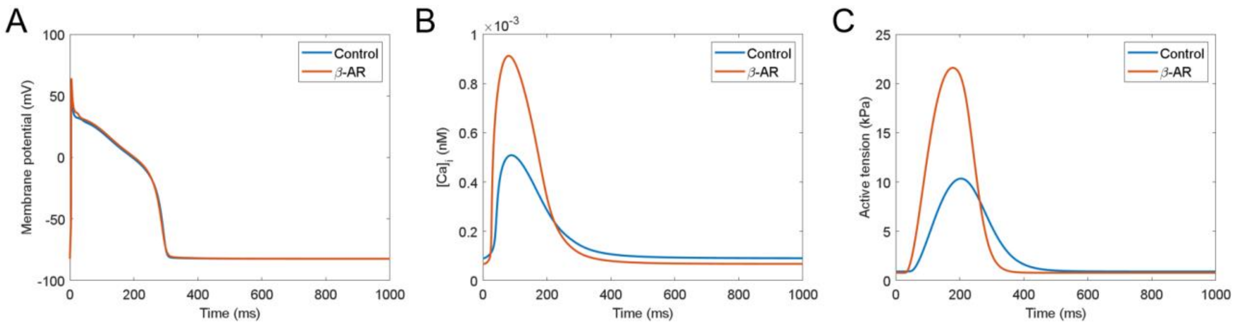

***Figure S17. Effects of βARS stimulation on the Morotti2021 model.*** *Shown are the effects of βARS on* ***A)*** *Action potential,* ***B)*** *$Ca^{2+}$ transient,* ***C)*** *Active tension.*

#### Supplementary note 5: Sex-differences

The female version of T-World manifests a prolonged APD (**Figure S18**), which is consistent with clinical and experimental data [52,53]. In addition, the female version has a slightly smaller CaT (**Figure S18B**) and peak developed force (**Figure S18C**), in agreement with available data [54,55]. Thus, sex-specific formulations of T-World correctly translate cellular sex differences in ionic currents and buffering into key differences in overall phenotype, enabling studies into sex differences in arrhythmic risk (discussed below).

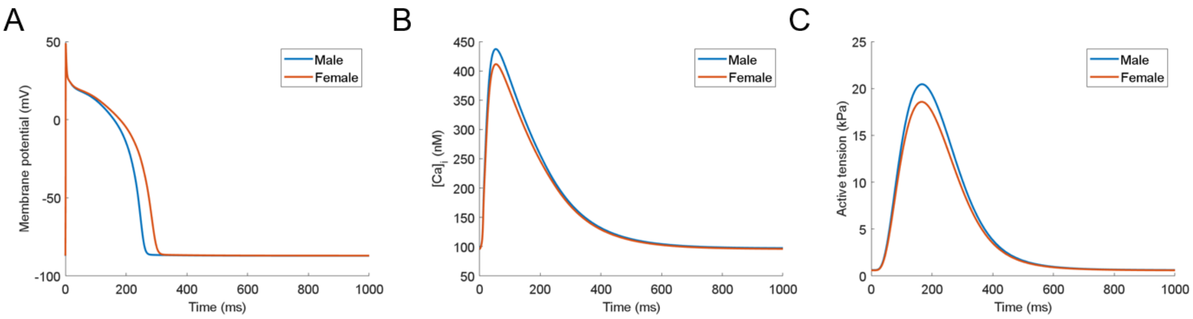

***Figure S18.*** *Sex differences between AP (**A**), CaT (**B**) and contraction (**C**).*

#### Supplementary note 6: Stability of arrhythmic behaviours

A population of 1000 models with key ionic currents and fluxes varied in the 67%-150% range was generated, with 787 models passing all the calibration criteria derived from human data (see Methods for details). To evoke EADs, we combined varying degrees of $I_{Kr}$ blockade with a concurrent smaller $I_{CaL}$ increase, recording at which point do EADs occur. In total, 782 out of 787 (99.4%) models in the population manifest EADs for a sufficient $I_{Kr}$

and $I_{CaL}$ perturbation (**Figure S19A**). The five remaining models have clearly anti-EAD parameters, having a small $I_{CaL}$ and $I_{NaCa}$, but large $I_{Kr}$, and would also show EADs if $I_{CaL}$ was increased further.

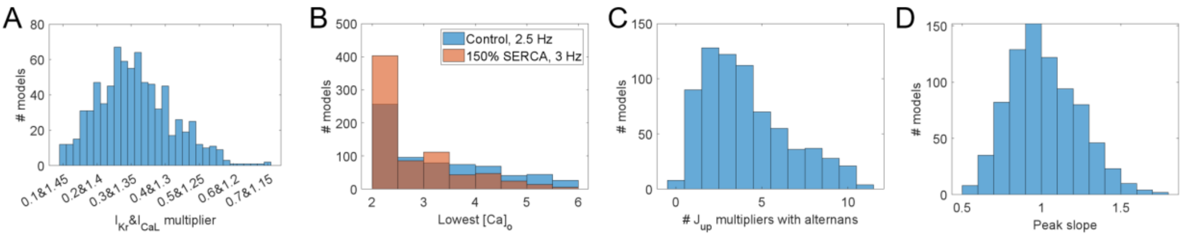

***Figure S19. Stability of arrhythmic behaviours in T-World. A)*** *Histogram describing the degree of pro-EAD alteration to the baseline cell (increasing $I_{CaL}$ and reducing $I_{Kr}$) that is sufficient to trigger EADs in a calibrated population of models (see Methods for details).* ***B)*** *A histogram of the level of extracellular Ca that is sufficient to trigger DADs following rapid pre-pacing in the presence of βARS stimulation. Both the baseline T-World and a version with 50% higher $J_{up}$ via SERCA pumps were simulated.* ***C)*** *A histogram of how many models among those with $J_{up}$ multipliers of 0.5, 0.6, ..., 1.5 (11 in total) manifest alternans at 260 ms basic cycle length.* ***D)*** *Distribution of peak S1-S2 slopes in the calibrated population.*

To evoke DADs and/or triggered activity resulting from spontaneous SR calcium release in baseline T-World (not including stochastic calcium release), we varied the extracellular calcium between 2 and 6 mM, recording the lowest level which produces spontaneous calcium release for each cell in the population. 686 of the 787 (87.2%) of control cells manifest DADs for a sufficiently high extracellular calcium (**Figure S19B**). This increases to 735 of 787 (93.4%) when increasing SERCA pump amount by 50% in the models to facilitate SR calcium overload. The key factor preventing DADs in a part of the population is increased $I_{NaCa}$ (**Figure S20**). This follows from the fact that with increased $I_{NaCa}$, the dyads clear released calcium more rapidly, which limits the pro-DAD condition of high dyadic calcium combined with high $[Ca]_{SR}$ load, as also observed previously [144].

To evoke alternans, we multiplied SERCA uptake by a range of multiplicators between 0.5 and 1.5 for each model in the population, counting how many corresponding models manifest alternans at 260 ms basic cycle length. 712 models in the population can capture all stimuli at such rapid pacing, developing two full action potentials with control $J_{up}$ level. 703 out of those 712 models (98.7%) manifest alternans for at least one tested $J_{up}$ multiplier (**Figure S19C**).

To assess S1-S2 restitution, we simulated all the models in the population with S1 interval = 1000 ms, recording the resulting slopes (**Figure S19D**). Under these conditions, 381 out of 787 (48.5%) models manifest restitution slope over 1 (mean across the population is 1.017). Such variability is generally consistent with human recordings, where the fraction of samples with slope > 1 was reported to be 44% [45], 74% [44], and 61% [40].

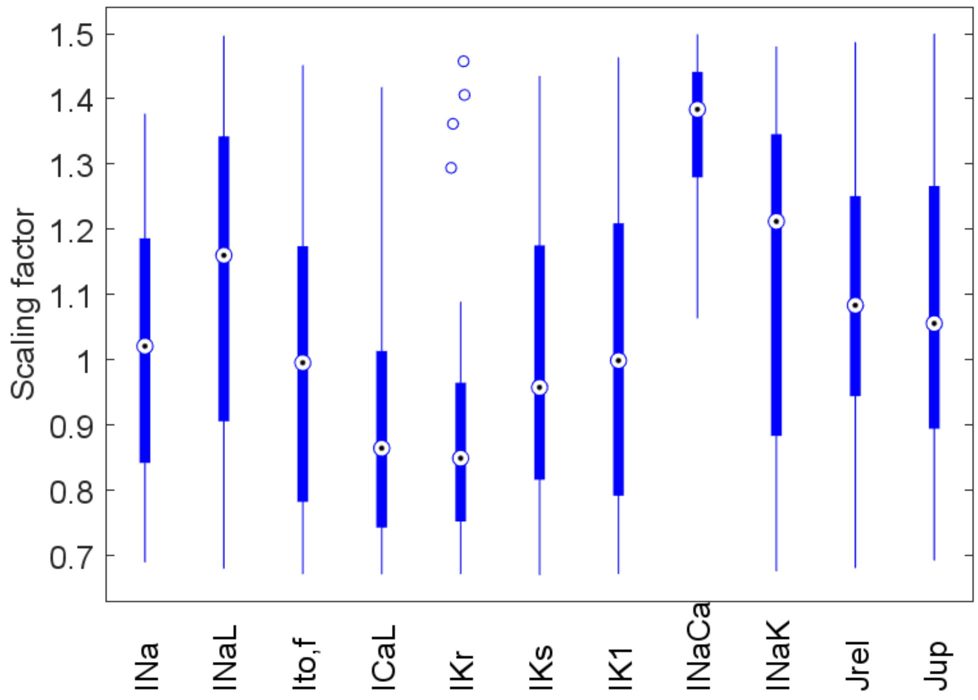

***Figure S20.*** *Boxplots of multiplier ranges in DAD-negative models in DAD stability assessment.*

#### Supplementary note 7: Disentangling anti-arrhythmic effect of mexiletine in LQTS2

We simulated T-World with emulated LQTS2, combining a 70% reduction in $I_{Kr}$ with an 82% increase in $I_{NaL}$, as reported by Crotti et al. [129]. To simulate mexiletine, we represented it either as described by Crumb et al. [128] ($MEX_{Crumb}$), used also in the drug safety assessment above, or as by Johannesen et al. [123] (see Methods for details). The two versions agree on mexiletine being primarily an $I_{NaL}$ blocker, partially also blocking $I_{CaL}$ and $I_{Kr}$, but differ in their effect on $I_{Ks}$. Both MEX versions induced APD shortening in the LQTS2 version of T-World, in line with clinical data (**Figure S21A**). The model also predicts reduction of EAD risk by both MEX versions, with $MEX_{Johannesen}$ (without $I_{Ks}$ inhibition) showing greater potency than $MEX_{Crumb}$ (with $I_{Ks}$ inhibition), despite less APD shortening at the given concentrations (**Figure S21B**). To understand the distinct effects of either mexiletine formulation, we compared the index of arrhythmic risk in the control model, with mexiletine, and for "single-knockouts", where each single effect of mexiletine on ionic currents is turned off. If turning off an effect increases the arrhythmic risk, it means that this effect is antiarrhythmic. In the case of $MEX_{Crumb}$, inhibition of $I_{NaL}$ and $I_{CaL}$ together drive the anti-arrhythmic effect, while the block of $I_{Kr}$ and $I_{Ks}$ increases arrhythmia vulnerability (**Figure S21C**). For $MEX_{Johannesen}$, the $I_{CaL}$ blockade and to a lesser extent $I_{NaL}$ block drive the anti-arrhythmic effect (**Figure S21D**). Given that there is no $I_{Ks}$ block, and only a marginal $I_{Kr}$ block at the given concentration, their effect on arrhythmic risk is none and small respectively. Together, these data suggest that $I_{Ca,L}$ inhibition plays a non-negligible role in the anti-arrhythmic effect of mexiletine and that further studies are needed to confirm or refute the effect of the drug on $I_{Ks}$. An $I_{Ks}$ blocking effect could be problematic particularly in the setting of high sympathetic drive, where this current is responsible for maintaining repolarization reserve and offsetting the $I_{CaL}$ increase.

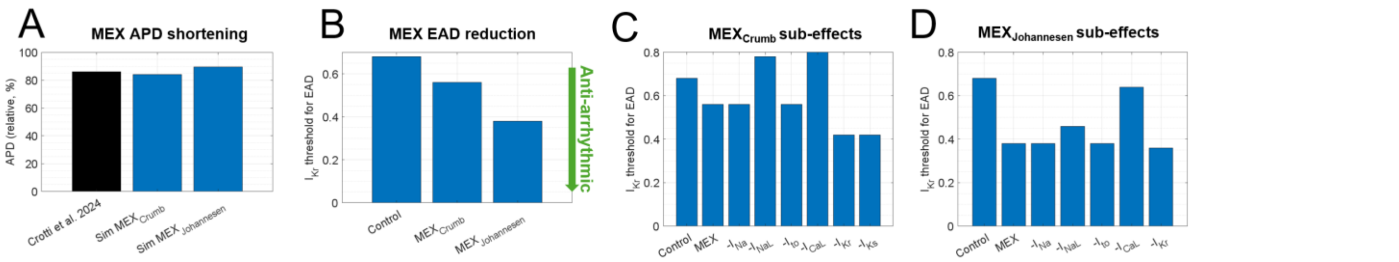

***Figure S21. Effects of mexilene on APD and EAD vulnerability. A)*** *APD shortening by two distinct simulated mexiletine formulations (see Methods), compared to data in* [129]. ***B)*** *A reduction in EAD risk for both mexiletine formulations (quantified using an EAD threshold: the largest* $I_{Kr}$ *multiplier sufficient to trigger EAD in a LQTS2 version of T-World).* ***C, D)*** *Comparison of EAD thresholds in the two mexiletine formulations, comparing an untreated LQTS2 cell, a cell treated with mexiletine, and then a range of mexiletine treatments where the effects of mexiletine on ionic currents were turned off one by one. An increase in those "knockout" bars compared to intact mexiletine indicate a protective effect of the given channel-blocking effect (omitting it increases EAD risk).*

#### Supplementary note 8: SERCA changes explain correlation between poor relaxation and alternans vulnerability in T2D

We hypothesized that the positive association between poor relaxation and greater alternans risk can be determined by the strength of SERCA pumps. As also investigated in our article, SERCA pumps largely determine alternans vulnerability, and they are the primary method of clearing calcium from the cytosol, thereby controlling relaxation. Comparing models from the first versus the fourth quartile in alternans vulnerability in the population used to produce **Figure 5E**, the $J_{up}$ flux carried by SERCA pumps is by far the clearest differentiating factor (**Figure S22A**). The same holds for the comparison between models with fast and slow relaxation (**Figure S22B**), confirming that changes in SERCA availability can explain the clinically observed connection between alternans and relaxation.

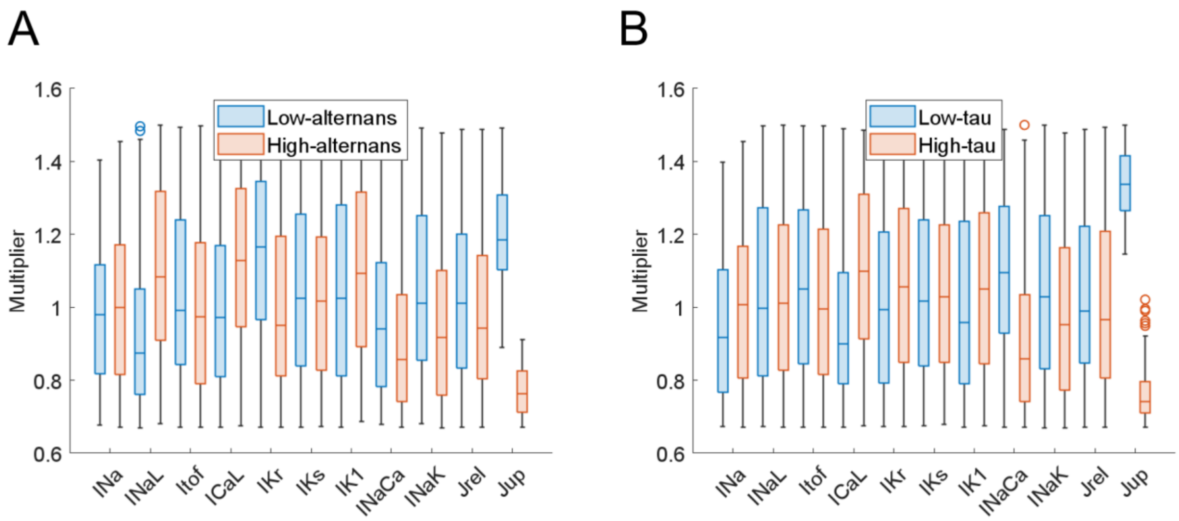

***Figure S22. Subpopulation differences in alternans and relaxation**. **A)** Distribution of parameter scaling constants in lower versus upper quartile with regards to alternans threshold in the T2D population of models. **B)** Similar comparison of distributions, comparing lower versus upper quartile with regards to the tau of relaxation.*

#### Supplementary note 9: Future NaV1.8 developments

More sophisticated models of cardiac $Na_V1.8$ current may be developed in the future, e.g. taking into account the fact that $Na_V1.8$ is modulated by CaMKII [145], and the channels may be thus more active at faster heart rates. In addition, a localization of $Na_V1.8$ near connexins in the heart was reported [146]. Beyond the possible role of those channels in conduction, their localized nature may potentially give rise to sufficiently high local elevation of sodium ions that would be translated into a local elevation of calcium via NCX. This could contribute to the formation of calcium waves, in the origin of which $Na_V1.8$ was also implicated [147].

#### Supplementary note 10: Components of T-World credibility

There are multiple properties of T-World that give confidence in its utility and trustworthiness. Most importantly, the model was extensively validated on unseen data (**Table S2**), correctly predicting the response to drugs, disease remodelling, sympathetic stimulation, sex differences, and modulation of arrhythmic risk by interventions with a known effect. Second, the advantage of biophysically detailed models is that they are mostly built from trusted components. Unlike machine learning models with billions of free parameters, T-World is built from components (ionic channels and signalling pathways), which were derived from experimental measurements, markedly reducing the risk of model overfitting and consequent incorrect predictions. Finally, an important component of trust and translational potential is the human nature of the model: it was built from human data wherever available, and it is calibrated to represent human physiology. There are important species differences between hearts of humans and animals, particularly the most popular animal models such as mice and rats[148]. This has crucial implications for applications such as arrhythmia research and/or studying the effect of drugs. For example, most episodes of drug-induced arrhythmia in human are due to the effect of drugs on hERG channel in human hearts[149] – but this channel is nearly absent in hearts of mice and rats. I.e., testing the safety of drugs in those species is of limited use, as those species lack the key component responsible for the issue.

#### Supplementary note 11: Possible omitted effects of drugs in in silico safety assessment

Here we discuss several drugs that were labelled as Safe by T-World, whereas in reality they have Possible or Conditional risk based on Crediblemeds[150]. For example, clozapine is known to sometimes cause myocarditis and cardiomyopathy[151], and may lead to sympathetic hyperactivity[152], neither of which is captured by the channel blocking description of the drug. Voriconazole appears to interact with metabolism of other, possibly riskier drugs[153], which could explain its pro-arrhythmic effect. The data on action of voriconazole used in our study indicate it blocks $I_{CaL}$ and $I_{Kr}$ (the former slightly more), which would be a generally safe profile, if these were the sole effects. Dasatinib is a tyrosine kinase inhibitor used in cancer therapy, which targets multiple signalling pathways and may promote heart failure in some subjects[154], so it is also possible it acts beyond QT prolongation.

#### Supplementary note 12: Limitations

The most general limitation of computer models such as T-World is that they may faithfully represent only phenomena emerging from the model components included. However, this limitation can be in fact leveraged as a research tool. A discrepancy between a computer model and reality may help researchers identify where our understanding of biology (formalized in the model) is lacking, and what may explain the discrepancy. This may be, e.g., the need to include an additional signalling pathway, demonstrating its likely involvement in the phenomenon of interest.

The second known limitation of models of this level of abstraction is the limited capability to fully capture DADs. A DAD typically arises as a spontaneous calcium release in a small portion of cell, which then diffusively spreads through the cell, recruiting other RyRs, leading to a large-scale calcium release. The aspect of diffusion is not captured by the nature of T-World and similar models, which represent an average calcium concentration in each compartment.

Other models with spatially distributed calcium handling may be used to study calcium wave emergence and spreading in myocytes[155–157], but they are far more computationally demanding, which limits scalability to whole heart level simulations, and they are often less broadly validated beyond the primary scope of use.

Our model does not reproduce the known phenomenon of extracellular calcium shortening APD (not shown), a limitation shared by most cardiomyocyte models, with mechanism unknown. If the role of extracellular calcium is a central focus of a study, the BPS2020 model[158] may be used, which in addition also recapitulates several arrhythmic behaviours. That said, the action potential shape is not particularly human-like, and the reported alternans is not a typical calcium-driven alternans, but results from calcium transient developing only in every other beat, which is then naturally translated into APD oscillation. Additionally, as the model relies on strong calcium-dependent inactivation of $I_{CaL}$, the resulting "alternans" exhibits electromechanical discordance (longer APD in CaT-absent beats). This contradicts most data in human-like species [24,30,31], and while $I_{CaL}$ inactivation likely contributes to extracellular calcium responses, further research is needed to confirm its central role.

### Supplementary figures

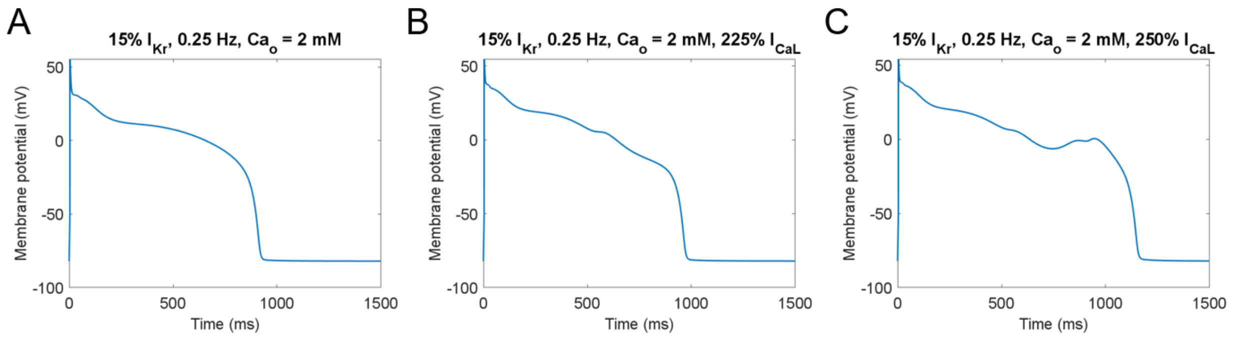

***Figure S23*****. *****EADs in the Morotti 2021 model.*** ***A****) Standard conditions used to evoke EADs as in [27]; 0.25 Hz pacing, 15%* $I_{Kr}$ *availability, and extracellular calcium of 2 mM. No EADs are evoked.* ***B)*** *Demonstration that even with a substantial increase in* $I_{CaL}$ *(225% of control, i.e., 115% increase) is still not sufficient to trigger EADs.* ***C)*** *With 250%* $I_{CaL}$*, an EAD is evoked. However, the rather wobbly AP morphology is not particularly similar to experimental measurements.*

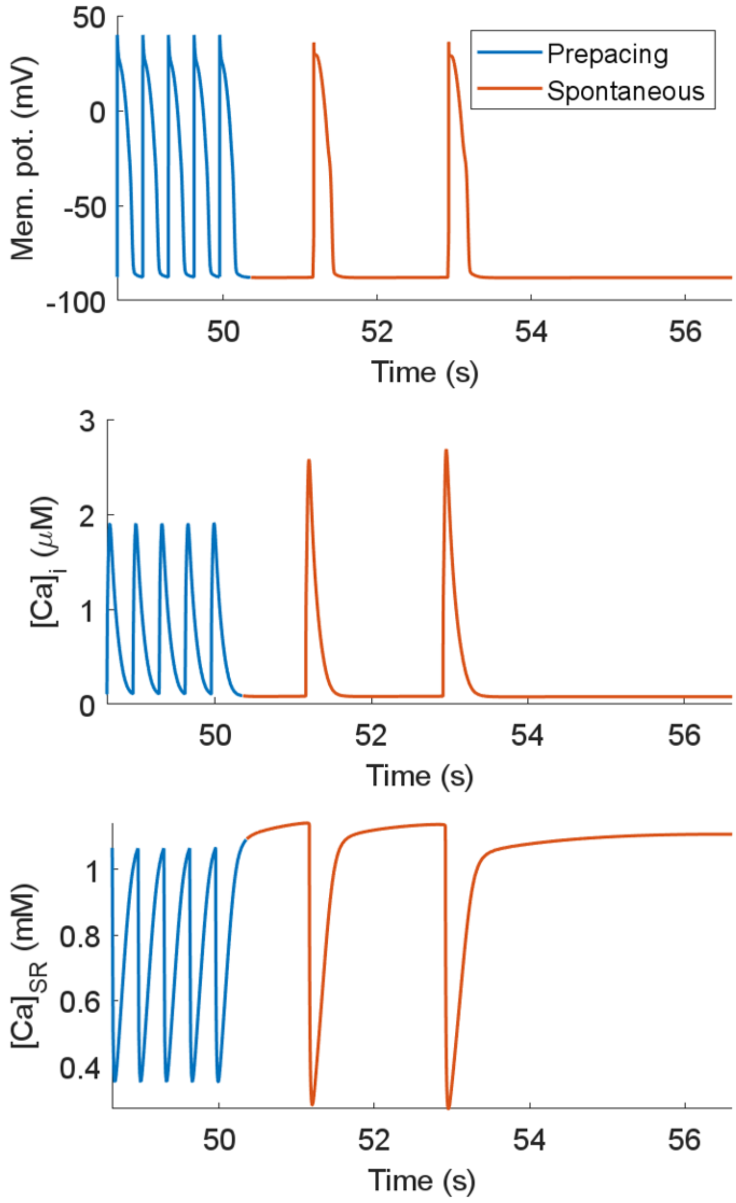

***Figure S24. Multiple consecutive DADs in T-World.*** *This was produced by pacing the model in similar conditions as in Figure 4D (extracellular calcium of 4.5, 150 beats of pre-pacing), except 3 Hz, rather than 2.5 Hz pre-pacing rate was used.*

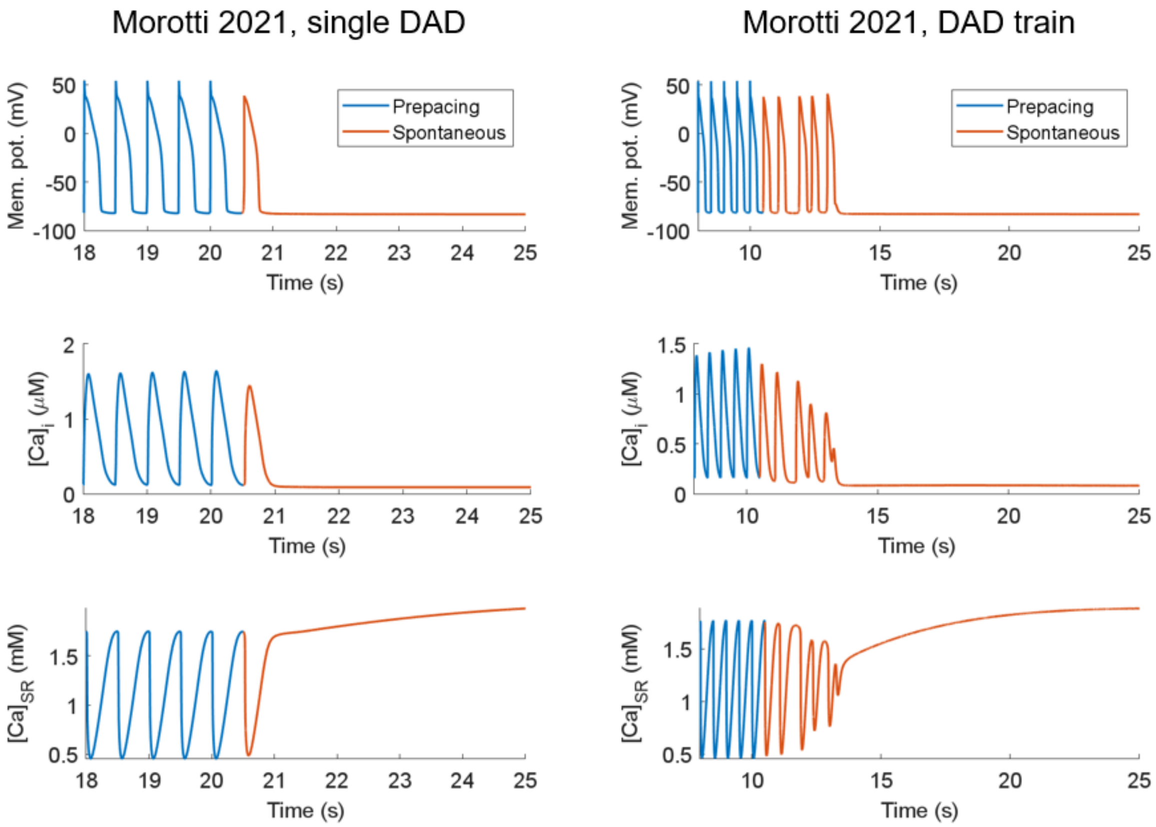

***Figure S25. DADs in the Morotti2021 model****. In the* ***left*** *column are the membrane potential, intracellular calcium, and SR calcium. The cell model was pre-paced for 40 beats at 2 Hz with 0.1 μM simulated isoproterenol and extracellular calcium of 3.5 mM, following which a single beat followed by long period of quiescence was applied. Similarly to Figure 4D of the main manuscript, blue traces indicate the paced beats, whereas the red ones arise from spontaneous calcium release. In the right panel is an example of a multi-DAD train, achieved using a similar protocol, which occurred at 4 mM extracellular calcium, and the pre-pacing was applied for 20 beats. In either case, endocardial model was used, with parameters myoFlag = 1 and mechFlag = 0.*

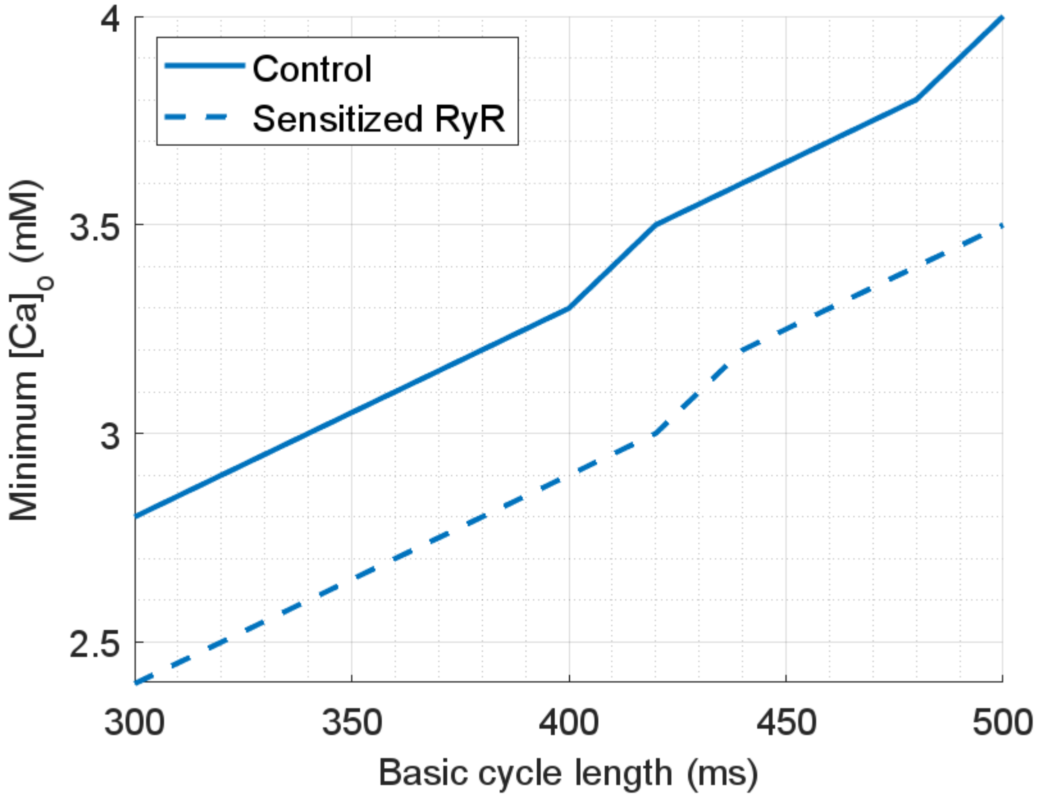

***Figure S26. DAD validation.*** *To validate the DAD generation in T-World, we confirmed that pre-pacing at higher frequencies facilitates DADs, as observed experimentally[159]. We also simulated a version of the model with sensitized RyR (doubled RyR opening rate), which also promoted DAD formation in the presence of β-AR stimulation, consistent with experimental findings [68]. The figure shows the DAD threshold (expressed as the minimum extracellular calcium concentration needed to evoke DADs) versus pre-pacing frequency. In solid line is shown the control model, with a sensitized model (doubled transition rate from closed to open RyR state) shown by dashed line. The lower threshold for shorter basic cycle length, and for the sensitized RyR indicates increased DAD vulnerability.*

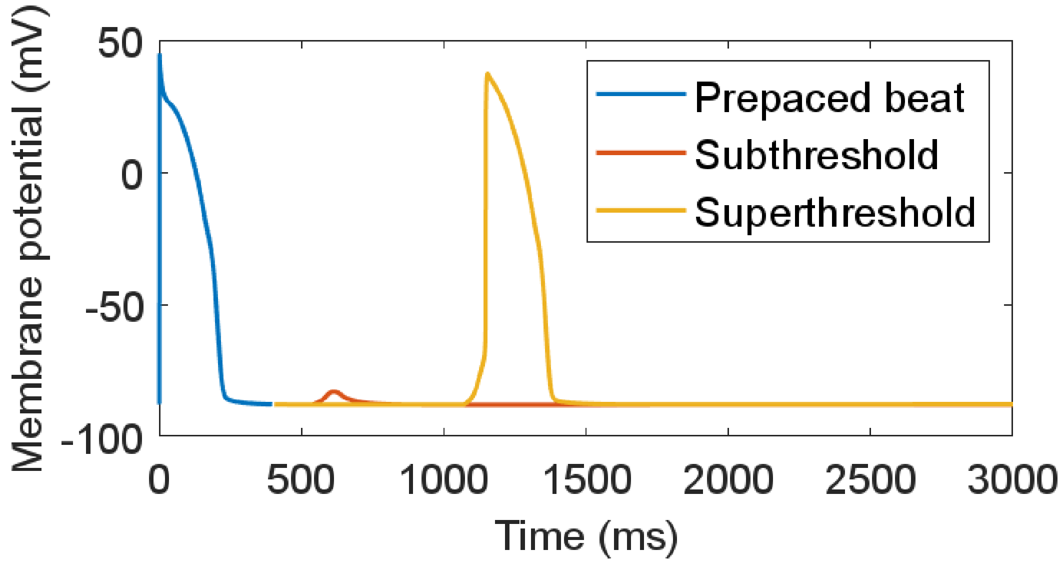

***Figure S27. Examples of DADs generated by stochastic spontaneous calcium release.*** *See Supplementary Methods-Arrhythmia Studies for details of implementation.*

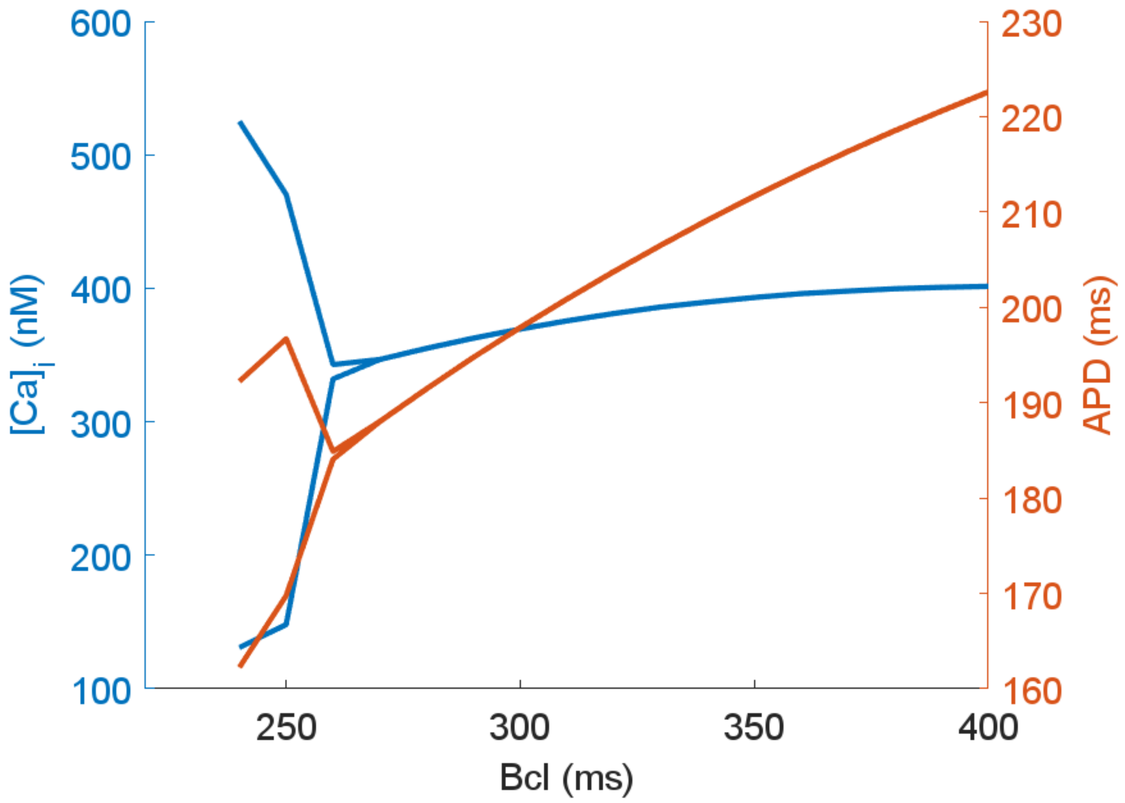

***Figure S28. Rate-dependent occurrence of CaT and APD alternans in T-World.***

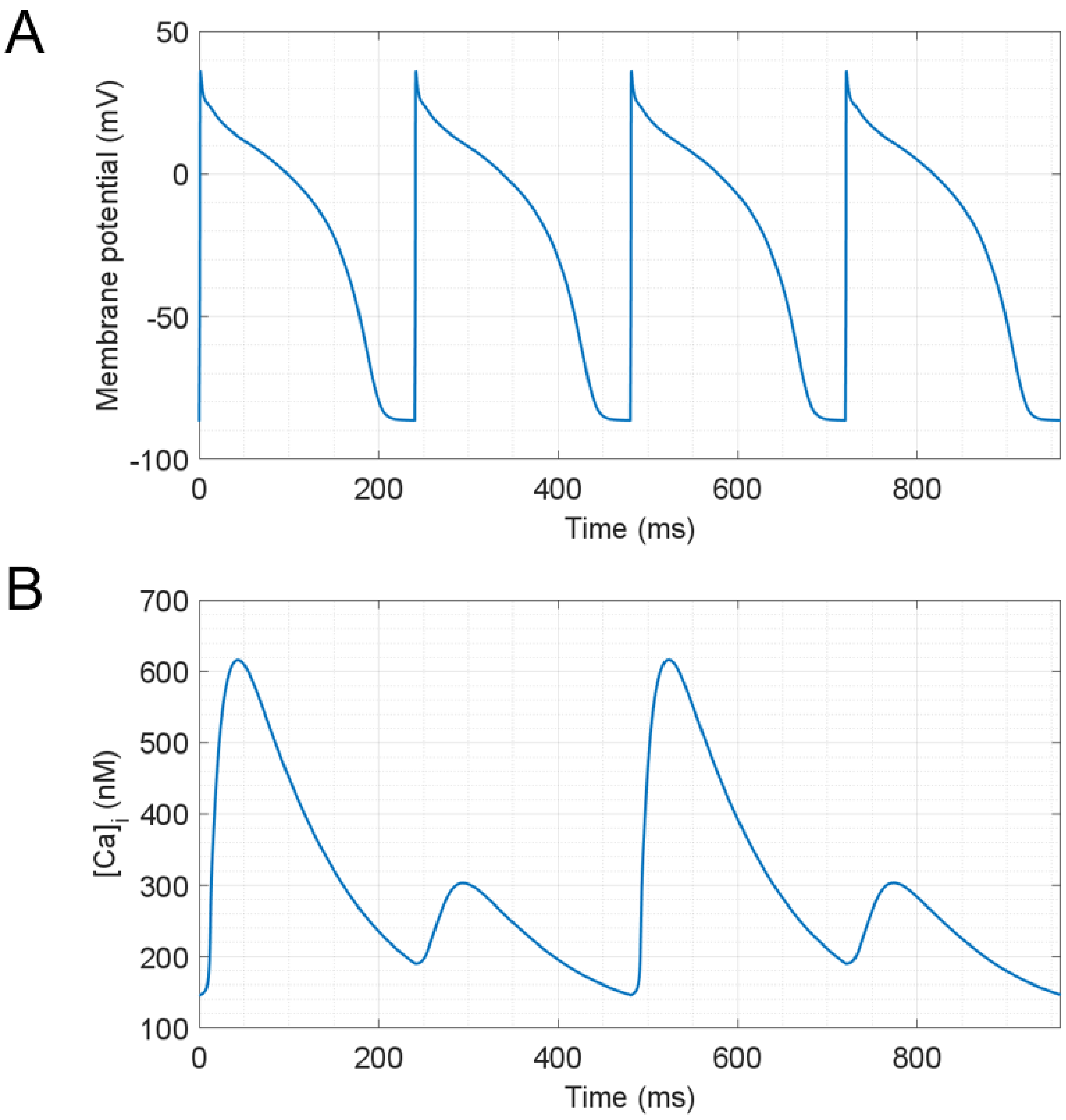

***Figure S29. Calcium alternans in T-World under AP clamp.*** *Here we used a fixed-shape action potential shape applied 250 times to T-World with a basic cycle length of 240 ms. A) Four identical consecutive action potentials. B) The last four calcium transients out of the 250 evoked, showing clear calcium alternans.*

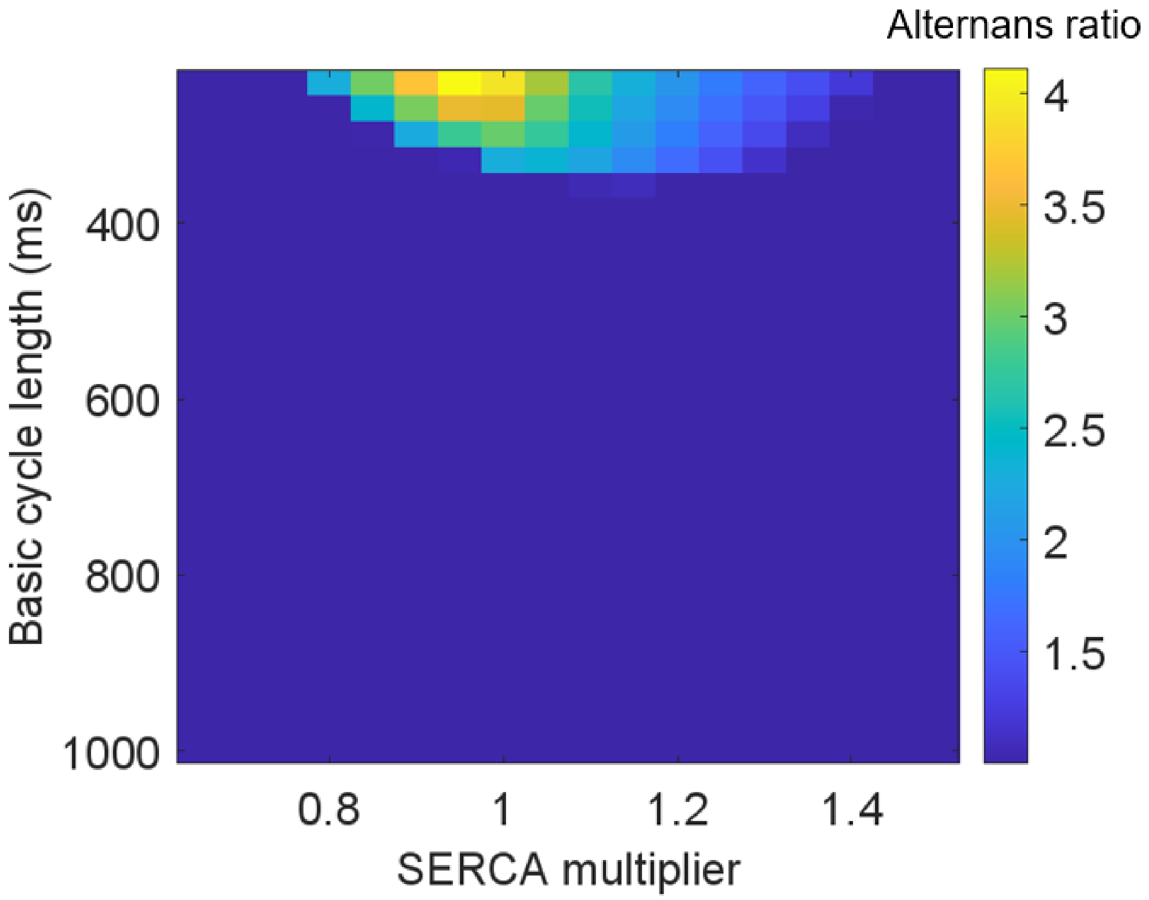

***Figure S30. Visualization of the response of alternans in ToR-ORd to changes in SERCA pumps**. The color codes calcium alternans ratio (amplitude of the larger CaT divided by the amplitude of the smaller CaT among the last two beats out of 250). On the x-axis is the multiplier of $J_{up}$ (1 corresponds to control value), and on the y-axis is the basic cycle length used to pace the cell. It can be clearly seen that reducing SERCA availability does not lead to extension of alternans to slower frequencies; rather, it attenuates or abolishes alternans, in disagreement with experimental data. We believe this is related to a less realistic alternans mechanism in ToR-ORd, which relies on slow refilling of junctional SR from the network SR (the model uses two-compartment model of SR, unlike T-World, which uses single compartment). Available experimental data indicate the refilling to be relatively fast, faster than needed by the model to achieve alternans [160]. In addition, the qualitatively correct behaviour of a large SERCA increase in ToR-ORd inhibiting alternans is also mechanistically problematic: it relies on full depletion of SR during a release, which trivially inhibits alternans, and that we described previously [161]. Such a depletion appears to be unrealistic: normal fractional release was reported to be 35%, increasing to 60% when a strong release trigger was used, but definitely not approaching full depletion [162].*

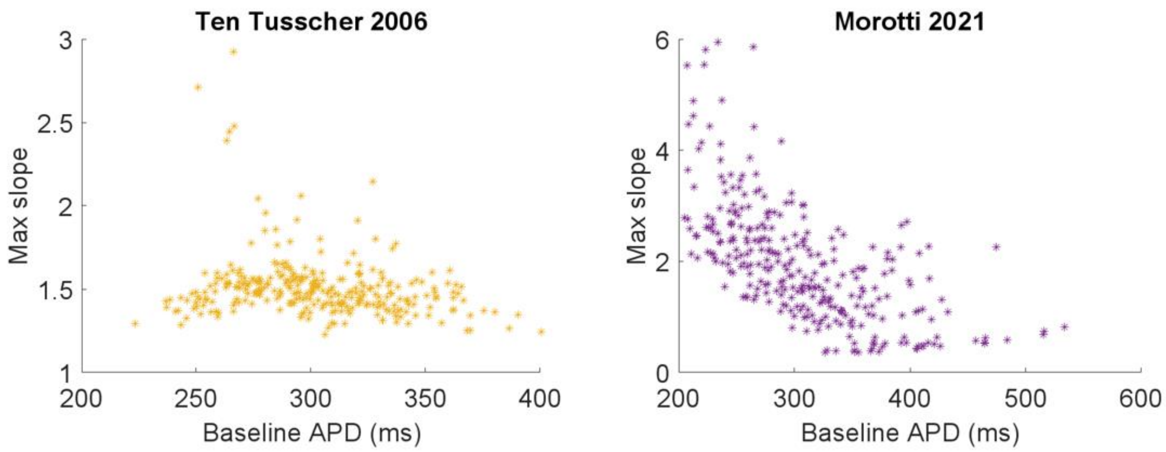

***Figure S31.** APD-Slope restitution in TP06 (left) and Morotti 2021 (right) models. The simulations were carried out using the same protocol as T-World in **Figure 3D** of the main manuscript.*

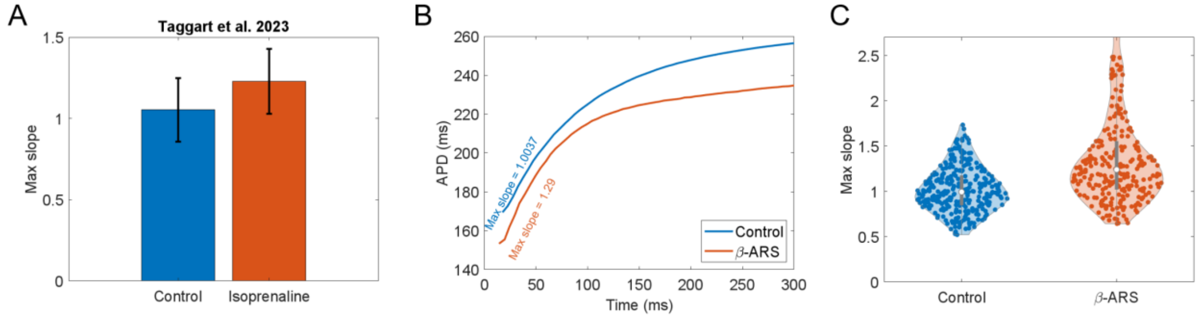

***Figure S32. The effect of βARS on restitution of T-World. A)*** *Human data demonstrating steepening of S1-S2 restitution with βARS activation [43]. **B)** Corresponding simulation of the impact of βARS activation on restitution in T-World. **C)** Simulation of a group of models with randomly perturbed parameters, showing overall restitution steepening with βARS.*

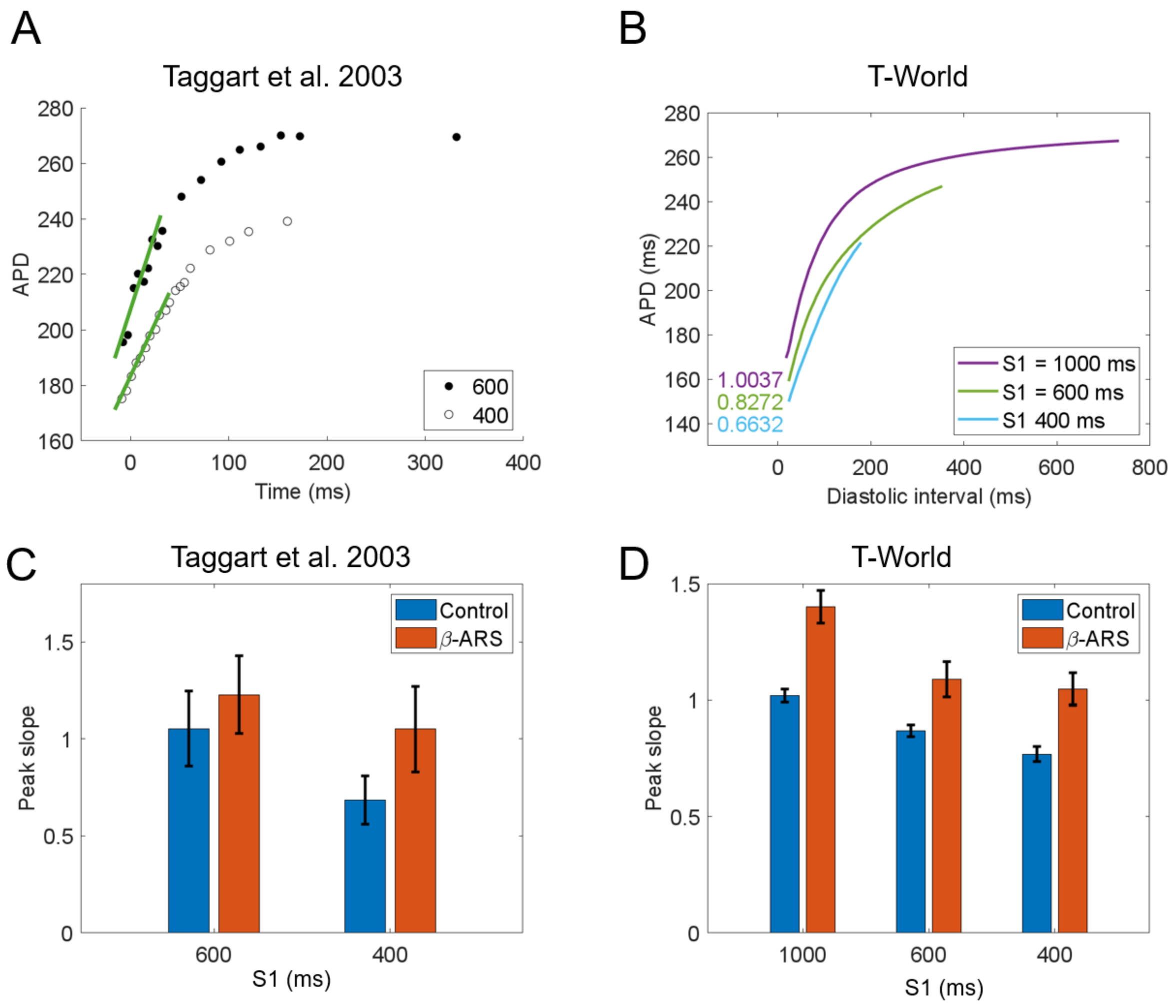

***Figure S33. Modulation of S1S2 restitution slope by pre-pacing rate. A**) Examples of restitution curves with S1=600 and 400 ms from the study by Taggart et al. [43]. **B)** Corresponding simulations using T-World. **C)** Quantitative summary of the peak slope at two S1 basic cycle lengths, with/without β-AR stimulation, based again on Taggart et al. [43]. **D)** Corresponding simulations summarizing 300 T-World models with varied parameters, as described in Methods.*

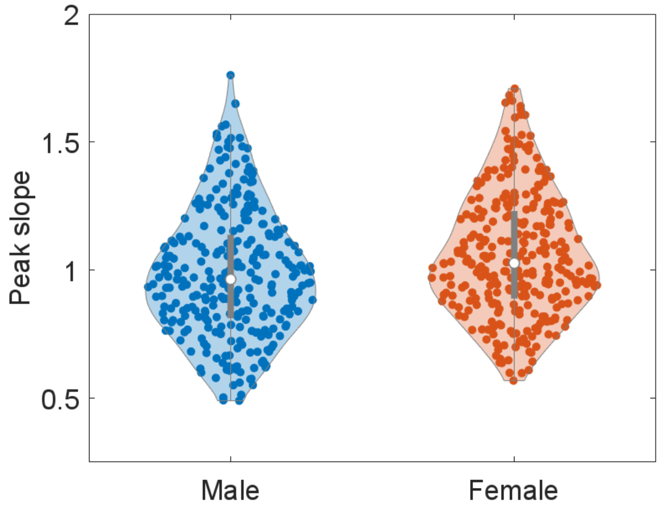

***Figure S34. Comparison of peak S1S2 slope in a population of male and female models.*** *On average, the maximum slope increases by 0.092 when a myocyte is switched from male to female, with 85.3% models showing an increase.*

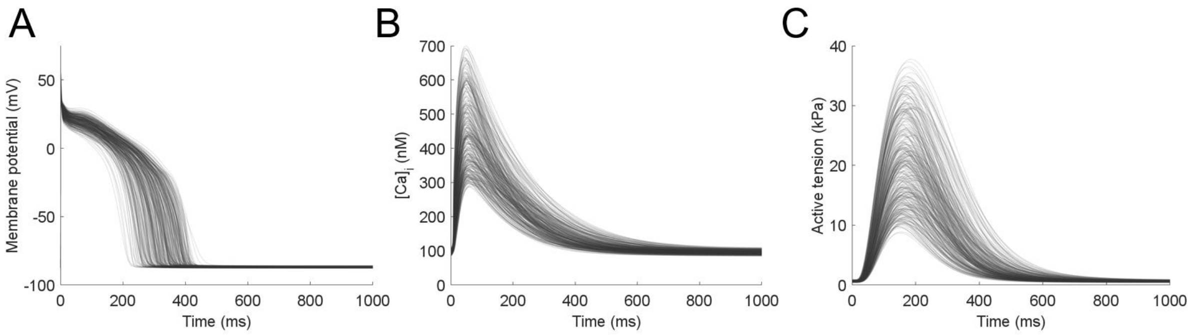

***Figure S35. Population of 337 models used in the in silico trial. A)*** *Action potentials,* ***B)*** *Calcium transients,* ***C)*** *Active tension.*

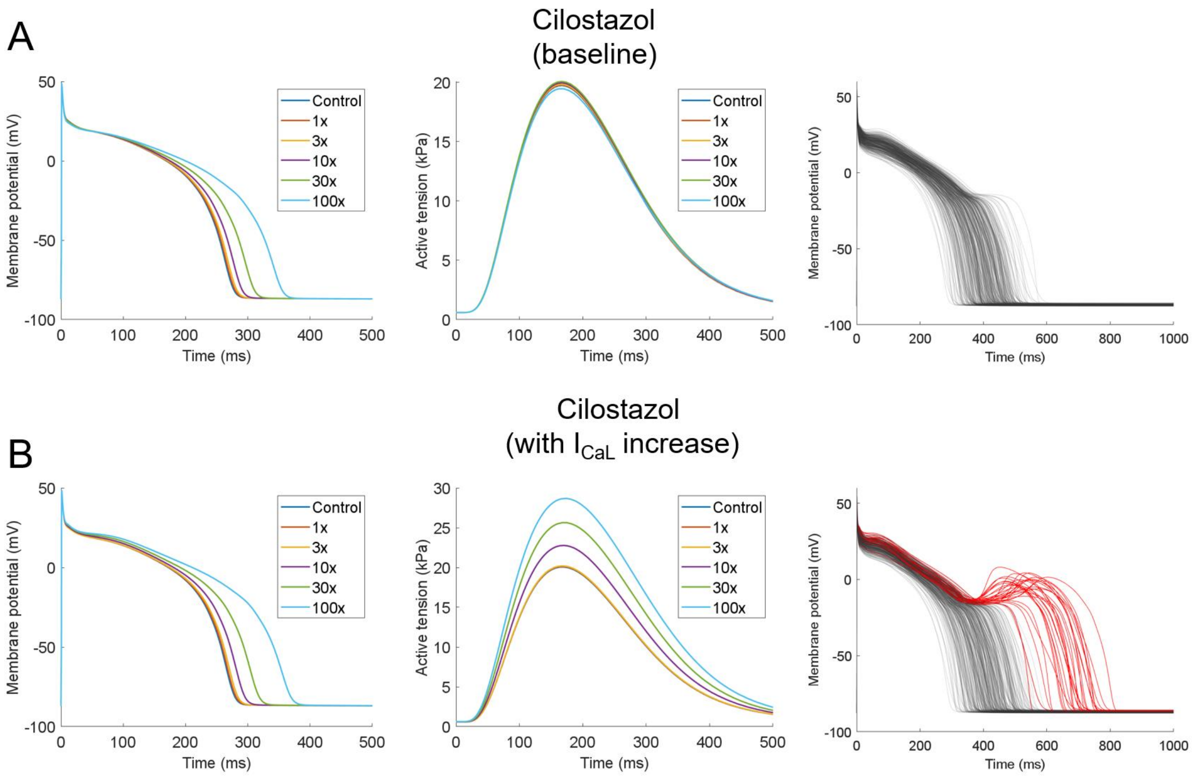

***Figure S36. Identifying the missing effect of cilostazol in original pharmacological data.*** *Cilostazol is known to be a relatively high-risk PDE3 inhibitor drug, which is positively inotropic. However, the original formulation of cilostazol as described by Kramer et al.* [163]*, used historically in in silico trials, only describes the drug a preferentially $I_{Kr}$ blocker (IC50 13.8 μM) with a weaker $I_{CaL}$ block (IC50 91.2 μM). Simulating such a drug formulation leads to dose-dependent QT prolongation, but virtually no change in predicted contractility (**panel A**, left and centre). This is at odds with data indicating a substantial increase in contractility following cilostazol exposure* [127]*. However, when we incorporated the PDE3 inhibitory effect leading to $I_{CaL}$ increase through PKA signalling based on* [127] *(+10% at 10x dose = 1.28 μM, +22% at 30x dose, and +40% at 100x), a pro-contractile effect has clearly emerged (**panel B**, centre). At the highest dose of 100x corresponding to 12.8 μM, peak active tension increased by 43% versus untreated cells, which is entirely comparable to the experimental data in guinea pig that observed a 35% increase in peak contractile tension with 10 μM cilostazol. The increased $I_{CaL}$ stemming from PDE3 inhibition leads to a substantially greater arrhythmic risk, triggering EADs in many more models in the population compared to the baseline formulation (**panel B**, right, versus **panel A**, right). It is noteworthy that the APD prolongation is only marginally greater in the model reflecting PDE3-inhibition-based $I_{CaL}$ increase compared to the baseline formulation, and APD effect thus cannot be used to identify a likely problem in the baseline drug formulation. This highlights how multiple phenotypic aspects need to be considered when trying to detect gaps in the pharmacological descriptions of drugs.*

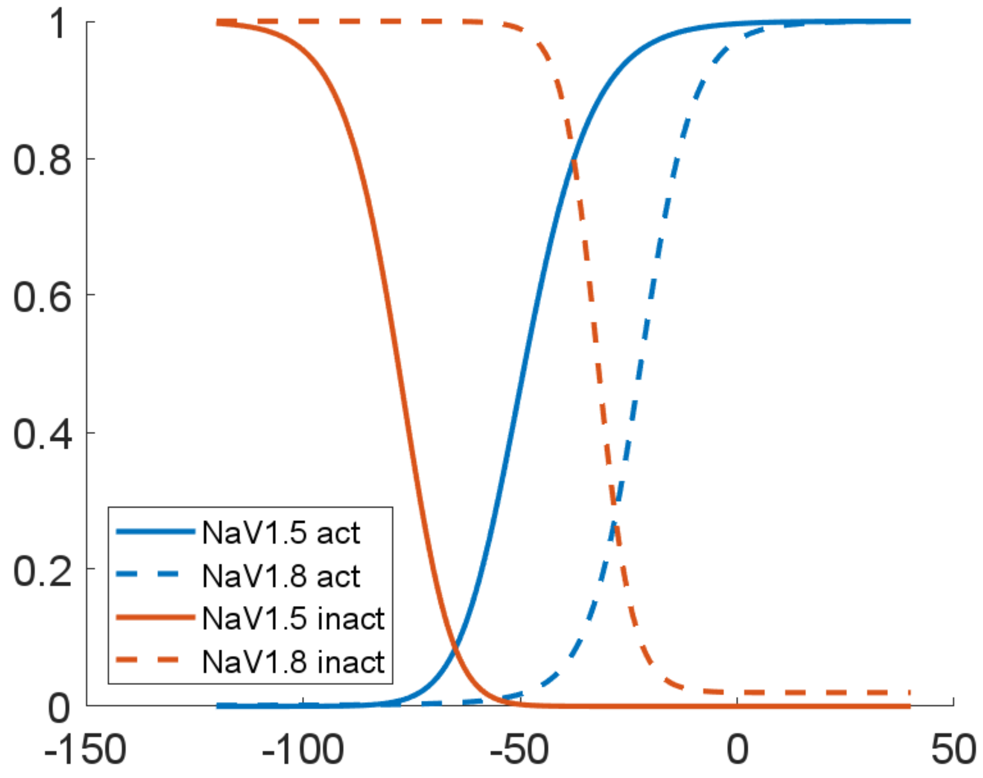

***Figure S37. Steady-state activation and inactivation curves of sodium currents in T-World.*** *Shown are the activation and inactivation of the standard cardiac $I_{Na}$ current through NaV1.5, versus the activation and inactivation of current carried by the predominantly neuronal NaV1.8 (see Methods for details), which are known to be right-shifted [164,165]. Please note that only the NaV1.5 current is used in T-World by default, with NaV1.8 an optional addition relevant in modelling, e.g., the heart failure.*

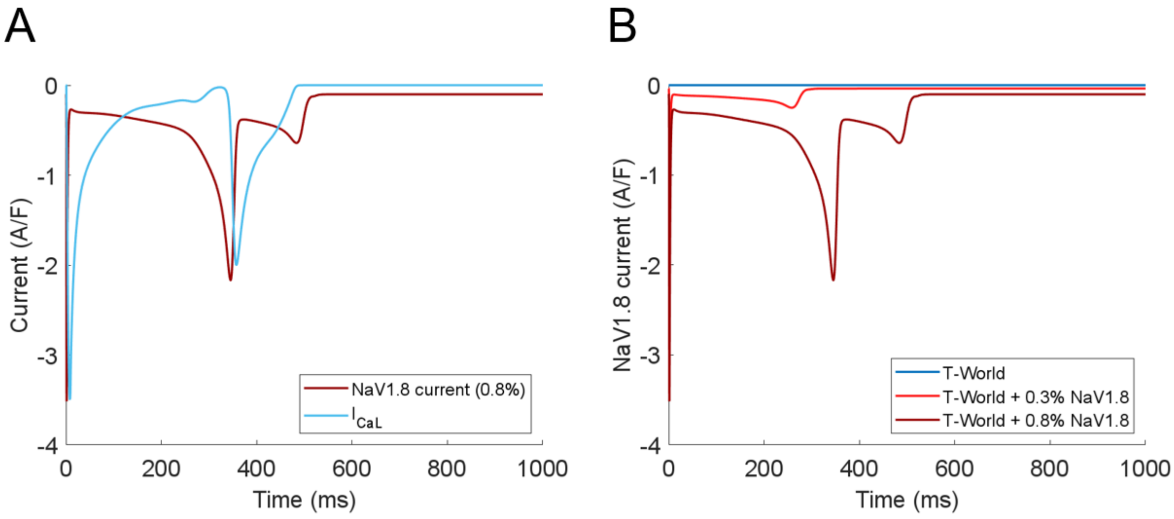

***Figure S38. NaV1.8 activation during EAD. A)*** *A plot of NaV1.8 current and $I_{CaL}$ in the model with the relatively highest NaV1.8 current density, showing that NaV1.8 current activates first, driving the early depolarization during the first phase of the EAD generated. This subsequently activates $I_{CaL}$.* ***B)*** *This panel illustrates that for 0.3% NaV1.8, the NaV1.8 current has a profile generally similar to standard late sodium current carried by cardiac NaV1.5. Only when the NaV1.8 current density is increased, it triggers cell depolarization in a manner distinct from NaV1.5 current.*
